# Abstract representation in the hippocampus predicts spontaneously adopted structure-based behavioral strategy

**DOI:** 10.64898/2026.09.20.752934

**Authors:** Yuhan Jiang, Yaorui Liu, Xinyu Zhao

## Abstract

Animals can solve the same problem using distinct strategies, ranging from memorization of specific sensory details to inference based on abstract task structure. Although the hippocampus has been implicated in cognitive map formation, how the structural information emerges in the hippocampus through learning and whether it is associated with behavioral strategies remain poorly understood. Here, we developed a structured cue-guided navigation task in virtual reality and performed longitudinal two-photon calcium imaging of hippocampal CA1 neurons throughout learning in mice. Although the task did not explicitly require learning of latent structure, most animals progressively transformed hippocampal map from sensory-oriented representations into cue-invariant representations of reward sequences. Strikingly, the emergence of such abstract hippocampal maps predicted whether individual animals adopted structure-based behavioral strategies when sensory information became partially unavailable. At the population level, reward-sequence representations formed a latent-state map capable for predicting future state transitions, generating a parsimonious Markov graph. Computational modeling further identified next-state prediction and representational regularization as two computational principles essential for generating similar latent-state structures across distinct architectures. Together, our findings suggest that the hippocampus spontaneously extracts abstract task structure through learning and that the geometry of hippocampal representations predicts problem-solving strategies.

## Introduction

One of the brain’s fundamental functions is to generate flexible behavioral strategies that enable animals to solve complex problems in ever-changing environments. In many situations, multiple strategies can be used to solve the same problem. Some strategies rely on memorizing specific sensory details, whereas others exploit sensory-invariant structures embedded within a task. Although both approaches may produce equally successful behavior in stable and fully observable environments, they can lead to markedly different outcomes when sensory cues become altered or unavailable. For example, by leveraging knowledge of spatial relationships among objects, an animal can precisely determine its location even when some local landmarks are modified or missing. Such structure-based reasoning is not only critical for navigation in natural environments, but may also provide a foundation for higher-order forms of human intelligence, including language, symbolic reasoning, and scientific inquiry.

Structure-based strategy requires the brain to represent generalizable task structures that are invariant to specific sensory details. Such representations are often referred to as ‘schemas’ in neuroscience^1^. Previous studies on schema representations have mostly focused on frontal cortical regions, including the orbitofrontal cortex (OFC) and medial prefrontal cortex (mPFC). In an olfactory go/no-go task containing multiple hidden odor sequences, ensembles in the rat OFC gradually developed low-dimensional representations that encoded positions within sequences independent of odor identity^2^. Similarly, in a periodic navigation task in which mice repeatedly traversed multiple goal locations in cycles, mPFC activity constitutes a ‘structure memory buffer’ that generalized behavioral sequences across different combinations of goals^3^. Furthermore, mPFC activity has been reported to represent high-level categorical information in both mice^4^ and macaques^5,6^. Beyond frontal cortical regions, the hippocampus has also been implicated in schema learning. Bilateral hippocampal lesions impair experience-dependent acceleration of learning in rats performing flavor-location association tasks with novel sensory cues^7^, suggesting a role for the hippocampus in acquiring abstract task rules.

Despite behavioral evidence linking the hippocampus to schema learning, how neural activities in the hippocampus represent abstract task structure remains controversial. In particular, there exists an apparent dichotomy between studies in human/non-human primates and those in rodents regarding the extent to which hippocampal representations are ‘abstract’. Studies in humans and non-human primates have long suggested that the hippocampus can represent abstract concepts invariant to specific sensory stimulations. For example, neurons in the human hippocampus exhibit image-invariant responses to celebrity identities^8^. Human hippocampal activity has also been shown to encode transition probabilities among visual stimuli in temporal sequences^9,10^. Such representations of state transition may support internally generated hippocampal sequences during model-based planning^11,12^. Moreover, abstract hippocampal representations have been linked to inferential reasoning in both humans^13^ and monkeys^14^. In contrast, classical physiological studies in rodents, particularly those focusing on place cells, have emphasized the sensory specificity of hippocampal representations. In spatial tasks, hippocampal place fields often undergo global remapping across environments with little preservation of structural organization, whereas generalized representations appear more prominently in regions such as the medial entorhinal cortex (mEC) and mPFC^15–17^. Nevertheless, recent studies increasingly suggest that the rodent hippocampus also encodes non-spatial task variables, including positions within sound sequences^18^, accumulated evidence^19^, and reward-contingency contexts^20,21^. Both theoretical^22–24^ and experimental^25–27^ works further support the view that the hippocampus constructs a predictive cognitive map that encodes task states and their relational structure. If such predictive maps can become invariant to alterations of sensory inputs—that is, become abstract—they may support structure-based decision making. However, we are yet to fully understand how hippocampal representations evolve during learning to acquire abstract task structures and whether such representations are associated with an animal’s structure-based behavioral strategy.

To address these questions, we developed a behavioral paradigm in which mice collected rewards in a dynamic environment and subsequently tested whether animals employed structure-based strategies by selectively masking sensory cues. We performed longitudinal two-photon calcium imaging in the hippocampus throughout learning to track the evolution of neural dynamics. Although our task did not explicitly require animals to learn the underlying task structure, most mice exhibited a transformation of hippocampal representations from sensory-oriented coding to abstract coding of reward sequences. Detailed characterization further revealed that this abstract hippocampal map manifests a parsimonious state graph of reward progression. Strikingly, the emergence of abstract hippocampal map predicted that animals spontaneously adopted a structure-based strategy, as revealed in probe trials in which sensory cues were unavailable at the last position of a sequence. Finally, through computational modeling using diverse network architectures and learning algorithms, we identified next-state prediction and strong regularization as two key computational principles necessary for learning such representations. Together, our findings suggest that abstraction of hippocampal representations may support structure-based decision making and thus potentially provides a neural mechanism for flexible behavior in dynamic environments.

## Results

### Structured cue-guided navigation

We developed a structured cue-guided navigation task for head-fixed mice navigating virtual reality (VR) environments (Fig. 1A). Mice ran through a 238-cm linear corridor containing sequences of visual patterns and were teleported back to the start position upon reaching the end of the track, followed by a 2-4 s inter-trial interval (black screen). Three distinguishable visual cues (A, B, and C) appeared at three fixed locations separated by 60 cm. Cue A and cue C were associated with reward delivery (10% sucrose water), whereas cue B was unrewarded (Fig. 1B, middle). Mice were trained to lick an empty lick port at rewarded cue locations to trigger reward delivery.

**Figure 1.**
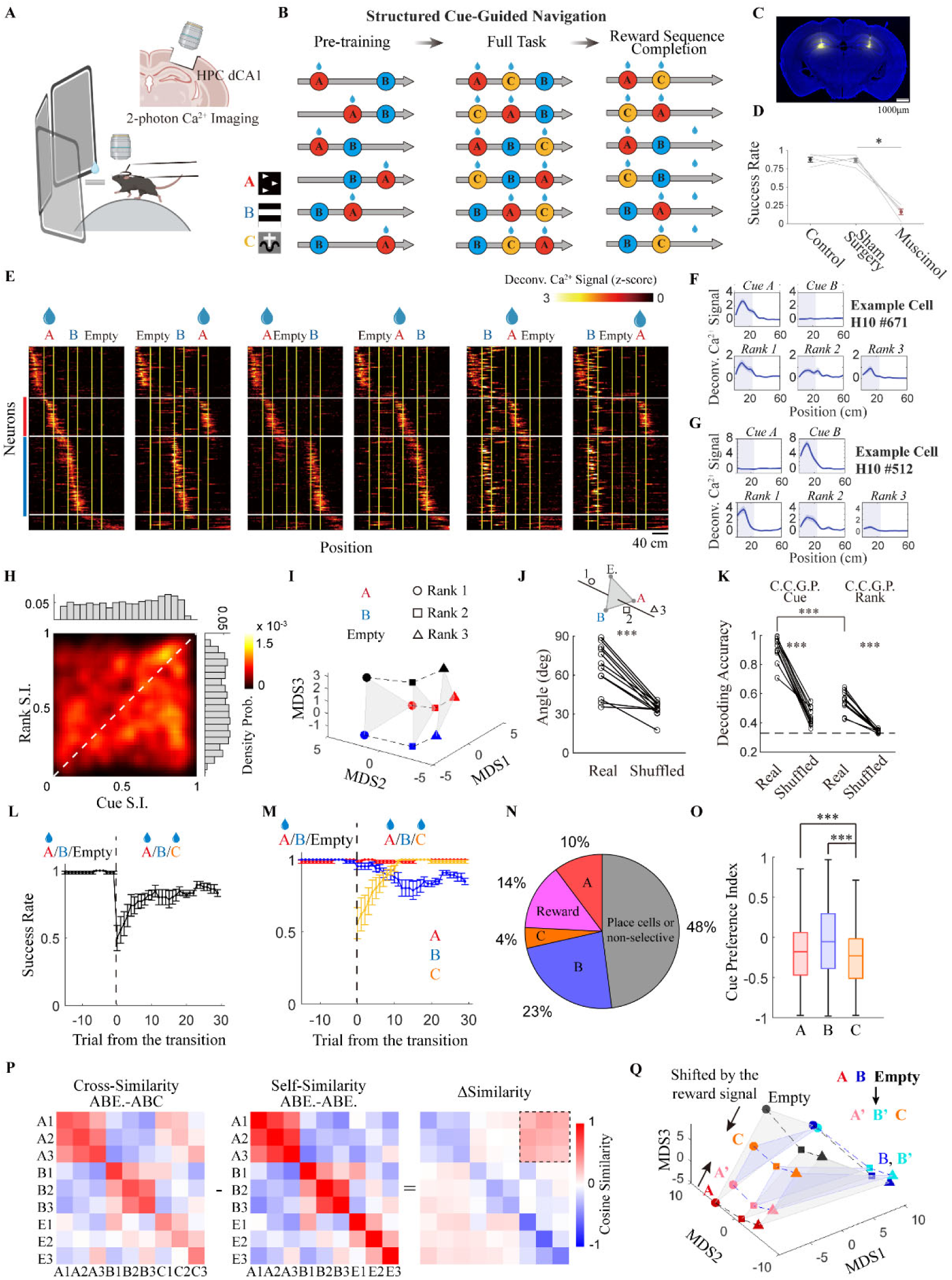
Disentangled representation of cue and rank in structured cue-guided navigation. **(A)** Experimental set-up. Head-fixed mice ran on a floating spherical treadmill surrounded by virtual reality (VR) screens. A lick port positioned near the mouth delivered sucrose reward during the task. Simultaneous two-photon calcium imaging was performed throughout behavioral training. Upper right, schematic illustration of the imaging area in the dorsal CA1 subregion of the hippocampus. **(B)** Structured cue-guided navigation task: pre-training (A/B/Empty sequences), full task (A/B/C sequences), and reward sequence completion. Bottom left: Actual patterns of cue A, B and C. **(C)** Histological verification of bilateral hippocampal silencing. Blue: DAPI staining; yellow: BODIPY TMR-X muscimol conjugate. **(D)** Success rates under control, sham surgery and muscimol conditions. *P* = 0.03125 for sham control versus Muscimol group; one-sided Wilcoxon signed-rank test; n = 5 mice. **(E)** z-scored neural activities in one example mouse. Cells were sorted based on their activity peaks in the A/B/Empty trial type (the first column). Yellow lines indicate possible cue positions. Red and blue bars marked cells anchored with cue A and cue B, respectively. **(F)-(G)** Activities across different cues and ranks of example neurons. Solid lines and shaded areas represent mean activity and s.e.m., respectively. Gray rectangles denote cue locations. **(H)** 2-D density map of cue selectivity indices and rank selectivity indices for highly responsive cells pooled across all mice during the expert stage after pre-training (n = 12 mice). **(I)** Organization of neural representations across different combinations of cue identity and sequence rank. High-dimensional population vectors were projected into a three-dimensional space using multi-dimensional scaling (MDS). Red, blue, and black denote cue A, cue B, and the empty slot, respectively. Circles, squares, and triangles denote rank 1, rank 2, and rank 3, respectively. Gray triangular planes and dashed lines indicate cue-coding planes and rank-coding axes, respectively. Cells from all mice were pooled together. **(J)** Angle between the cue-coding plane and rank-coding axis. For each mouse, the cue-coding plane and rank-coding axis were calculated as the averages of the three planes and three axes shown in panel (I), respectively. *P* = 0.0009766, real versus shuffled cell identity; two-sided Wilcoxon signed-rank test; n = 12 mice. **(K)** Cross-condition generalization performance (CCGP) for cue identity and rank. *P* = 0.0004883 for cue identity, real versus shuffled data; *P* = 0.0004883 for rank, real versus shuffled cell identity; *P* = 0.0004883 for cue identity vs. rank; two-sided Wilcoxon signed-rank test; n = 12 mice. The dashed line represents chance level (0.33). **(L)** Success rate following introduction of the novel rewarded cue (cue C) on day 1. Success rate was calculated using a sliding window of 10 trials. Curves and error bars represent mean ± s.e.m.; n = 12 mice. **(M)** Success rates separated by cue type. Correct trials were defined as licking responses at rewarded cues (A and C) and withholding licking responses at the unrewarded cue (B). Curves and error bars represent mean ± s.e.m.; n = 12 mice. **(N)** Fraction of neurons preferentially responding to different cues on the transition day (day 1). **(O)** Cue preference index across different cue conditions. Center lines, box boundaries, and whiskers indicate median, 25th–75th percentiles, and minimum– maximum, respectively. *P* = 2.548 × 10⁻⁸ for A versus C; P < 1 × 10^-300^ for B versus C; two-sided Wilcoxon signed-rank test; n = 5,458 cells. **(P)** Cosine similarity of cue/rank representations across A/B/Empty and A/B/C conditions, within A/B/Empty conditions, and their differences. Cells from all mice were pooled. **(Q)** Representational geometry in 3-dimensional MDS space under A/B/Empty and A/B/C conditions.

Although cue sequences were randomized across trials, the task contained a stable underlying structure: each cue appeared exactly once per trial, while its rank in a sequence was unpredictable. Consequently, each trial always consisted of two rewarded and one unrewarded slots. To determine whether animals learned this latent reward structure, we designed a reward sequence completion test in which the visual cue at the final slot was masked while reward contingency remained unchanged (Fig. 1B, right). Under this condition, animals could correctly infer reward availability only if they learned the underlying reward structure.

Our primary goal was to characterize how the brain extracts abstract structure from sensory-rich experiences during learning. However, learning is often accompanied by substantial changes in motor variables such as running and licking, making it difficult to dissociate cognitive and movement-related signals. To address this issue, we leveraged an observation from our previous studies that mice could rapidly acquire the full task following a pre-training stage involving only two cues. During pre-training, only cue A and cue B were presented while the third slot was left empty (Fig. 1B, left). This design potentially allowed us to capture a snapshot of neural activity after animals acquired correct task performance but before learning the latent “two-reward” structure, thereby enabling longitudinal tracking of representational changes during structural learning.

We first asked whether the dorsal hippocampus was required for task performance. Mice were trained to expert levels during pre-training and subsequently received bilateral injections of fluorescently conjugated muscimol into the dorsal hippocampus. Post-hoc histological analysis confirmed that injection sites were restricted to the dorsal hippocampus (Fig. 1C). Bilateral hippocampal silencing substantially reduced behavioral performance (Control: 0.8761 ± 0.0322; Sham: 0.8653 ± 0.0302; Muscimol: 0.1644 ± 0.0380, mean ± S.E.M.; Fig. 1D), resulting in non-specific licking across the entire virtual track (Supplementary Fig. 1).

## Disentangled hippocampal representation of cue and rank

We trained eight Thy1-GCaMP6s mice and four Thy1-GCaMP6f mice in the structured cue-guided navigation task, and data from both groups were pooled for analysis. To track hippocampal activity longitudinally, cortex overlying CA1 was aspirated unilaterally and a titanium cannula was implanted for chronic two-photon calcium imaging of CA1 pyramidal neurons throughout learning (Supplementary Fig. 2A). Calcium signals were extracted using Suite2P and fluorescence changes (ΔF/F0) were calculated. Because our study focused on history dependence in visual sequence representations, we further applied deconvolution to estimate putative spiking activity, thereby minimizing contamination from residual calcium signals from previous cues (Supplementary Fig. 2B–E).

We first characterized CA1 activity after animals reached >80% success during pre-training. To improve reliability, analyses were restricted to highly responsive neurons (see Methods). Cells were sorted according to their peak response locations in Empty–A–B trials. In an example animal, distinct neuronal ensembles were observed that were anchored to trial start, cue A, cue B, and trial end (Fig. 1E). Consistently, unsupervised hierarchical clustering based on response similarity across trial types identified three major response clusters corresponding to trial boundaries and visual cues (Supplementary Fig. 3A–D). An additional smaller cluster was preferentially active when the first slot was empty (Supplementary Fig. 3D, lower row).

Importantly, these neuronal ensembles encoded more than instantaneous sensory cues. Instead, they generated sequential activity patterns spanning the interval following cue presentation. Correlation analysis showed that activity within each cluster began slightly before cue onset and persisted until the appearance of the subsequent cue (Supplementary Fig. 3E), resembling previously described landmark-vector cells^28^. Consistently, decoding analysis demonstrated that landmark vector representations in our task were predominantly retrospective, anchored to cues animals were at or just past rather than future ones (Supplementary Fig. 3F).

We next asked how empty slots were encoded in these landmark-vector representations. Because cue spacing varied depending on the position of the empty slot, we examined how CA1 activity represented short and long gaps. Consistent with a previous study^21^, neural activity patterns representing the end of short gaps partially persisted during extended gaps (Supplementary Fig. 3G,H). Interestingly, however, activity patterns associated with empty slots also remained positively correlated across distinct trial conditions (Supplementary Fig. 3G–J). Although representation correlations across various empty slot conditions exhibited substantial variability (smeared in the correlation map rather than forming a tight line), suggesting imprecise distance estimation in the absence of visual cues, the observed correlations suggest that hippocampus encoded empty slots as distinct task positions rather than simply elongated intervals between cues. These findings imply that hippocampal representations capture the ‘three-slot’ structure of the task.

As illustrated by example neurons, CA1 cells were selective not only for visual cue identity but also for sequence rank (Fig. 1F,G), consistent with previous reports showing conjunctive coding of item identity and position in the hippocampus^29^. To quantify these tuning properties, we calculated cue selectivity and rank selectivity indices for individual neurons. Collectively, CA1 neurons exhibited mixed selectivity for both cue identity and rank (Fig. 1H).

To investigate the change of feature selectivity through learning, we compared naïve and expert mice. Behaviorally, learning dramatically reduced anticipatory licking at non-rewarded positions (Supplementary Fig. 4A–C). Both cue and rank selectivity increased significantly throughout learning, as demonstrated by comparisons between naïve and expert animals (Supplementary Fig. 4D–F). Learning also increased overall response magnitude and reduced trial-to-trial variability (Supplementary Fig. 4G,H). In parallel, the fractions of both cue A-and cue B-preferring neurons increased over the course of training (Supplementary Fig. 4I,J).

We next investigated the organization of cue and rank representations at the population level. Because most responsive CA1 neurons generated landmark-vector-like activity spanning from cue onset through the subsequent gap period, we defined these intervals as slot units. Neural activity was averaged within slot units for each trial type, yielding nine population vectors (PVs) corresponding to all combinations of cue identity (A, B, Empty) and sequence rank (1, 2, 3). For example, neural activity during the first slot in A–B–Empty and A–Empty–B trials was averaged to generate the population vector representing “A at rank 1”.

To examine representational geometry, we applied multi-dimensional scaling (MDS) to these nine PVs. MDS minimizes distortion of global distances among data points and has thus been widely used for geometric analysis of neural representations^13,14^. Projection into three-dimensional space revealed a highly disentangled organization of hippocampal representations. The nine PVs formed a triangular prism-like structure, in which cue identity was encoded within a plane while rank occupied a nearly orthogonal dimension (Fig. 1I). Across individual animals, angles between the cue-encoding plane and rank axis were significantly larger than expected from shuffled controls (65.2 ± 5.4° for real data; 33.8 ± 1.8° for shuffle control; Fig. 1J).

Linear dimensionality reduction approach, principal component analysis (PCA), yielded qualitatively similar results. Even without averaging across conditions (18 PVs corresponding to three slot units across six trial types), the first three PCs explained approximately 80% of the variance, suggesting a low-dimensional organization of hippocampal representations (Supplementary Fig. 5). To characterize the single-cell basis of the disentangled low-dimensional representation, we constructed a generalized linear model (GLM) using cue identity, rank, and their interaction as predictors of neural activity. Nonlinear interaction terms (cue × rank) contributed minimally to explained variance (Supplementary Fig. 6). This observation indicates that most variance in the neural activity could be explained by linear combinations of cue and rank, which explains the disentangled representation on the population level.

Disentangled representations have been proposed to facilitate generalization because they allow a common linear decoder to extract one variable independently of the value of another variable. This property can be quantified using cross-condition generalization performance (CCGP). We therefore trained support vector machine (SVM) decoders and measured CCGP for both cue identity and rank. CCGP significantly exceeded shuffled controls for both variables (cue: 0.908 ± 0.024 and 0.440 ± 0.016 for real and shuffled data; rank: 0.548 ± 0.019 and 0.340 ± 0.003 for real and shuffled data; Fig. 1K), consistent with the disentangled organization of hippocampal representations.

## Rapid learning of the newly introduced rewarded cue with preserved but modified representation geometry from the pre-training stage

Following pre-training, we introduced a novel reward-associated visual cue, cue C, into the previously empty slot. Remarkably, mice rapidly learned the newly rewarded cue within only 10–20 trials (Fig. 1L). As expected, errors during the earliest trials were primarily concentrated at cue C. Interestingly, however, errors at later stages occurred predominantly at the unrewarded cue B (Fig. 1M). In another word, animals gradually increased licking probability at cue B while learning the newly rewarded cue, potentially reflecting exploratory behavior following environmental changes.

We next asked how hippocampal representations adapted to the introduction of a novel rewarded cue. One possibility was that learning recruited a new neuronal population selectively representing cue C through rapid plasticity mechanisms ^30–32^. Surprisingly, however, only a small fraction of neurons (∼4%) selectively responded to cue C (Fig. 1N), and the cue preference index for cue C was significantly lower than that for cues A and B (A: -0.18 ± 0.11; B: -0.04 ± 0.13; C: -0.25 ± 0.10; Fig. 1O). More than C-selective neurons, a subset of neurons previously selective for cue A during pre-training responded to both cue A and cue C, suggesting sensitivity to reward-related factors rather than cue identity itself.

To characterize how the introduction of the novel cue reorganizes the populational coding in the hippocampus, we quantify correlations between the A/B/Empty task and A/B/C task. Consistent with single cell analysis, population vectors associated with cue C in the A/B/C task showed strong similarity to representations of the empty slot, as well as the rewarded cue A in the A/B/Empty task (Fig. 1P). Thus, introduction of the novel cue did not completely alter hippocampal representational geometry, but instead modified a pre-existing representational scaffold. The overall disentangled structure was preserved, while representations of the two rewarded cues moved closer together in neural space, reflecting shared reward-related responses (Fig. 1Q).

## CA1 neurons jointly encoded reward expectation and consumption

To further characterize the nature of reward-related responses, we analyzed error trials and catch trials to dissociate sensory and reward-associated factors, including cue identity, reward expectation, and outcome (i.e., reward consumption). In trials in which mice failed to lick at cue C (C-error trials), responses of A/C-selective neurons were markedly reduced to near-baseline levels (Supplementary Fig. 7A,C), suggesting that activity within the cue C slot was primarily associated with reward expectation and/or reward consumption rather than the sensory properties of cue C itself. By contrast, cue B-selective neurons showed no detectable changes during C-error trials (Supplementary Fig. 7D), indicating that their activity primarily reflected sensory information rather than absence of reward. Consistently, in trials where animals incorrectly licked at cue B (B-error trials), responses of cue B-selective neurons partially preserved, though reduced (Supplementary Fig. 7B,E). Reduced activity may arise from decrease in locomotion speed accompanying licking or from exploratory behavioral states induced by environmental uncertainty. Future studies will be required to distinguish between these possibilities.

Error trials alone could not distinguish contributions of reward expectation and reward consumption. To further dissociate these factors, we introduced catch trials (∼10% of trials) in six mice, in which reward delivery was omitted at rewarded cues. When rewards at either cue A or cue C were omitted, the early phase of activity in A/C-selective neurons was markedly reduced, whereas later responses remained relatively preserved (Supplementary Fig. 7F–M). These findings suggest heterogeneous encoding of reward-related information within hippocampal populations, consisting of an early component reflecting reward consumption and a later component resembling landmark-vector activity associated with learned reward expectation independent of actual reward delivery.

Together, these findings suggest that responses associated with the newly introduced rewarded cue primarily reflect reward expectation and reward consumption rather than sensory identity, whereas responses associated with the unrewarded cue are dominated by sensory features.

## Emergence of abstract reward-sequence maps through learning

Because rapid recruitment of novel cue-selective neurons was not observed on day 1, we next asked whether cue-selective representations gradually emerged with extended training. To address this question, mice continued to be trained in the A/B/C task for an additional 3–7 days, after which hippocampal activity was characterized in a late training stage (hereafter referred to as the *late session*). Behaviorally, mice maintained high task performance throughout this period (Fig. 2A).

**Figure 2.**
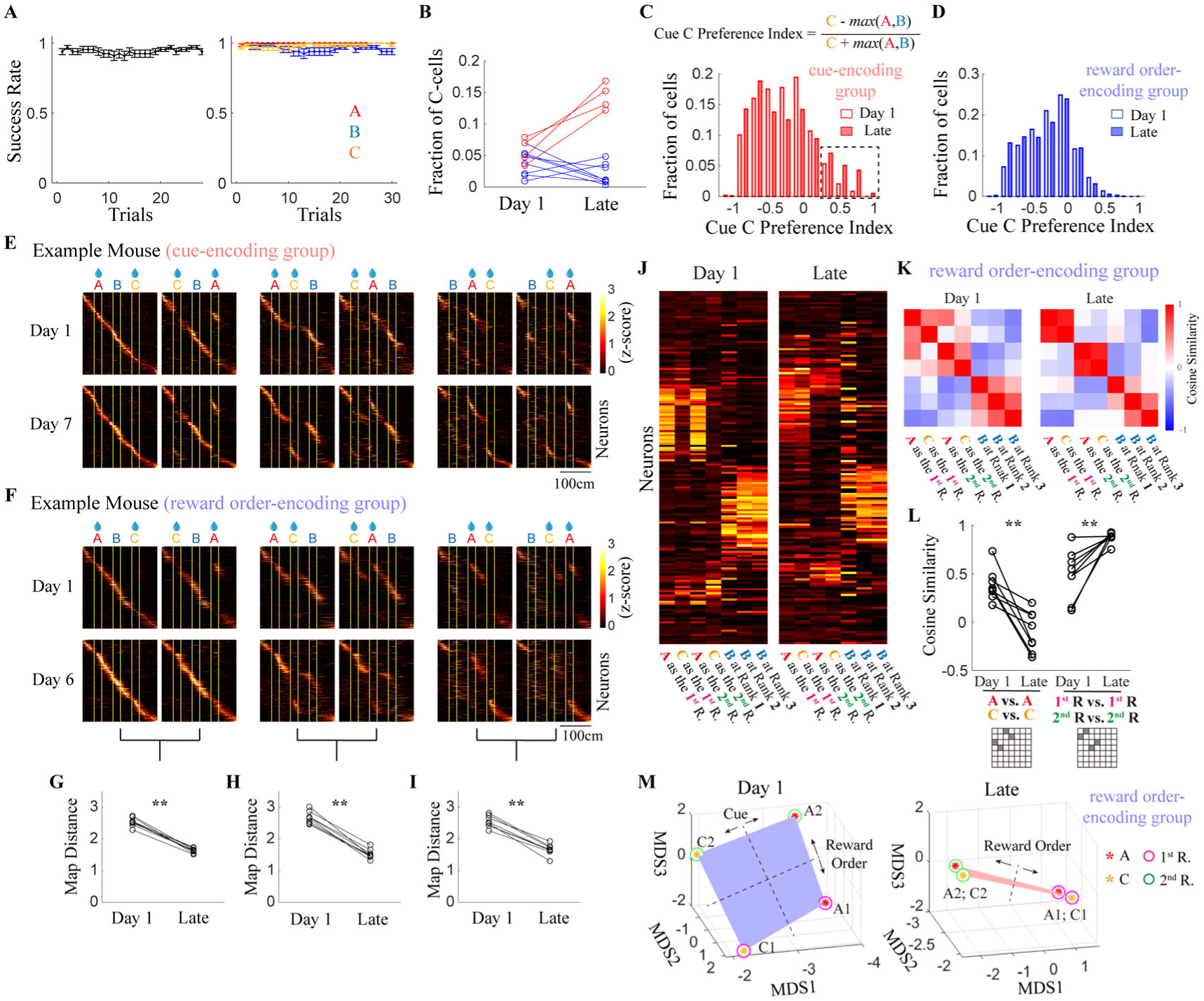
Emergence of abstract hippocampal maps encoding reward sequences through learning. **(A)** Overall (left) and cue-specific (rate) success rates in late session. Curves and error bars represent mean ± s.e.m.; *n* = 12 mice. **(B)** Fractions of cells preferentially responding to the novel cue (cue C). Blue and red indicate cue-encoding and reward order-encoding groups, respectively. **(C)** Distributions of cue C preference index on day 1 (open) and late sessions (filled) in the cue-encoding group (4 mice). *P* < 1 × 10^-300^, day 1 vs. late, Wilcoxon rank sum test, n=1552 cells for day 1 and 1008 cells for late. **(D)** Distributions of cue C preference index on day 1 (open) and late sessions (filled) in the reward order-encoding group (8 mice). *P* < 1 × 10^-300^, day 1 vs. late, Wilcoxon rank sum test, n=3906 cells for day 1 and 1653 cells for late. **(E)** z-scored neural activities in one example mouse in the cue-encoding group. In all heatmaps, cells were sorted based on their activity peaks in the ABC trial type (the first column). **(F)** z-scored neural activities in one example mouse in the reward order-encoding group. All cells were sorted based on their activity peaks in the ABC trial type (the first column) in all plots. **(G)-(I)** Map distances in the reward order-encoding group on day 1 and late session. The map distance is calculated as the summation of mean squared differences for each cell’s tuning curves between the two trial types with the same reward sequence. *P* = 0.0078 for all three pairs, day 1 vs. late; two-sided Wilcoxon signed-rank test; *n* = 8 mice. **(J)** Population vectors in one example mouse in the reward order-encoding group. **(K)** Representation similarity matrix (RSM) on day 1 and late sessions. Cells from all mice in the reward order-encoding group were pooled together. **(L)** Cue similarity and reward-order similarity, calculated as averages among shaded squares in the RSM, on day 1 and late sessions. *P* = 0.0078 for cue similarity, day 1 vs. late; *P* = 0.0078 for reward-order similarity, day 1 vs. late; two-sided Wilcoxon signed-rank test; *n* = 8 mice. Matrix symbols on the bottom indicate squares in representation similarity matrix (J) used for quantifications. **(M)** Representational geometry of rewarded conditions in 3-dimensional MDS space on day 1 (left) and late sessions (right). Red and yellow stars denote cue A and C, respectively. Magenta and green circles denote the first and the second reward for cue A/C, respectively.

In a subset of mice (4/12), the fraction of neurons selectively responding to the novel cue progressively increased, as reflected by a rightward shift in the distribution of cue C preference indices (Fig. 2B,C, red). We refer to these animals as the *cue-encoding group*. As shown in one example animal, a distinct neuronal population selectively tracked cue C (Fig. 2E).

In contrast, the majority of mice (8/12) did not develop selective representations of the novel cue during extended training. In these animals, the fraction of cue C-selective neurons remained low and, in some cases, further decreased (Fig. 2B,D, blue). Importantly, however, this did not indicate a lack of learning-related plasticity. Rather, hippocampal representations underwent a substantial reorganization. To characterize these changes, we grouped the six cue sequences into three pairs according to their reward sequences. Within each pair, rewarded cues A and C were swapped, resulting in distinct sensory configurations but identical reward sequences (e.g. ABC and CBA). On day 1, hippocampal maps differed substantially between paired sequences, largely due to cue A-selective activity. Strikingly, representations within each sequence pair became highly similar during late sessions (Fig. 2F), demonstrated by significant reductions in map distance (2.53±0.05 vs. 1.63±0.03, 2.68±0.07 vs. 1.53±0.05, and 2.54±0.07 vs. 1.66±0.06, for three pairs of maps, respectively; Fig. 2G–I). These findings suggest that hippocampal representations transformed from sensory-oriented maps into reward-sequence representations. We therefore refer to these mice as the *reward order-encoding group*. Because these representations became invariant to the identities of rewarded cues, we refer to them as *abstract maps*.

To further characterize representational transformation within the reward order-encoding group, we performed representational similarity analysis. Population vectors were computed for seven conditions corresponding to A as first reward (A1), C as first reward (C1), A as second reward (A2), C as second reward (C2), and cue B at each sequence rank (B1–B3) (Fig. 2J). Similarity between representations of the two rewarded cues increased throughout learning and became nearly merged in late sessions. Conversely, representations corresponding to first and second reward positions became increasingly separated (cue similarity: 0.38 ± 0.05 vs. -0.10 ± 0.06; reward order similarity: 0.50 ± 0.07 vs. 0.88 ± 0.02; day 1 vs. late session; Fig. 2K,L). Notably, this late-stage representation did not simply reflect pure reward signals, as neural representations were anti-correlated for the first and second rewards in a trial.

We next examined how representational geometry evolved during learning. On day 1, neural representations associated with the four reward-related conditions (A1, C1, A2, C2) occupied approximately the four corners of a rectangle, producing a linearly separable organization in which cue identity and reward order could each be decoded along independent dimensions. In late sessions, however, the cue-encoding dimension largely collapsed while reward-order representations remained preserved, consistent with a progressive abstraction process (Fig. 2M). This geometric transformation was observed both in pooled data and in all individual animals (Supplementary Fig. 8).

We further examined representational transformation using encoding and decoding models. Generalized linear models revealed increased contributions of reward order and sequence rank to neural responses, accompanied by reduced contributions from rewarded cue identity (Supplementary Fig. 9). Consistently, Bayesian decoding assigned similar posterior probabilities to trial types sharing identical reward sequences during late sessions (Supplementary Fig. 10). Together, both encoding and decoding analyses supported a transition toward abstract reward-sequence representations.

We next asked whether reward-order representations reflected learned task structure or merely instantaneous counting of previously consumed rewards within each trial. To distinguish these possibilities, we analyzed catch trials during late sessions. Consistent with observations from day 1, reward omission primarily affected early but not late response components (Supplementary Fig. 11A–F). Individual neurons exhibited heterogeneous responses to reward omission, including complete suppression, complete preservation, delayed responses, or selective reductions of early activity (Supplementary Fig. 11G–I). More importantly, when the first reward was omitted, neuronal ensembles activated by the second reward remained highly similar to those observed under control conditions when the second reward was delivered. Thus, hippocampal representations continued to treat the second reward as the "second" despite omission of the preceding reward (Supplementary Fig. 11A–F). Consistent with this observation, overall population vectors remained highly similar between control and catch trials (Supplementary Fig. 11J,K). These findings suggest that reward-order representations reflect learned task structure rather than immediate reward history in each trial.

Finally, because changes in hippocampal dynamics could potentially arise from alterations in movement patterns, we examined locomotion and licking throughout learning. In the reward order-encoding group, running speed and lick rate showed minimal differences between day 1 and late sessions across the vast majority of spatial bins and trial types (Supplementary Fig. 12A,B). These results indicate that the observed transformation of hippocampal maps cannot be ascribed to changes in movements. More interestingly, speed and lick profiles provided additional behavioral evidence that animals acquired task structure. Divergence in running speed between rewarded and unrewarded conditions emerged substantially earlier at the third slot (∼12 cm before cue onset) than at the first two positions (∼0 cm) (Supplementary Fig. 12C). Similarly, shorter response latency was also seen for lick rates in late sessions (Supplementary Fig. 12D). These observations are consistent with the fact that only reward availability at the third slot could be predicted from task structure.

## A subset of neurons switched tuning properties during learning

We next sought to identify the single-cell changes underlying the population-level transformation of hippocampal representations. Changes in population coding of task-relevant variables may arise through two distinct processes: (1) recruitment of a new neuronal population encoding different variables; or (2) changes in the tuning properties of individual neurons that remain active through learning.

In order to address this question, we tracked activities of single cells through learning. For reliable registration of individual neurons across sessions, imaging planes were carefully aligned each day to maximize overlap with reference maximum-projection images from previous sessions. Cells were registered between day 1 and late sessions in the reward order-encoding group using CellReg^33^. Six mice with reliable alignment quality were included in this analysis, with approximately 50% of imaged neurons successfully registered across sessions (Supplementary Fig. 13).

To characterize tuning properties, we quantified the mutual information between trial-by-trial neural activity and multiple task variables, including cue identity, sequence rank, reward, and reward order (Supplementary Fig. 14A–E; see Methods for details). All registered neurons were included in this analysis. Example neurons revealed substantial changes in tuning properties across learning (Fig. 3A–F). Consistent with our population-level analyses, mutual information associated with cue identity decreased over learning, whereas mutual information associated with reward order increased (Cue: 0.1275 ± 0.0025 vs. 0.1005 ± 0.0021; Reward order: 0.0428 ± 0.0010 vs. 0.0588 ± 0.0018; day 1 vs. late sessions; Fig. 3G).

**Figure 3.**
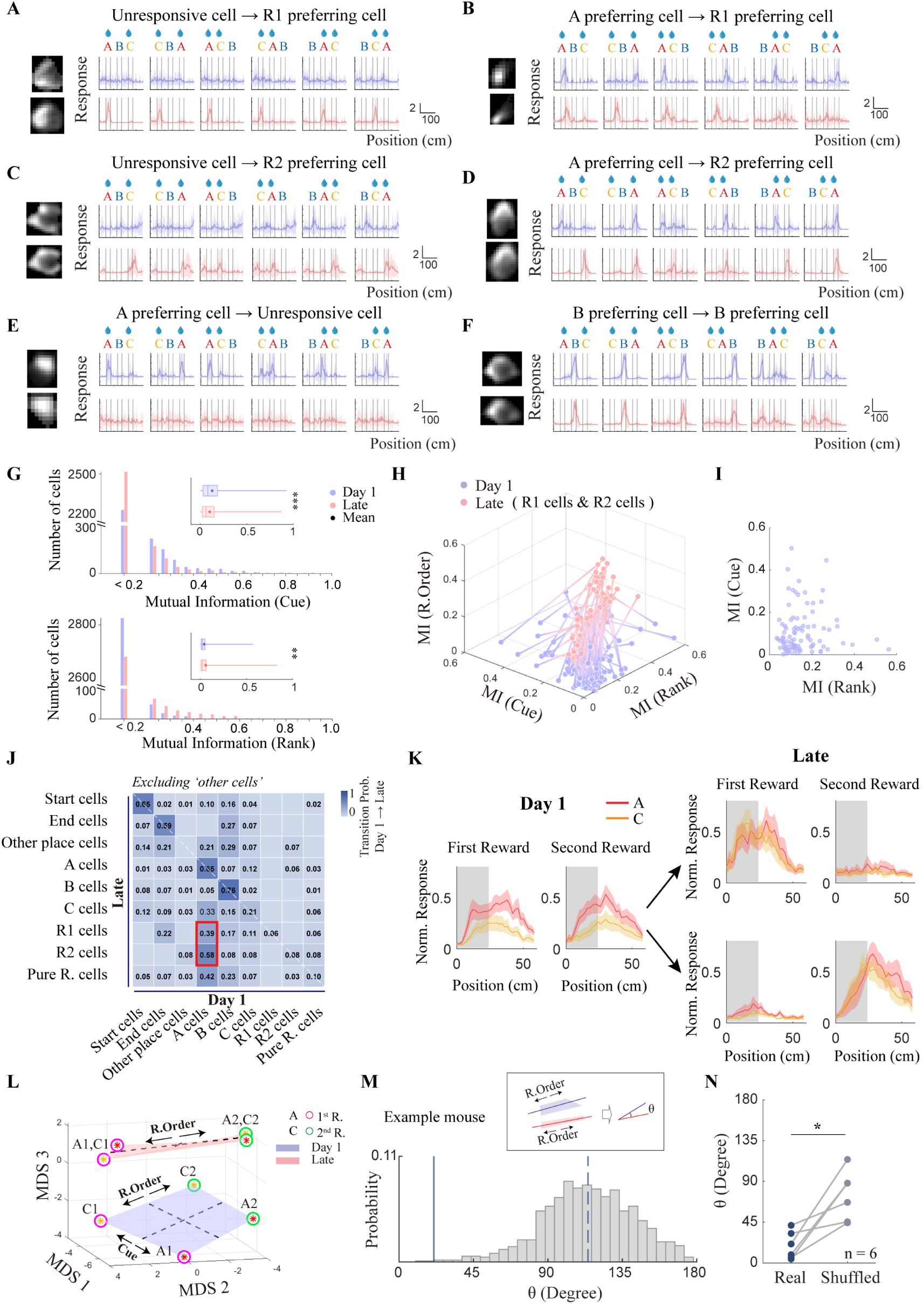
Individual cells switched tuning properties during learning with stable population coding dimension of task variables. (A)-(F) Deconvolved calcium responses across the six trial types of example cells on day 1 (purple) and late sessions (pink). Traces represent mean z-scored activity. Pictures on the left show registered images of the corresponding cells across sessions. **(G)** Distributions of mutual information for cue (upper) and reward order (bottom) on day 1 and late sessions. Inset: mutual information for cue and rank on day 1 and late sessions. Center lines, circles, box boundaries, and whiskers indicate the median, mean, 25th–75th percentiles, and minimum–maximum values, respectively. Cells included in the analysis included all registered neurons except those classified as ‘other’ in both day 1 and late sessions. P = 4.517 × 10-9 and 0.0012 for cue and reward order, respectively, day 1 vs. late; two-sided Wilcoxon signed-rank test; n = 2914 cells. **(H)** Changes in mutual information for rank, cue, and reward-order variables from day 1 to late sessions. Cells included in this analysis were neurons classified as reward-order cells (R1 or R2) in late sessions; n = 95 cells. **(I)** Mutual information for cue and rank on day 1 for neurons classified as first reward or second reward cells in late sessions (as shown in panel H). **(J)** Cell-type transition probabilities from day 1 to late sessions. Probabilities smaller than 0.01 are not labeled in the plot. Each row represents probabilities of ‘source’ cell types transitioned to one specific cell type on late sessions (probabilities on each row summed to 1). Two red squares marked largest sources of first-reward and second-reward preferring cells, respectively. Only neurons that were not classified as “other” on either day 1 or late sessions were included to better visualize transitions among the remaining cell types; n = 694 cells. **(K)** A subset of cue A-preferring cells on day 1 (left) developed into two populations of cells which preferred the first reward (right, top) and the second reward (right, bottom) in late sessions, respectively. Normalized responses to cue A (red) and cue C (yellow) were plotted. Curves and shades represent mean ± s.e.m., respectively. Cells classified as ‘A cell’ on day 1 and ‘R1 cell’ or ‘R2 cell’ in late sessions were included in this analysis. **(L)** Representational geometry in 3-dimensional MDS space on day 1 and late sessions. Red and yellow stars denote cue A and C, respectively. Magenta and green circles denote the first and the second reward for cue A/C, respectively. Dashed lines indicate connections between centers of different variables. Cells from all mice were pooled together. **(M)** Angles between the reward-order dimensions on day 1 and late sessions in an example mouse (H10). The solid and dashed lines indicate mean angles calculated from real (21.01°) and shuffled (114.20°) data, respectively. Bars represent the distribution of angles in shuffled data. *P* = 0.0021; one-sided permutation test based on 1,000 shuffled datasets; n = 1399 cells. **(N)** Angles between the reward-order dimensions on day 1 and late sessions for each mouse. *P* = 0.0313; two-sided Wilcoxon signed-rank test; n = 6 mice.

To investigate how reward order-selective representations emerged, we next performed mutual information-based cell classification (Supplementary Fig. 14F). We chose relatively strict classification criteria to focus on neurons that strongly encoded certain variables. It should be noted that assigning neurons into discrete classes serves only as an approximation for visualizing representational transformations, as many CA1 neurons exhibited mixed selectivity for multiple variables (Supplementary Fig. 15). For example, some neurons responded preferentially to both trial onset and cue B (Supplementary Fig. 15F), whereas others selectively encoded the first or second reward only in specific trial types (Supplementary Fig. 15C). Nevertheless, this classification framework allowed us to identify the dominant tuning property of individual neurons.

Interestingly, most reward order-selective neurons emerged from a population categorized as “other” on day 1, defined as neurons lacking strong selectivity for any tested variables (Supplementary Fig. 14G,H). Consistent with this observation, most reward order-selective neurons during late sessions carried relatively weak cue and rank information on day 1 (Fig. 3H,I). To further identify neurons that exhibited significant tuning in both sessions while changing selectivity over learning, we analyzed day 1 tuning identities conditioned on late-session tuning categories, excluding the “other” population. This analysis revealed that the majority of both first reward-selective and second reward-selective neurons originated from cue A-selective neurons present on day 1 (Fig. 3J). Consistently, a subset of neurons encoded the rewarded cue on day 1 (Fig. 3K, left) preserved their coding for reward but developed into two reward-order selective groups through learning (Fig. 3K, right). Transformation between preference across reward and non-rewarded conditions was rare.

Together, these findings suggest that the emergence of abstract reward sequence representations involved at least two processes: recruitment of previously weakly tuned neuronal populations and transformation of cue-oriented representations into reward order-oriented representations.

We next performed geometric analyses using only registered neurons, allowing direct comparison of representational dimensions across sessions (Fig. 3L). Consistent with analyses using unregistered populations, the cue-encoding dimension largely collapsed during late learning. Interestingly, the reward-order dimension remained relatively stable across sessions. Angles between reward-order dimensions from day 1 and late sessions were significantly smaller than expected from shuffled cells in both the example and population mice (19.17 ± 6.25 ° for real data; 74.53 ± 11.18° for shuffled data; Fig. 3M,N).

These findings suggest that, despite substantial tuning changes at the single-cell level, population-level representations preserved stable task-relevant coding dimensions. As a result, downstream circuits could potentially extract reward-order information without requiring extensive modification of decoding mechanisms. More broadly, these results raise the possibility that hippocampal representational drift may be preferentially constrained within dimensions orthogonal to task-relevant information (i.e., the null space).

## Formation of a parsimonious Markov graph by integrating physical and abstract information

We showed that animals in the reward order-encoding group developed abstract representations of reward sequences, distinguishing first and second rewards. These abstract reward sequences correlated with, but did not fully overlap with, the physical structure of the task, namely sequence rank along the virtual corridor. Specifically, rewards at rank 1 and rank 3 always corresponded to the first and second reward, respectively, whereas rewards at rank 2 could correspond to either. We next sought to determine how hippocampal representations integrate physical sequence information (rank) and abstract sequence information (reward order) into a unified state map.

One possibility is that the hippocampus represents the task using a simple reward-rank map (Fig. 4A). Such a representation naturally contains partial information about reward order. However, a pure rank representation cannot fully specify future state transitions. Specifically, the rank-2 reward state can move to either a rewarded or non-rewarded future state depending on the preceding trajectory (brown arrows in Fig. 4A). An alternative possibility is that the hippocampal representation forms a pure reward-order map. Neural activities separately map the first and second rewards, but do not encode the physical rank (Fig. 4B). Similar to the pure rank map, such a pure reward-order map contains ambiguous states that cannot, by itself, determine the transition probability. Therefore, neither the pure rank map nor the pure reward-order map satisfies the Markov property (i.e., history independence), which has been proposed highly beneficial for learning and decision making.

**Figure 4.**
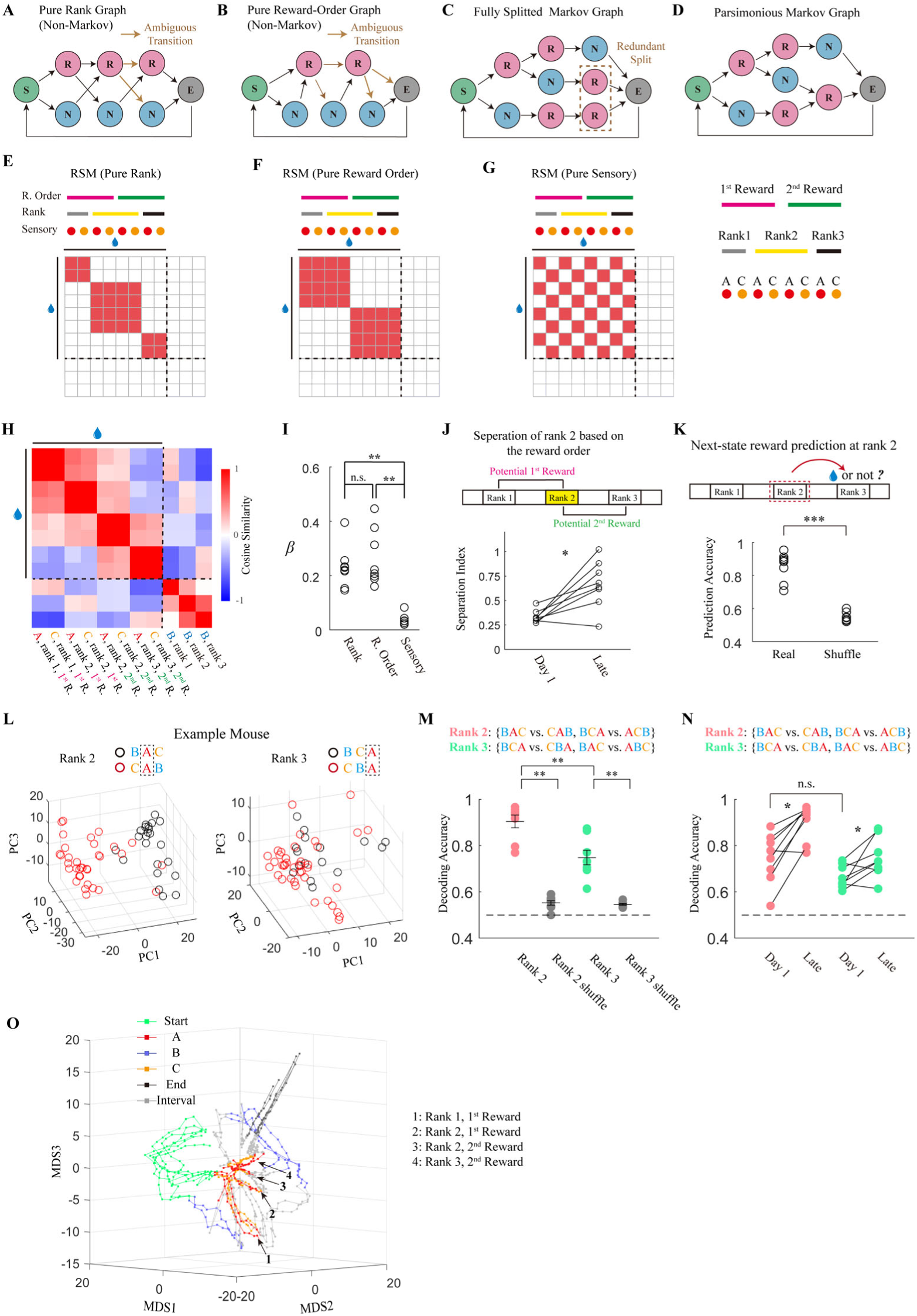
Hippocampal representation manifested a parsimonious Markov graph. (A)-(D) Different models of reward sequence encoding. Green, red, blue, and dark gray represent trial start, reward, non-reward, and trial end states, respectively. In A and B, brown arrows indicate history-dependent state transitions. **(E)-(G)** Theoretical representation similarity matrix (RSM) for coding different task-relevant variables. Red squares indicate high similarity. Only the similarity across different rewarded conditions (indicated by the water drop symbol) were shown. **(H)** RSM with real neural responses in the reward order-encoding group. Cells were pooled from all mice in the reward order-encoding group. **(I)** Regression coefficients for different variables. All data point were included in statistics. *P* = 0.0025, Friedman test; *P* = 0.0078 (α=0.05), sensory vs. reward order; *P* = 0.0078 (α=0.0167), sensory vs. rank; *P* = 0.6406, rank vs. reward order; two-sided Wilcoxon signed-rank test; n=8 mice. **(J)** Change of separation index at rank 2 through learning. Separation index was defined as the difference between representation similarity (cosine similarity) with match (A and C both as the 1^st^ or 2^nd^ reward) and unmatched (A and C with one as the 1^st^ whereas the other as the 2^nd^ reward) reward order conditions. *P* = 0.0156, day 1 vs. late, two-sided Wilcoxon signed-rank test; n=8 mice. **(K)** Decoding accuracy of reward availability at rank 3 using neural activity at rank 2. SVM was used for decoding. *P* = 1.554 × 10^-4^, real vs. shuffled data, two-sided Wilcoxon signed-rank test; n=8 mice. **(L)** Neural responses at the same rewarded cue (cue A) at rank 2 (left) and rank 3 (right) in one example mouse. Each circle represents one trial. Red and black denotes different past trajectories. **(M)** Decoding accuracies of past trajectories at rank 2 (pink) and rank 3 (green). *P* = 0.0078 for both rank 2 and rank 3, real vs. shuffled cell identity, two-sided Wilcoxon signed-rank test; n=8 mice. **(N)** Decoding accuracies of past trajectories at rank 2 (pink) and rank 3 (green) on day 1 and late sessions. *P* = 0.0078 for both rank 2 and rank 3, day 1 vs. late; *P* = 0.0781 for day 1, rank 2 vs. rank 3; two-sided Wilcoxon signed-rank test; n=8 mice. **(O)** Neural activity trajectories in 3-dimensional MDS space. Six trial types were plotted together. Cells were pooled from all mice in the reward order-encoding group.

A Markovian map can be constructed by integrating the rank and reward-order information. One way to integrate these two pieces of information is that hippocampal representations split all trajectory-dependent states into separate hidden states, with each distinguishable trajectory forming an independent branch (Fig. 4C). We refer to this map as a ‘full Markov graph’. In this scenario, each reward state carries combined information of rank and reward order. Although this representation fully specifies future transitions, it may contain redundant state splitting. Separating rank-3 reward states according to trajectory history is unnecessary because these distinctions do not alter subsequent transitions.

We therefore considered a fourth possibility: the hippocampus forms a parsimonious Markov graph (Fig. 4D), in which only states necessary for predicting future transitions are separated. Under this framework, rank-2 rewards would split into distinct hidden states according to reward order, whereas rank-3 rewards would remain unified.

To distinguish among these possibilities, we first performed representational similarity analyses using the population vectors organization shown in Fig. 4E–G, representing all possible combinations of rank, reward order, and sensory cue. Unlike previous analyses (Fig. 2), reward population vectors from different ranks were analyzed separately. This organization generates distinct representational similarity patterns depending on which variable hippocampal activity primarily encodes. We next used linear regression to quantify contributions of each candidate similarity map to observed data. Hippocampal representations in late sessions were primarily explained by rank and reward-order components, with only a minimal contribution from cue identity (Fig. 4H,I; regression coefficient: 0.23 ± 0.03, 0.26 ± 0.04, and 0.04 ± 0.01, for rank, reward order, and sensory, respectively). This pattern is consistent with Markov graphs, either a fully split one (Fig. 4C) or a parsimonious one (Fig. 4D), in which reward states emerge from combinations of rank and reward order. Learning promoted the encoding of reward order, beyond physical rank, by increasing reward order-dependent state separation at rank-2 (separation index: 0.33 ± 0.02 vs. 0.67 ± 0.09, day 1 vs. late; Fig. 4J).

To investigate how coding dimensions for rank and reward order are organized, we next examined representational geometry in neural state space. Population vectors were projected into a three-dimensional space. Within the first two MDS dimensions, rank-2 reward states showed minimal separation on day 1 but became clearly separated during late learning along the rank-coding dimension (Supplementary Fig. 16A,B). Specifically, the first-reward state was closer to rank-1 while the second-reward state was closer to rank 3. This is reasonable since rank-1 and rank-3 rewards correspond to first and second rewards, respectively. Consistent with this observation, the distance between rank-2 reward states measured along the rank axis increased significantly throughout learning (Supplementary Fig. 16E–H).

Interestingly, rank-2 reward states also separated along a third dimension orthogonal to the rank coding axis (Supplementary Fig. 16C,D). Separation along this orthogonal axis also increased during learning, although this effect did not reach statistical significance in our dataset (Supplementary Fig. 16I). These findings suggest that abstract and physical variables may organize in partially overlapping representational dimensions. Previous studies have reported orthogonal representations for entirely independent variables such as physical space and accumulated evidence strength^19^. By contrast, our results suggest that when abstract and physical variables exhibit partial correlations, their neural representations may contain both parallel and orthogonal components.

A Markov graph is predictive. Each state determines the distribution of transition probability into the next one. To demonstrate this property, we performed decoding analysis to test whether the hippocampal activity at rank 2 could predict the reward availability at rank 3. As expected, the decoding accuracy is significantly higher than the shuffled control (Fig. 4K; 0.855 ± 0.031 for real data and 0.549 ± 0.010 for shuffled control), showing that hippocampal activity is not only indicative about the current reward, but also predictive for future ones.

Having established the Markovian nature of hippocampal map, we sought to distinguish the ‘full Markov graph’ vs. ‘parsimonious Markov graph’. The key difference between these two models is whether trajectory information could be decoded from rank-3 reward state(s). Although previous trajectory information could be decoded at rank 3 above the shuffled control, decoding accuracy was significantly lower than rank-2 states (rank-2: 0.904 ± 0.027 vs. 0.552 ± 0.010; rank-3: 0.748 ± 0.031 vs. 0.546 ± 0.004; real vs. shuffled data; Fig. 4L,M). Importantly, this difference was insignificant on day 1 and emerged through learning (rank-2: 0.744 ± 0.039; rank-3: 0.665 ± 0.018; decoding accuracies on day 1; Fig. 4N). Representational similarity analyses for trials with different history cues yielded consistent conclusions (Supplementary Fig. 17). Compared to rank 2, reward states at rank 3 remained partially merged.

Together, these findings support the hypothesis that hippocampal representations form a parsimonious Markov graph. This latent-state structure was further apparent when neural activity trajectories from all six trial types were visualized jointly in low-dimensional space.

Reward-related activity segregated into four distinct segments corresponding to the four reward states predicted by the parsimonious Markov graph (compare Fig. 4O and Fig. 4D).

## Hippocampal map abstraction predicts structure-based behavioral strategies

Hippocampus forms a predictive map of the task. Do animals utilize this map to predict rewards behaviorally? We leveraged the spontaneously emerging *cue-encoding* and *reward order-encoding* groups, in which representational abstraction either did or did not occur, to test whether abstract hippocampal maps were associated with structure-based behavioral strategies.

Specifically, we tested whether animals could infer reward availability at the third slot when visual cues were masked, potentially using previously encountered rewards within the sequence to infer reward structure (Fig. 5A; see Methods). Strikingly, reward order-encoding mice performed significantly better than cue-encoding mice during probe trials with masked third cues (Fig. 5B). This difference was particularly pronounced when the masked cue corresponded to the unrewarded condition (success rates: 0.2179 ± 0.0857 vs. 0.6383 ± 0.0581 for the unrewarded condition, 0.6824 ± 0.0426 vs. 0.8820 ± 0.0056 for the rewarded condition, cue-encoding group vs. reward order-encoding group; Fig. 5C,D). Interestingly, even cue-encoding animals performed relatively well when the third masked cue corresponded to a rewarded condition. This observation suggests that these animals may adopt a simpler heuristic strategy when the third slot was un-cued—effectively treating all locations without an explicitly learned non-rewarded cue as potentially rewarded. Overall, despite equivalent behavioral performance when sensory cues were fully available, the two groups exhibited markedly distinct behaviors under cue-masking conditions. These findings suggest that hippocampal representational modes strongly predict the behavioral strategies spontaneously adopted by individual animals.

**Figure 5.**
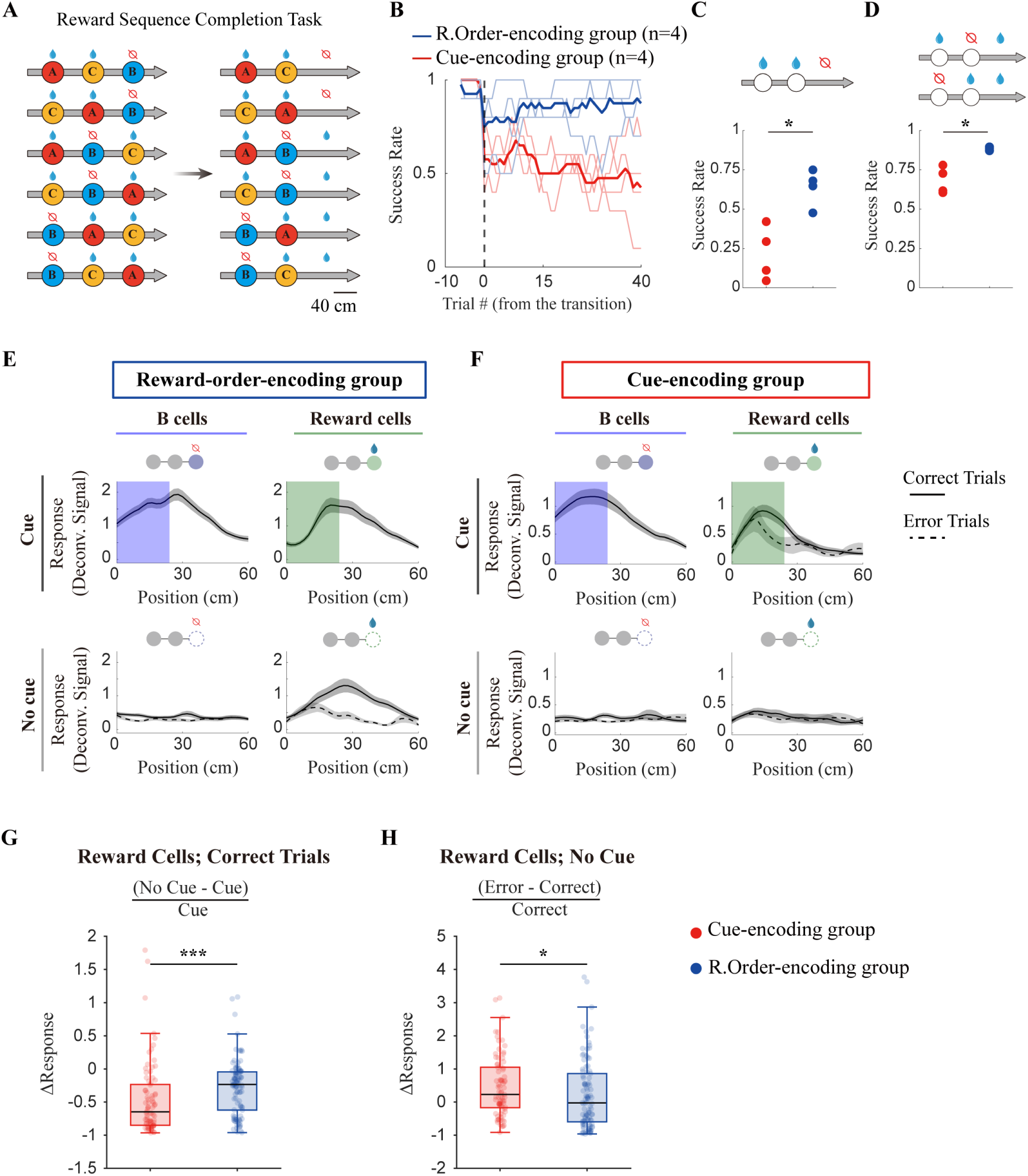
Abstract hippocampal representation predicted structure-based behavioral strategy. **(A)** Schematic of the reward sequence completion test. **(B)** Success rates in probe trials with the third cue masked across different groups of mice. Success rates were calculated using a sliding window of 10 trials. Dark curves indicate the mean, and light curves indicate individual mice; n = 4 mice for the reward order-encoding group (blue) and n = 4 mice for the cue-encoding group (red). **(C)** Success rates in probe trials in which the last slot (cue masked) was unrewarded. *P* = 0.0286; two-sided Wilcoxon rank-sum test; n = 4 mice, respectively. **(D)** Success rates in probe trials in which the last slot (cue masked) was rewarded. *P* = 0.0286; two-sided Wilcoxon rank-sum test; n = 4 mice, respectively. **(E-F)** Responses of B-preferring cells (left column) and second reward-preferring cells (right column) in the presence (upper panels) or absence (lower panels) of the third cue in correct (solid lines) and error trials (dashed lines). Curves and shaded areas represent mean ± s.e.m., respectively. Panels E and F correspond to the reward order-encoding group and cue-encoding group, respectively. **(G)** Normalized response differences of reward-preferring cells between trials in the presence and absence of the third cue (blue: reward order-encoding group; red: cue-encoding group). Center lines, box boundaries, and whiskers indicate median, 25th–75th percentiles, and minimum– maximum, respectively. Outliers were defined as point that fall beyond 1.5 times interquartile range from the nearest quartile. It should be noted that outliers were just for visualization purposes. All data point were included in statistics. *P* = 1.917 × 10^-4^; two-sided Wilcoxon rank-sum test; n = 102 cells for the reward order-encoding group and n = 81 cells for the cue-encoding group. **(H)** Normalized response differences of reward-preferring cells between error and correct trials in the reward order-encoding group (blue) and cue-encoding group (red). Center lines, box boundaries, and whiskers indicate median, 25th–75th percentiles, and minimum–maximum, respectively. Outliers were defined as point that fall beyond 1.5 times interquartile range from the nearest quartile. It should be noted that outliers were just for visualization purposes. All data point were included in statistics. *P* = 0.0224; two-sided Wilcoxon rank-sum test; n = 102 cells for the reward order-encoding group and n = 81 cells for the cue-encoding group.

We next examined hippocampal activity during trials in which the third cue was masked. In the reward order-encoding group, neuronal responses remained elevated when the masked cue corresponded to a rewarded condition during correct trials (solid lines, Fig. 5E, right). Consistent with an association to behavioral performance, responses were significantly reduced during error trials in which animals failed to lick at rewarded masked locations (dashed lines, Fig. 5E, right). In contrast, responses associated with the non-rewarded cue (cue B) were largely abolished after cue masking, suggesting that these responses primarily reflected sensory information (Fig. 5E, left). These observations reveal an asymmetry between hippocampal representations of rewarded and non-rewarded conditions. In contrast to the reward order-encoding group, responses to both rewarded and non-rewarded conditions were substantially reduced following cue masking in cue-encoding animals, consistent with predominantly sensory-driven representations (Fig. 5F). Quantification of neural activity changes in response to cue masking and behavioral error across groups further supported this distinction (Fig. 5G,H), suggesting that hippocampal representations remain sensory-oriented in cue-encoding mice but become structure-oriented in reward order-encoding animals.

Behavioral strategies are shaped dynamically by experience. We next asked whether cue-encoding animals could be shifted toward structure-based strategies. Previous studies have shown that exposure to multiple related problems facilitates learning of generalized schemas ^2,3^. We therefore continued training using two additional rewarded cues while maintaining the same unrewarded cue B (Supplementary Fig. 18A).

Consistent with the previously proposed “excluding B” strategy, both cue-encoding and reward order-encoding animals performed relatively well when novel rewarded cues were introduced, without requiring extensive retraining (Supplementary Fig. 18B). More importantly, exposure to novel rewarded cues promoted a transition of hippocampal representations toward reward-sequence encoding in animals originally classified as cue-encoding. Specifically, similarity between the two rewarded cues (A and C) increased whereas similarity between representations of first and second rewards decreased (Supplementary Fig. 18C–E). Accompanying this representational transformation, animals also exhibited improved performance during structure-inference probe trials with masked cues (Supplementary Fig. 18F–H).

Together, these findings suggest that experience-dependent abstraction of hippocampal representations predicts the emergence of structure-based behavioral strategies. We note that our experiments do not rule out the possibility that mice could predict the cue identity at rank 3 based on the task structure, and in turn infer the reward availability using cue-reward association. However, the strong association between cue-invariant neural responses and behavioral performance in the reward sequence completion task implies the implementation of a cue-insensitive reward counting strategy, which is simpler than prediction of specific cues.

## Computational principles underlying structural map formation

Having established the emergence of abstract structural maps and their predictive relationship to behavioral strategy, we next sought to identify computational principles underlying the formation of such representations. Because hippocampal representations resembled a parsimonious Markov graph, we hypothesized that two computational principles may be essential: (1) next-state prediction during learning and (2) strong regularization of state representations. These two principles correspond naturally to the *Markov* and *parsimonious* aspects of the latent-state map, respectively.

To test whether these principles were sufficient for structural map formation, we implemented them in several distinct computational frameworks (Fig. 6A), including two state models and two neural network models. The first model is a hierarchical Dirichlet process hidden Markov model (HDP-HMM, Fig. 6B), which extends the traditional HMM by inferring the number of hidden states from data. The second model, termed clone structure cognitive graph, or CSCG ^34^, is an HMM initialized with a large number of potential latent states associated with identical sensory observations, thereby facilitating hidden-state inference (Fig. 6E). The third model consisted of a predictive recurrent neural network (RNN), trained to predict reward availability at both the current and subsequent time steps (Fig. 6H), with strong regularization implemented through 60% dropout within the hidden layer. The fourth model was the Tolman-Eichenbaum Machine, or TEM ^35^, which combines recurrent prediction mechanisms with a Hopfield-like associative network and was originally proposed as a model of hippocampal function (Fig. 6N).

**Figure 6.**
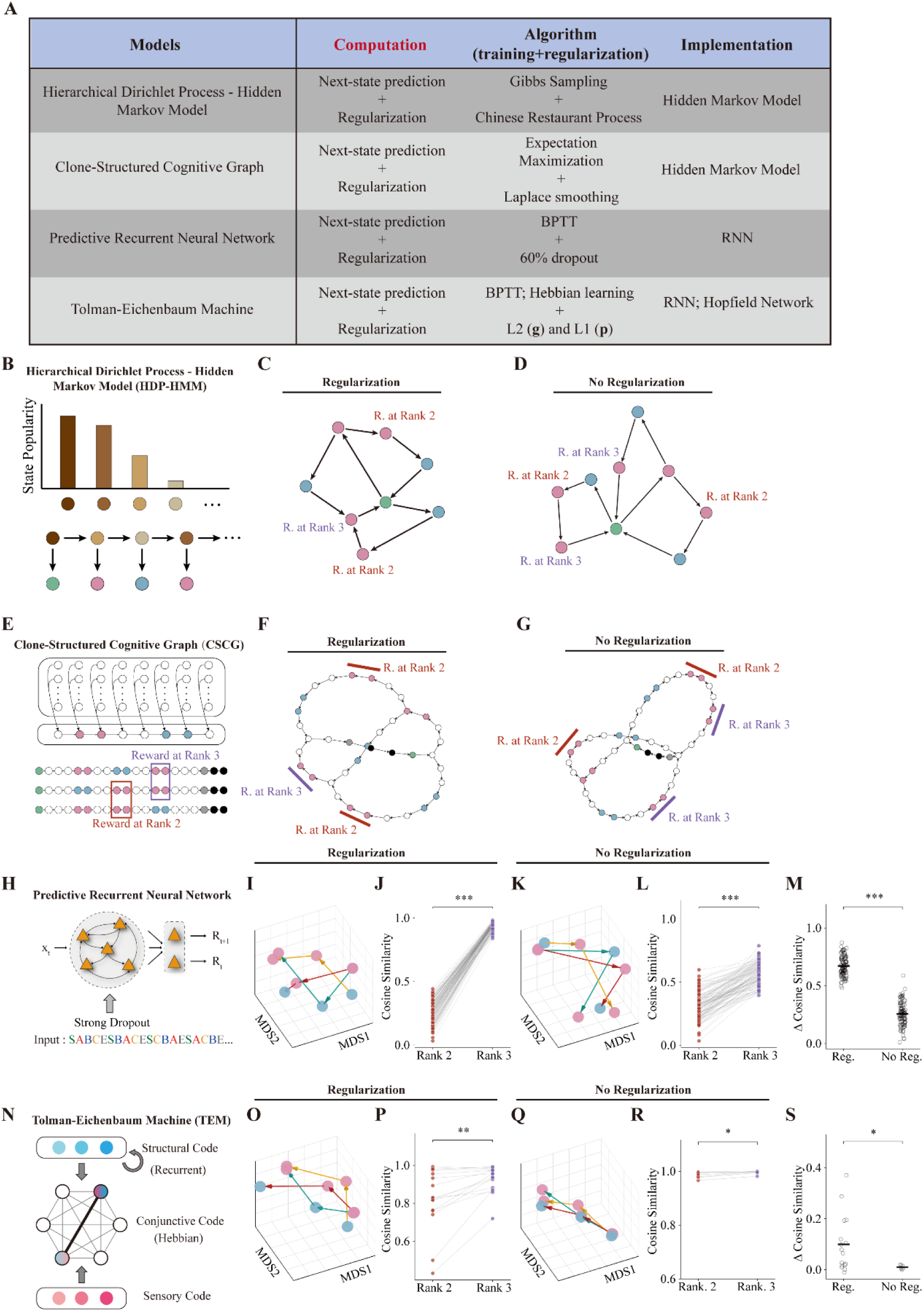
Next-state prediction and strong regularization promote the formation of structural map in computational models. **(A)** Comparison of four models (HDP-HMM, CSCG, P-RNN, TEM) on levels of computation, training/regularization algorithm, and implementation. **(B)** Schematic of hierarchical Dirichlet process hidden Markov model (HDP-HMM). State transition probabilities were sampled from a Dirichlet process centered at a global state distribution. The number of states was inferred from observed sequences. **(C)-(D)** State maps of HDP-HMM after training. Red, reward at rank 2; purple, reward at rank 3. Concentration parameter γ was set to 3 (regularization) and 100 (no regularization). **(E)** Schematic of clone-structure cognitive graph (CSCG). Each sensory input is modeled with 100 clones. Green, red, blue, gray, black, white represent trial start, reward, non-reward, trial end, inter-trial interval, and inter-cue interval, respectively. **(F)-(G)** Trained CSCG state transition graph. Red, reward at rank 2; purple, reward at rank 3. Pseudocount κ was set to 1e-6 (regularization) and 1e-10 (no regularization). **(H)** Schematic of recurrent neural network (RNN). Triangles, units; xₜ, sensory cue sequence; Rₜ, reward output. Dropout applied to hidden layer during training. **(I)** MDS plot of RNN hidden layer representations under the regularized condition. Red points: reward; blue points: non-reward. Colored arrows indicate transition directions: yellow, NRR; red, RRN; green, RNR. **(J)** Comparison of cosine similarity for reward at rank 2 (red) vs. rank 3 (purple) of RNN hidden layer representations under the regularized condition. *P* = 4 × 10^-18^, two-sided Wilcoxon signed-rank test, n = 100 models with different initialization. **(K)** Same as (I) but under the unregularized condition. **(L)** Same as (J) but under the unregularized condition. *P* = 4 × 10^-18^, two-sided Wilcoxon signed-rank test, n = 100 simulations. **(M)** Difference in cosine similarity (rank 3 minus rank 2) under regularized vs. unregularized conditions. P = 4 × 10^-34^, Wilcoxon rank sum test. **(N)** Schematic of Tolman-Eichenbaum Machine (TEM). Structural codes (blue) are updated recurrently; sensory codes (red) are factorized inputs. These two are conjunctively combined into conjunctive codes. The memories of conjunctive code are stored in a Hopfield network with Hebbian weights. **(O)** MDS plot of TEM structural code representations under the regularized condition. **(P)** Comparison of cosine similarity for reward at rank 2 (red) vs. rank 3 (purple) of TEM structural code representations under the regularized condition. *P* = 0.0001, two-sided Wilcoxon signed-rank test, n = 16 models with different initialization. **(Q)** Same as (O) but under the unregularized condition. **(R)** Same as (P) but under the unregularized condition. *P* = 0.0313, two-sided Wilcoxon signed-rank test, n=7 simulations. **(S)** Difference in cosine similarity (rank 3 minus rank 2) under regularized vs. unregularized conditions. *P* = 0.0119, Wilcoxon rank sum test.

From the perspective of David Marr’s framework, these models differ substantially at the levels of algorithm and implementation while sharing similar computational objectives. Interestingly, despite these differences, all four models generated representational structures resembling those observed experimentally in reward order-encoding animals.

Both HDP-HMM and CSCG produced a state topology equivalent to the experimentally observed latent-state map (Fig. 4D and O), characterized by splitting of rank-2 reward states (Fig. 6C and F, red) while preserving merged rank-3 reward states (Fig. 6C and F, purple). Similarly, predictive RNNs trained to predict future rewards generated hidden-layer representational similarity structures closely resembling experimental observations (compare Supplementary Fig. 19 and Fig. 4H). An alternative RNN only trained to output the current reward did not produce a predictive map as observed in experiments (Supplementary Fig. 19). TEM produced similar population representation compared to predictive RNN, although the two networks exhibited different tuning curves for individual neurons (Supplementary Fig. 20). Both predictive RNNs and TEM exhibited splitting of rank-2 reward states together with partial merging of rank-3 reward states, as revealed by both decoding analyses and representational geometry (Fig. 6I,J, 6O,P; see Methods and Supplementary Fig. 21 for details of our TEM model).

We next examined the importance of regularization by removing regularization mechanisms from each model, a manipulation difficult to perform experimentally. Across all model architectures, removal of regularization substantially disrupted representational organization. In HDP-HMM and CSCG, removing regularization generated additional splitting of rank-3 reward states (Fig. 6D and G). Likewise, in both RNN and TEM, eliminating regularization significantly reduced the difference in history-decoding performance between rank-2 and rank-3 states (Fig. 6K–M, 6Q–S).

Notably, regularization mechanisms in RNN and TEM did not explicitly constrain the number of latent states. Instead, they restricted representational dimensionality. Nevertheless, dimensionality constraints alone naturally promoted formation of parsimonious state representations. These findings suggest that low-dimensional constraints may provide a simple computational mechanism for preventing excessive state fragmentation.

Together, these results suggest that predictive learning and representational regularization may constitute core computational principles underlying the formation of abstract structural maps. Future studies will be required to determine how these computations are implemented in biological neural circuits.

## Reward-dominated representations in the OFC

Finally, we asked whether abstract representations similar to those observed in hippocampus could also emerge in other cognitive brain regions. The orbitofrontal cortex (OFC) has been implicated in encoding abstract task structure and cognitive maps. We therefore implanted GRIN lenses and performed two-photon Ca^2+^ imaging of OFC neurons with the same behavioral training protocol used for hippocampal recordings (Supplementary Fig. 22). OFC activity was strongly concentrated at rewarded zones (Fig. 7B,C). Activity in most OFC neurons constituted two nearly identical ‘goal-progression’ sequences per trial around reward onset locations (Fig. 7B and Supplementary Fig. 23A), whereas a small subset of neurons was suppressed by rewards (Supplementary Fig. 23B). Unlike reward preferring neurons, reward suppressed OFC neurons showed persistent activation at non-rewarded locations, rather than forming a sequence. As a result, for each trial type (cue sequence), neural activity trajectory parked at similar locations for non-reward zones (track start, track end, and cue B) and moved along a circle each time approaching and exiting reward zones (Fig. 7F). In contrast to the hippocampus, which encoded the two rewards differently based on rank and reward order, OFC representations were highly similar across the two rewarded cues. This pattern was evident in both representational similarity analyses (Fig. 7D; compare Fig. 4H for the HPC) and generalized linear model analyses (Fig. 7E), suggesting that OFC did not discriminate the two rewards based on cues, ranks, or reward orders, in our virtual navigation task. Given the lack of strong selectivity of cue, rank, or reward order, neural trajectories corresponding to the two reward zones were nearly identical. With all six trial types plotted together, OFC neural trajectories showed twelve overlapping circles (Fig. 7F).

**Figure 7.**
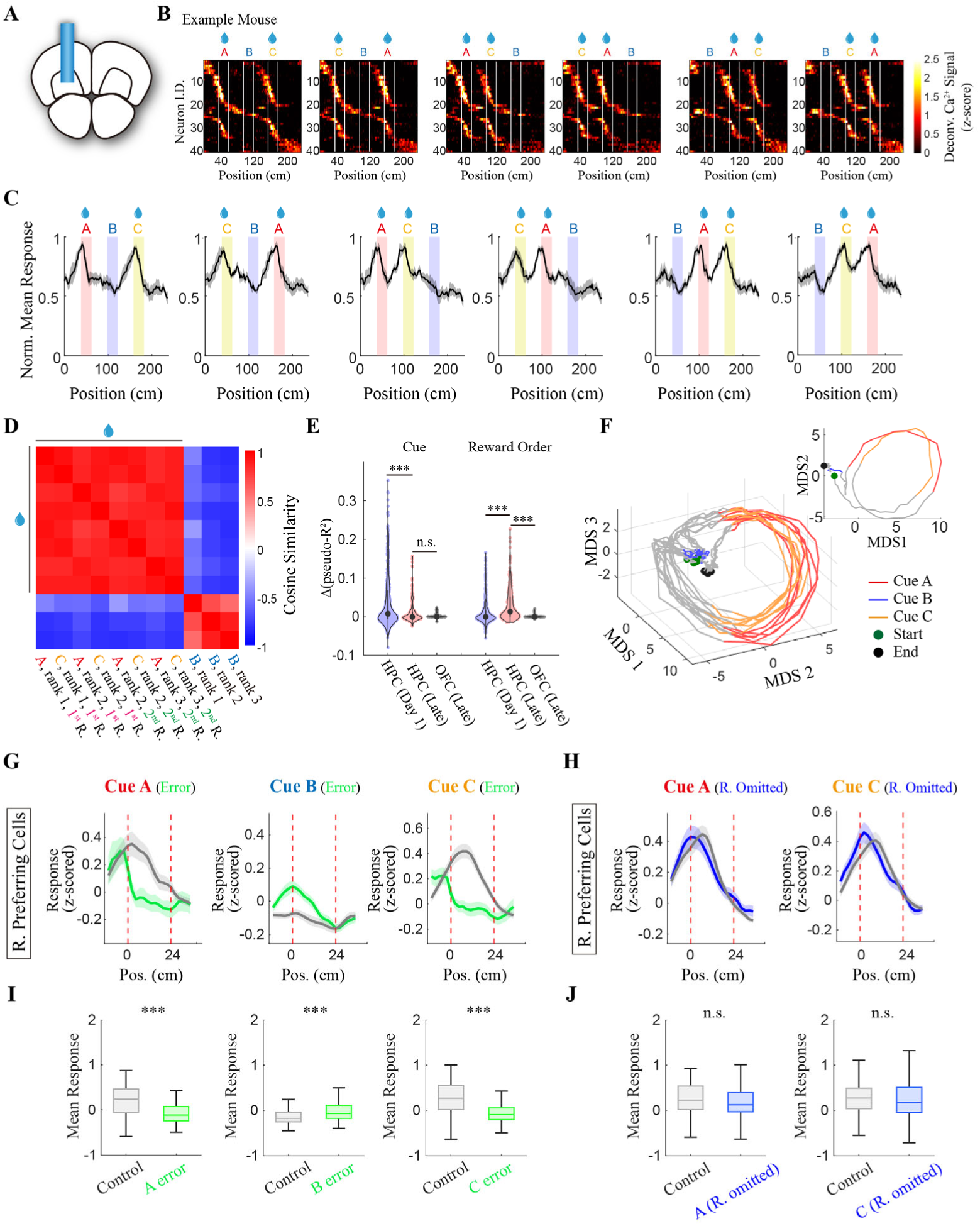
OFC representations were dominated by reward expectation responses but not task states. **(A)** Schematic of grin lens implantation in the orbitofrontal cortex. **(B)** z-scored neural activities in one example mouse. Cells were sorted based on their activity peaks in the ABC trial type (the first column). White lines indicate cue positions. **(C)** Averaged response in the orbitofrontal cortex on late sessions. Curves and shaded areas represent mean ± s.e.m., respectively; n=5 mice. **(D)** Representation similarity matrix (RSM) of the orbitofrontal cortex on late sessions. Cells from all mice were pooled together. Population vectors were calculated as averaged responses ±20 cm around the cue onset for each condition. **(E)** Δ(pseudo-R^2^) for rewarded cue and reward order. Purple, pink, and gray dots represent HPC day 1 session (n=477 cells), HPC late session (n=231 cells), and OFC late session (n=73 cells). Only reward preferring cells were included in analysis. For direct comparison with OFC, responses of individual cells in HPC were re-calculated by averaging activities in the interval ±20 cm around the cue onset. Δ(pseudo-R^2^)(cue), HPC day 1 vs. HPC late: *P* = 1.854 × 10^-4^; Δ(pseudo-R^2^)(cue), HPC late vs. OFC late: *P* = 0.2884; Δ(pseudo-R^2^)(reward order), HPC day 1 vs. HPC late: *P* = 1.036 × 10^-16^; Δ(pseudo-R^2^)(reward order), HPC late vs. OFC late: *P* = 1.768 × 10^-10^; two-sided Wilcoxon rank sum test. **(F)** Neural activity trajectories in 3-dimensional MDS space. Six trial types were plotted together. Cells were pooled from all mice. Inset: two-dimensional MDS projection of the trajectory for the C–A–B trial sequence. **(G)** Responses of reward preferring cells to different cues in control (grey) and error (green) trials. Curves and shaded areas represent mean ± s.e.m., respectively. **(H)** Responses of reward preferring cells to different cues in control (grey) and catch (blue) trials. Curves and shaded areas represent mean ± s.e.m., respectively. **(I)** Quantification of reward preferring cells in (G). Averaged activities at cue zone in each cell were used for statistical comparison. *P* = 1.524 × 10^-4^, response at A in control trials vs. response at A in A-error trials; n=55 cells; *P* = 8.804 × 10^-4^, response at B in control trials vs. response at B in B-error trials; n=82 cells; *P* = 2.548 × 10^-11^, response at C in control trials vs. response at C in C-error trials; n=124 cells; two-sided Wilcoxon signed-rank test. **(J)** Quantification of reward preferring cells in (H). Averaged activities at cue zone in each cell were used for statistical comparison. *P* = 0.2079 response at A in control trials vs. response at A in A-catch trials; n = 124 cells; *P* = 0.2825, response at C in control trials vs. response at C in C-catch trials; n = 124 cells; two-sided Wilcoxon signed-rank test.

Because OFC activity was predominantly associated with rewards, we next leveraged error trials and catch trials to dissociate different reward-related factors. The observation that many OFC neurons showed activation preceding reward delivery makes us hypothesize that their activities were associated with reward expectation (or licking) rather than the reward itself. Consistent with this hypothesis, responses of reward preferring cells markedly reduced when mice did not lick at rewarded cues, whereas ectopically enhanced when animals licked at the non-rewarded cue (Fig. 7G,I). Analysis of catch trial activity further supported our hypothesis. No significant response change was detected following reward omission (Fig. 7H,J), suggesting that OFC neurons did not primarily encode outcome (reward consumption).

Together, these results highlight distinct roles for hippocampus and OFC in representing our structured cue-guided navigation task. Hippocampal representations captured reward-sequence structure while abstracting across the identities of the two rewarded cues. Through conjunctive coding of sequence rank and reward order, hippocampal activity distinguished latent states with different transition structures. In contrast, OFC formed a more abstract representation, grouping task segments into different categories based on their behavioral relevance (rewarded vs. non-rewarded) without preserving detailed information about state transitions.

## Discussions

In this study, we developed a structured cue-guided navigation paradigm and performed longitudinal two-photon imaging to track hippocampal neural activity throughout learning. Our findings lead to four major conclusions: (1) the hippocampus exhibits disentangled representations of distinct task variables; (2) abstract hippocampal representations of reward sequences emerge through learning in most animals; (3) abstraction of hippocampal representations predicts structure-based behavioral strategies; and (4) in our task, orbitofrontal cortex (OFC) activity primarily encodes reward expectation rather than task structures.

Our findings suggest that the geometry of hippocampal representations predicts the behavioral strategies spontaneously adopted by individual animals. Two notable features of our behavioral paradigm design distinguish it from most previous studies on schema learning. First, our task did not explicitly require animals to learn abstract structures. In principle, perfect task performance could be achieved solely through immediate visual information, as observed in a subset of mice (the cue-encoding group). Nevertheless, structure-based strategy emerged spontaneously in most mice. This finding raises the possibility that the hippocampus continuously constructs internal models of environmental structure even in the absence of explicit behavioral demand to utilize the structural information, consistent with classical studies of latent learning^36^ and a recent report demonstrating predictive representations in the human hippocampus during unconscious states^37^. Second, unlike most schema-learning paradigms, our task did not expose animals to multiple related problems. Despite training on only a single task structure, most animals developed cue-invariant hippocampal maps and adopted structure-based behavioral strategies. These findings highlight a remarkable property of biological intelligence: a strong intrinsic tendency to extract abstract structure from experience.

The hippocampus has long been considered a central substrate for cognitive map. However, the precise content encoded within hippocampal cognitive maps remains under investigation. Early studies in rodents primarily emphasized spatial coding, owing to the discovery of place cells during navigation. Subsequent work has substantially expanded this view. The hippocampus has been shown to encode a wide range of non-spatial environmental variables, including time ^38,39^, reward^40–42^, and the location of social counterparts^43,44^, as well as behavioral variables such as speed^45,46^ and choice^32,47,48^. Furthermore, hippocampal activity has been reported to encode abstract variables relevant to cognitive tasks, including accumulated evidence^19^ and lap numbers^49^. Our findings indicate that learning can dynamically shape information encoded in the hippocampal map, which depends on the animal’s behavioral strategy. Specifically, the geometry of hippocampal population representations predicts whether an animal adopts a sensory-based or structure-based problem-solving strategy.

Differences in hippocampal representations can explain individual behavioral variability in our experiments. In structured cue-guided navigation paradigm, two groups of animals naturally emerged during learning: mice with abstract hippocampal representations and mice with sensory-oriented representations. Notably, these groups arose despite experiencing identical task rules. The factors that bias animals toward sensory-based versus structure-based learning remain unclear. Although we cannot rule out contributions from genetic variability, our results suggest an important role for experience. Individual animals do not appear to be irreversibly committed to a particular strategy. Training across multiple tasks that share common underlying structure promoted the transition from sensory learners to structure learners, consistent with extensive evidence that broader training distributions facilitate generalization. Thus, experience prior to experiments may establish inductive biases that influence subsequent behavioral strategies.

Our study further reveals an abstraction process within hippocampal representations during learning. Consistent with previous work^29^, hippocampal neurons exhibited mixed selectivity for multiple task-relevant variables in the structured cue-guided navigation task, including sensory cues, sequence rank, and reward order. Increasing evidence suggests that representational geometry plays a critical role in cognitive functions such as generalization and classification^50^. One mechanism that supports generalization while preserving detailed sensory information is the formation of disentangled representations, in which distinct task variables occupy orthogonal coding dimensions. Such representational structures have been reported in both humans^13^ and non-human primates^14^ during inference of reward contingency contexts. We observed a similar disentangled representational geometry in mice following pre-training, where sensory cue and sequence-rank information occupied nearly orthogonal dimensions. Interestingly, introduction of a novel rewarded cue preserved rather than disrupted this organization. More strikingly, our findings suggest that prolonged learning may induce a further stage of abstraction. Over training, cue-related coding dimensions collapsed whereas reward-order representations expanded. It should be noted that, in our task, the number of passed rewards correlated with the number of available rewards in future until the end of one trial. Future studies with changing reward numbers are required to distinguish the encoding of consumed rewards versus reward availability. Nonetheless, we show that generalization of hippocampal map does not stop at the disentangled geometry, but continues to shrink the cue-coding dimension through learning. Whether similar process occurs in primate hippocampus under some conditions remains unknown.

The abstraction process reflects changes in the animal’s internal representation of task structure rather than alterations in overt behavior or task settings. Recent studies have illustrated that the hippocampus can encode action plans^51^ and task rules, i.e. reward contingencies^20^. By incorporating a pre-training stage, we captured an early learning phase in which animals had already achieved high behavioral performance, yet abstraction within hippocampal representations had not fully emerged. With continued training, hippocampal representations underwent substantial reorganization despite stable task rules and largely unchanged behavioral performance. This phenomenon resembles delayed generalization observed in artificial neural networks, including the recently described *grokking* phenomenon^52^, in which generalization emerges only after training performance has already saturated. Our findings raise the possibility that biological and artificial neural systems may share common computational principles, such as regularization mechanisms, that promote delayed emergence of generalized representations.

Our findings also provide an example of latent-state map formation within hippocampal representations. The concept of hidden-state inference in the hippocampus has arisen from observations of splitter cells^47,48,53,54^. In spatial alternation tasks, splitter cells display distinct activity patterns at the same location depending on an animal’s future behavioral choice. Subsequent works extended this concept to more complex cognitive tasks, demonstrating how hippocampal circuits can divide identical sensory inputs into distinct latent states^20,21^. Consistent with these findings, we observed that hippocampal representations separated rank-2 rewards into distinct states depending on preceding trajectories. Importantly, however, not all task stages underwent equivalent splitting. Rank-3 rewards showed substantially weaker separation compared with rank-2 rewards. This observation suggests that the hippocampus does not indiscriminately partition every state associated with distinguishable trajectories. Instead, hippocampal representations appear to form a parsimonious Markov graph in which the minimal number of hidden states are generated such that each state sufficiently predicts future state transitions. This finding aligns closely with the theory proposing that the hippocampus constructs predictive cognitive maps through successor representations^24^, which encode time-discounted future occupancy across states.

Representations of both physical and abstract variables are distributed across multiple brain regions. Both the hippocampus and OFC have been implicated in constructing cognitive maps. Silencing either area significantly impaired model-based learning^55^. Interestingly, abstraction of cognitive maps similar to that observed here has previously been reported in the rat OFC during an olfactory sequence learning task^2^. In contrast, our data suggest that OFC activity in this task is dominated by reward-related signals rather than task-state representations. Several differences between behavioral paradigms may account for this discrepancy. First, our task relied on visual stimuli, whereas previous studies used olfactory cues. In rodents, OFC maintains strong reciprocal connectivity with olfactory regions such as piriform cortex but receives comparatively sparse direct inputs from the visual cortex. Given differences in anatomical connectivity across sensory modalities, cognitive representations may exhibit modality-dependent biases. Second, our task did not require animals to explicitly maintain previous information through working memory. The ability to acquire task representations through latent learning may rely more strongly on the hippocampus. Third, our mice were trained using a single cue set, whereas prior studies involved multiple related problems. OFC abstraction may require broader experience for the generalized map to emerge. Future work should investigate how hippocampus, OFC, and other cognitive regions interact to establish coordinated representations supporting learning and decision-making.

## Acknowledgements

We thank Dr. Jingfeng Zhou for generous sharing of mouse strains and insightful discussions. We thank Dr. Quan Wen, Dr. Tianming Yang, and Dr. Jianing Yu for insightful discussions. We thank all Zhao lab members for technical help and insightful discussions. This work is supported by the National Key R&D Program of China, Project Number 2025YFA1016700, National Natural Science Foundation of China General Grant, Project Number 04130102323, and Tsinghua-Peking Center for Life Sciences.

## Methods

### Animal Surgery

#### Head bar Implant

Adult (P60-120) male mice (C57BL/6J, n = 6, from Charles River Laboratories) were anesthetized with isoflurane (3–4% for induction and 1% for maintenance) and mounted onto a stereotaxic apparatus with ear bars. After scalp removal, coordinates for craniotomies were determined using a micromanipulator and marked with ink. Bilateral dorsal CA1 sites (2.2 mm posterior and 2.2 mm lateral to bregma) were identified and marked. The skull surface was then cleaned and roughened using a dental drill and scalpels to improve adhesion of the head bar. The skull surface was first coated with glue, after which a customized titanium head bar was affixed using dental cement (Super-Bond, Sun Medical). The central opening of the head bar was sealed with silicone elastomer (Kwik-Cast, World Precision Instruments). All animal experiment protocols are approved by the Institutional Animal Care and Use Committee (IACUC) at Tsinghua University.

#### Muscimol Injection

Muscimol injections were performed following the pre-training task. After animals reached a performance criterion of >90% success rates for 3–4 consecutive days, sham control and muscimol inactivation experiments were conducted sequentially. To control for the effects of surgery and anesthesia, sham experiments involved craniotomy only, with anesthesia maintained for approximately 1 h to match the typical surgery length of muscimol injection. Mice were then allowed to recover in their home cages for 1 h before behavioral testing. On the following day, mice were anesthetized again for muscimol injections. BODIPY TMR-X-conjugated muscimol (3.2 mM; Invitrogen) was bilaterally injected into the dorsal hippocampus using a Nanoject Ⅲ injector (Drummond Scientific). For each hemisphere, injections were delivered at two depths (1.5 mm and 1.9 mm below the pia), with 40 nL injected at each depth. Animals were allowed to recover for 1 h before behavioral testing.

#### Cannula Implantation in Hippocampus

Adult male mice (Thy1-GCaMP6f, GP 5.17 line, n = 4; Thy1-GCaMP6s, GP 4.3 line, n = 8; all from the Jackson Laboratory) were used for hippocampal cannula implantation. Anesthesia and surgical preparation were performed as described above. After scalp removal and clearance of connective tissue, a circular craniotomy (1.5 mm radius) centered at AP −2.2 mm and ML 2.2 mm was made above the right dorsal hippocampus. After removal of the dura, cortical tissue and the transverse fibers of the corpus callosum were aspirated using a flat-ended needle connected to a vacuum pump. The appearance of oblique fibers in the lower-left region of the surgical field indicated exposure above the hippocampus. After complete hemostasis, a custom titanium cannula (1.4 mm in height) pre-assembled with a glass coverslip (thickness: 0.15 mm), was implanted above the hippocampus. A customized titanium head bar was subsequently secured to the skull using dental cement, as described previously. Following surgery, mice were allowed to recover for at least 1 week before further experimental procedures.

#### GRIN Lens Implantation in Orbitofrontal Cortex

Adult male mice (Thy1-GCaMP6f, GP 5.17 line, n = 4; Jackson Laboratory) were used for GRIN lens implantation in the orbitofrontal cortex. Anesthesia and surgical preparation were performed as described above. After scalp removal and clearance of connective tissue, a circular craniotomy (1.0 mm diameter) centered at AP +2.3 mm and ML +1.45 mm was made above the right orbitofrontal cortex. After superficial blood vessels were removed and bleeding was controlled, a GRIN lens (1 mm diameter, 4.39 mm length, 250 μm working distance in water; GRINTECH GmbH, Germany) was slowly lowered through the cortical tissue to a target depth of 2.0 mm below the dural surface. The lens was secured on the skull with dental cement, and a customized titanium head bar was subsequently affixed to the skull as described above. Mice were allowed to recover for at least 2 weeks before further experimental procedures.

### Histology

For post hoc verification of muscimol injection sites and imaging window placements, mice were transcardially perfused with PBS followed by 4% paraformaldehyde (PFA). Brains were post-fixed overnight, sectioned coronally at 100 μm using a vibratome (Leica Microsystems), and mounted onto glass slides. Sections were counterstained with DAPI (2.5 mM in glycerol mounting medium), coverslipped, and sealed with nail polish. Images were acquired using an Olympus VS200 slide scanner.

### Behavioral Training

#### Virtual Reality Apparatus and Behavioral Tasks

Behavioral training was conducted in a virtual reality (VR) system in which head-fixed mice navigated on an air-supported floating ball positioned in front of surrounding VR screens. The hardware design of the VR system was adopted from previous reports ^56,57^, whereas the control software was developed using Unity. Since our behavioral tasks only required running in linear tracks, we fixed the animal’s view angle to be straight and locked its lateral position at the center of the track. A lickport positioned in front of the animal delivered 10% sucrose solution as reward, and licking was monitored using infrared lick sensors that detected interruptions of the infrared beam by tongue protrusions. Micro-processors (Arduino Uno) were used to record signals from lick sensors and control the roller pump for reward delivery.

Water restriction, with 1.0ml per day, started 3-10 days before the training began. The water schedule continued throughout the entire experiment.

To facilitate training, a curriculum of three training stages was implemented. First, before imaging sessions began, mice were habituated to head fixation and locomotion on the spherical treadmill for 3–5 days with the VR screens turned off until they could run naturally in the setup. No reward was delivered to the animal during this stage. Second, mice were exposed to VR environments with a higher gain between locomotion and virtual movement to facilitate adaptation to the VR navigation. Once mice exhibited stable running, the gain was gradually reduced so that virtual movement closely matched the actual locomotion speed. Rewards were automatically delivered when mice entered the rewarded cue zone. Third, after mice learned to actively lick for reward acquisition, the task was switched to the ‘operant conditioning’ mode, in which mice were required to lick to trigger the reward delivery.

#### Pre-Training Task

In the pre-training task, mice learned the association between two visual cues and reward outcomes. Cue A was consistently associated with reward delivery, whereas cue B was not rewarded. In each trial, Cue A and cue B were randomly assigned to two of three possible slots distributed sequentially along a 238-cm linear virtual track, leaving the remaining slot empty. This arrangement generated six possible trial types, which were presented randomly trial-by-trial. The three cue slots were located at 38.8–62.8 cm, 98.8–122.8 cm, and 158.8–182.8 cm along the track. At the end of each trial, a blue visual block was presented, followed by a 2–4 s black-screen inter-trial interval before the onset of the next trial. Mice typically completed ∼140–200 trials per day and received approximately 1 mL of sucrose solution daily.

#### Full Task

After mice reached the >80% success rate in the pre-training task (A-B-Empty), a third novel visual cue, cue C, was introduced into the previously empty slot. Similar to cue A, cue C was associated with reward. During the transition session, mice first performed ∼60 trials in the pre-training task and were then switched to the full task within the same session. Training in the full task continued for several additional sessions.

#### Reward Sequence Completion Task

For the reward sequence completion task, the visual cue presented at the third slot was not rendered while the associated reward contingency remained unchanged. To minimize potential confusion in task-rule interpretation, mice were first trained with intact visual cues for ∼20–30 trials. Subsequently, probe trials with the third cue masked were added, which randomly interleaved with trials containing full visual cues. Masked and intact trials were presented at a ratio of 2:1.

#### Full Task with Novel Rewarded Cues

In a subset of mice, we trained them with a different set of rewarded cues to promote generalization. In this task, the two originally rewarded visual cues, A and C, were replaced with two novel cues, E and F, respectively. Cue B remained unchanged and continued to be unrewarded. During the transition session, mice first performed ∼60 trials in the first full task(A-B-C) before being switched to the second full task(E-B-F) within the same session.

### 2- Photon Calcium Imaging

Two-photon calcium imaging was performed using an Olympus microscope equipped with a 25× water-immersion objective (XLPLN25XSVMP2, NA 1.0, Olympus) and controlled by FV31S-SW software. Images were acquired at 30 Hz using resonance scanning with a frame size of 512 × 512 pixels during behavioral sessions lasting approximately 60 min. Two consecutive frames were averaged first before image processing. In addition to the GCaMP fluorescence channel, a second acquisition channel recorded synchronization TTL pulses generated by the behavioral program.

A template image was generated using maximum projection of images acquired in the first session. In later sessions, the field of view and focal plane were carefully adjusted to align with blood vessel landmarks and cell patterns in the template image before imaging started.

## Data Analysis

Data analysis was implemented using customized MATLAB codes.

### Extraction of Deconvolved Ca^2+^ Activity Using Suite2P and Alignment with Behavioral Data

ROI registration, selection, and raw fluorescence trace extraction were performed using Suite2p^58^ with default parameters. Deconvolved calcium activity was inferred from neuropil-corrected fluorescence traces using the OASIS deconvolution algorithm implemented in Suite2p (decay time constant: 0.7s for GCaMP6f and 1.25s for GCaMP6s). Synchronization pulses recorded in the second imaging channel were used to align imaging and behavioral data. Timestamps of synchronization pulses detected in the imaging data were matched to corresponding pulses recorded by the behavioral program, and alignment was corrected at each pulse to compensate for temporal drift between clocks in the two systems.

### Cell Registration Across Days

Following automatic ROI identification in Suite2p, ROIs were aligned across days using CellReg^33^. Registration was first performed based on spatial correlation, and the accuracy of cross-day registration was then validated using centroid distance (Supplementary Fig. 13). Cells included in the final analysis were those that exhibited both high spatial correlation and low centroid distance.

### Selection of Highly Responsive Cells

To improve reliability of quantitative analyses, including selectivity indices, encoding models, and representational similarity analyses, most analyses were performed with highly responsive neurons. One exception was Fig. 3, in which all recorded neurons were included to reproduce the key finding of rewarded-cue coding collapse during learning. Another exception was supplementary Fig. 4, in which we showed increased cue and rank selectivity through learning.

For each neuron, activity was first z-scored across the entire imaging session. Median z-scored activity across trials was then computed separately for each of the six trial types. The largest value among these six median responses was defined as the neuron’s peak response. Neurons with peak responses exceeding 2 were classified as highly responsive.

Taking the maximum response across trial types allowed identification of neurons exhibiting trial-type-specific tuning. This procedure typically selected approximately 10–30% of imaged neurons as highly responsive.

### Selectivity Indices for Cues and Ranks

Selectivity indices (SI) for cue and rank were quantified based on deconvolved calcium activity across different cue identities and cue ranks in the pre-training task. Because hippocampal neurons exhibited cue-anchored activity sequences (Supplementary Fig. 3), the virtual track was divided into three cue-anchored slots, defined as the spatial interval from the onset of one cue to the onset of the next cue.

Cue SI was computed by comparing neuronal responses to cue A and cue B:

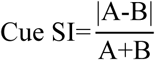

where A and B represent the peak z-scored deconvolved calcium activity of the neuron in trials containing cue A or cue B, respectively.

Rank SI was calculated by comparing neuronal responses across the three ranks, adopted from previous studies ^29^:

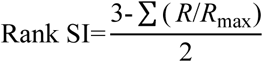

where R denotes the peak neuronal response at each rank and R_max_ represents the maximum value of R across three ranks.

### Geometric Analysis

To characterize representational geometry, we applied multidimensional scaling (MDS; mdscale in MATLAB) to project high-dimensional neural population activity into a three-dimensional space. Angles between coding axes were quantified within the resulting 3D-MDS space. Using the first three dimensions from a four-dimensional MDS projection yielded qualitatively similar results (data not shown).

Representational similarity analyses were performed using raw z-scored neural activity without dimensionality reduction.

For analyses shown in Fig. 1J and Fig. 3M,N, shuffled distributions were generated using 1000 random permutations of neuron identities before dimensionality reduction.

#### Cue Preference Index

For each neuron, mean activity was first calculated within each slot unit (defined as the interval from cue onset to the end of the subsequent gap period) for each trial type. Responses to individual cues were then obtained by averaging across all six trial types.

The preference index for a given cue *X* was defined as:

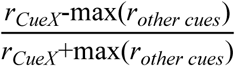

where *r_CueX_*denotes the mean response to cue *X*, and *r_other cues_*denotes responses to the remaining two cues.

Therefore, the cue preference index ranges from -1 to 1, with larger values indicating to preferred activation by cue X.

Cue-selective cells were defined as cells with preference index larger than 0.33 for a given cue (i.e. the response to the give cue is at least twice as large as others).

Similarly, the reward preference index was defined as:

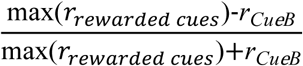

Reward-selective cells were defined as cells with preference index larger than 0.33.

#### Cross-Condition Generalization Performance (CCGP)

CCGP was quantified to assess whether neural representations generalized across conditions.

For cue decoding, a linear support vector machine (SVM; one-versus-rest approach) was trained to classify the three cues using neural responses at rank 1. Decoder performance was subsequently tested on neural responses at rank 2 and rank 3, yielding two cross-condition decoding accuracies. This procedure was then repeated by training decoders separately at rank 2 and rank 3. CCGP for cue identity was defined as the average of all six cross-condition decoding accuracies.

CCGP for rank was quantified analogously by training decoders on one cue condition and testing on the remaining cue conditions.

Shuffled controls were generated using 500 random permutations of neuron identities.

#### GLM Model

Generalized linear models (GLMs) were trained and evaluated using five-fold cross-validation. A logarithmic link function was used assuming a Poisson distribution of responses.

For each iteration, the dataset was randomly divided into five subsets, with four subsets used for training and one for testing. Model performance was quantified using pseudo-*R*^2^, computed from the difference between the log-likelihood ratio (LLR) of the fitted model and that of a null model containing only a constant term (*β*_0_).

This procedure was repeated across all five folds, and the average pseudo-*R*^2^ value was used to quantify model performance.

To estimate the contribution of individual variables, pseudo-*R*^2^ values obtained from reduced models lacking a given predictor were subtracted from that of the full model.

Mean activity within slot units was used as the response variable. Predictor variables for individual models are described in the corresponding Results sections and figure legends.

#### Analysis of Error and Catch Trials

Because error and catch trials occurred relatively infrequently, the distribution of trial types within these subsets was often unequal. To avoid biases arising from unbalanced trial-type composition, responses in correct trials were matched to the distribution of error or catch trials.

For each neuron, the control response was calculated as the weighted average of responses in correct trials across the six trial types, where weights were determined by the number of error or catch trials in each trial type:

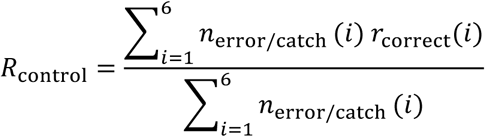

where *n*_error/catch_(*i*) denotes the number of error or catch trials of trial type *i*, and *r*_correct_(*i*) denotes the mean neuronal response during correct trials of the same trial type.

#### Mutual Information

Mutual information analyses were performed to quantify the relationship between neuronal firing rate and four task-related variables: cue identity, rank, reward, and reward order. The detailed categorization of these variables is illustrated in Supplementary Fig. 14E. Mutual information between neuronal firing rate (R) and task variable (X) was calculated as:

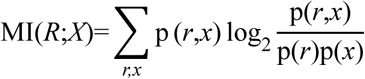

where p(*r*,*x*) denotes the joint probability distribution of firing rate and task variable, and p(*r*) and p(*x*) represent their marginal probability distributions.

Because neuronal activities exhibited a heavy-tailed distribution (Supplementary Fig. 14A-D), activities were discretized using unequal-width binning to ensure adequate sampling across activity levels. For each neuron, mutual information values were computed separately for cue identity, rank, reward, and reward order. Neurons were subsequently classified into different functional categories based on the dominant mutual information component together with the position of their mean response peaks, as illustrated in Supplementary Fig. 14F. Numbers of categories for cue, rank and reward are 3, 5, and 2, respectively. In the classification, the cut-off mutual information for reward was set as 0.1, and that for cue and rank were set as 0.16 (0.1 × *log*3/*log*2) and 0.23 (0.1 × *log*5/*log*2) to balance different numbers of categories.

#### SVM Decoding of Past Trajectories

Linear SVM classifiers were trained to decode preceding trajectories using neural responses recorded during rank-2 or rank-3 states across trials. Decoder performance was quantified using five-fold cross-validation.

Null distributions were generated using 500 random permutations of neuron identities.

#### Bayesian Decoding

Because the goal of Bayesian decoding was to characterize information related to trial type and position contained within neural populations, uniform prior probabilities were assumed. Under this assumption, Bayesian decoding was equivalent to maximum-likelihood estimation.

Due to strongly skewed distributions of deconvolved calcium activity, non-uniform activity binning was used. For each neuron, the highest activity bin was defined as the range spanning 50–100% of maximal activity, whereas the remaining range (0–50%) was equally divided into nine bins, yielding ten bins in total.

Activity probabilities within each bin were estimated empirically from training data to generate likelihood functions. No assumptions regarding underlying response distributions (e.g., Poisson distributions) were imposed to fit the likelihood function.

Posterior probabilities across all trial types and positions were calculated as the product of likelihood probabilities across neurons. A leave-one-trial-out approach was used: one trial served as test data while all remaining trials were used for training. This procedure was repeated across all trials, and posterior distributions were averaged across iterations to generate the final decoding estimates.

## Computational Modeling

All models were implemented in Python.

### Hierarchical Dirichlet Process Hidden Markov Model (HDP-HMM)

#### Model Architecture

We modeled latent task states using a Hierarchical Dirichlet Process Hidden Markov Model (HDP-HMM), a nonparametric extension of the Hidden Markov Model that allows the number of hidden states to be inferred from data. The model is formulated using the Chinese Restaurant Franchise (CRF) representation of the Hierarchical Dirichlet Process. A vector of global state popularity, ***β***, was sampled from a Griffiths–Engen–McCloskey (GEM) distribution:

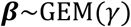

where γ is a concentration hyperparameter that controls the complexity of the state space. Larger γ allows more hidden states to be created.

For each hidden state *i*, transition probabilities were sampled from a Dirichlet Process (DP) centered on the global state distribution Q:

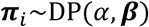

where α is a hyperparameter which determines how similar each ***π***_i_ is.

The emission from the hidden state *i* to different observations is defined as ф_i_ = *P*(***X***|*z* = *i*). Gibbs sampling was used to optimize Q, ***π***, and ф.

In each iteration, a sequence of hidden states, *z*_t=1:T_, is first sampled from the observation sequence *x*_t=1:T_ following the Bayesian equation:

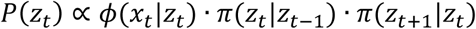

Given this newly sampled *z*_t=1:T_, Q was updated using the CRF process:

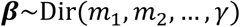

where Dir means Dirichlet distribution and *m*_j_ denotes the number of ‘tables’ in CRF that choose the state *j*. ***π*** was then updated based on the new Q and *z*_t=1:T_:

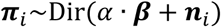

where ***n***_i_ denotes numbers of transitions from the state *i* to other states in *z*_t=1:T_.

Finally, the emission matrix was updated as:

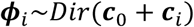

where ***c***_0_ is the prior pseudo-count and ***c***_i_ denotes counts of all observations assigned to the state *i* in the *z*_t=1:T_, *x*_t=1:T_ sequences. In all simulations, ***c***_0_ = [1,1, …].

Throughout optimization, γ was a fixed hyperparameter whereas α was optimized using Escobar-West algorithm.

Chinese restaurant process was previously proposed to model how the hippocampus map different environments (global remapping)^59^. We here extend similar concept to the map of different states within one environment.

***Performance evaluation*.**

To evaluate latent-state representations, we generated 5,000 simulated trials. Each trial consisted of a ‘Start’ observation followed by two ‘Reward’ observations and one ‘Non-reward’ observation presented in random order. The resulting dataset contained 20,000 observations. Model fitting was performed using batched replay. During each Gibbs sampling iteration, a contiguous sequence of 2,000 observations was randomly sampled from the full-length sequence and used for parameter updates. The maximum number of hidden states was truncated at *K*_max_ = 30.

To investigate the effect of regularization, the concentration parameter γ was varied across conditions. Specifically, γ = 3 was used in the regularized condition, whereas γ = 100 was used in the no-regularization condition. 400 iterations were performed for all conditions.

### Clone-Structure Cognitive Graph (CSCG)

Training of CSCG followed the original publication ^34^. Details of the model are described below.

#### Model Architecture

CSCG was implemented as a variant of the hidden Markov model (HMM) in which observed data were assumed to be generated from a hidden Markov process. In a standard HMM, the joint distribution over observations *x*_1_,*x*_2_,⋯,*x_N_* and hidden states *z*_1_,*z*_2_,⋯,*z_N_* factorizes as

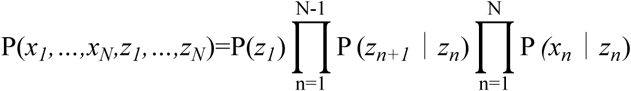

The CSCG was implements with a structured emission matrix, where each hidden state deterministically emits a single observation, and multiple hidden states (clones) could map to the same observation. Formally, the emission probability is defined as

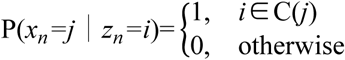

The probability of a sequence is obtained by marginalizing over the hidden states

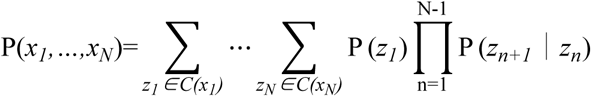

#### Task details

The task was represented as a long sequence of discrete symbols, reflecting progress along a virtual linear track with three reward zones. The reward schedule across the three zones defined three trial types: RRN (reward, reward, non-reward), RNR (reward, non-reward, reward), and NRR (non-reward, reward, reward). Each trial was encoded as a sequence of symbols: a start symbol “3”, a number of interval symbols “0”, three reward zones where a reward was indicated by “1” and a non-reward by “2”, an end symbol “4”, and an inter-trial interval symbol “5”. For example, an RNR trial took the form [3, 0, 0, 0, 1, 1, 0, 0, 2, 2, 0, 0, 1, 1, 0, 0, 0, 4, 5, 5]. All symbols were one-hot encoded. Training sequences of 1000 time steps were generated by concatenating trials chosen uniformly at random from three types.

A CSCG model with 100 clones per observation was initialized with random transition probabilities (each entry drawn from a uniform distribution and then row-normalized) and a deterministic emission matrix (each clone assigned to exactly one observation). The number of clones was kept constant across all observations. The transition matrix was learned using the Baum-Welch algorithm to maximize the likelihood of the training sequence. Training proceeded for 2000 iterations. Regularization was implemented via a pseudocount parameter *κ* added to the accumulated transition counts before normalization. For the non-regularization condition, *κ* was set to 1e-10; for the strong-regularization condition, *κ* was set to 1e-6.

### Performance evaluation

The average prediction probability per time step was computed on a held-out test sequence of 1,000 steps. The most likely state sequence was decoded using the Viterbi algorithm, and the transition graph between clones was visualized.

### Recurrent Neural Network (RNN)

#### Model architecture

A vanilla RNN was implemented with an input layer, a hidden layer (100 units, ReLU activation), a dropout layer (rate 0.6 in the strong-regularization condition; omitted otherwise), and an output layer. The hidden state dynamics are defined by

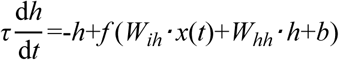

where *x*(*t*) was the input vector, the hidden state h, and learnable parameters *W_ih_*, *W_hh_*, and *b* (biases included as PyTorch default). The equation was discretized using the forward Euler method with Δ*t*/*τ*=1.

#### Task details

Inputs were one-hot vectors for cues {S, A, B, C, E}; outputs were one-hot vectors for reward (R) or no-reward (N). Each trial comprised five steps (S, three cues in random order, E). The network received a long concatenation of trials sampled uniformly from the six possible orderings. At each time step, predictions were made for both the current and the next reward. The RNN was trained using the Adam optimizer (learning rate = 1e-3) with a batch size of 64, a cross-entropy loss, and backpropagation through time (BPTT) over 3,000 epochs. The hidden layer comprised 100 units with ReLU activation, followed by a dropout layer with a rate of 0.6 in the strong-regularization condition.

#### Performance evaluation

A test sequence of 1000 trials was generated. Hidden states were averaged per trial type and time step, yielding a 6 (trial types)×5 (time steps)×100 (neurons) matrix of mean activities. Individual neuron activity was visualized with line plots. Pairwise cosine similarities were computed between activity patterns of different states, and classical multi-dimensional scaling (MDS) was applied to the cosine distance matrix (1–cosine similarity) for projection into three dimensions, with resulting trajectories visualized as 3D arrows showing state transitions for each trial type.

#### Tolman Eichenbaum Machine (TEM)

Construction and training of TEM followed the original publication ^35^. Details of the model are described below.

### Model architecture

At each time step, TEM receives a one-hot encoded sensory observation *x_t_* and an action *a_t_* that caused a state transition. Its objective is to predict the next sensory observation *x_t_*_+1_ given the history of observations and actions. TEM comprises three latent variables: a structural variable *g*, a sensory variable *x_t_*, and a conjunctive variable *p* defined as

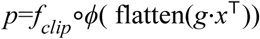

where *f_clip_* bounded the value to [-1, 1] and ϕ denoted LeakyReLU with slope 0.01.

TEM is implemented within a variational inference framework, consisting of a generative model and an inference model. The generative model defines the probabilistic relationship *p*(*x*_≤*T*_,*g*_≤*T*_,*p*_≤*T*_∣*a*_≤*T*_), and the inference model approximates the posterior *q*(*g*_≤*T*_,*p*_≤*T*_∣*x*_≤*T*_,*a*_≤*T*_). Specifically, at each time step the generative model receives the action *a_t_* and the previous memory matrix 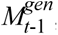, and proceeded as follows:

1. path integration based on *g_t_*_-1_ and *a_t_* yielded

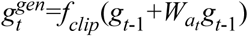

where *W_at_* was an action-dependent transition matrix.
2. using 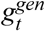 as a query, the attractor network retrieved a conjunctive memory and decoded it to obtain a sensory prediction

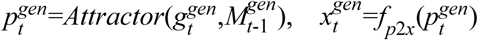

Here the *Attractor*(⋅) is defined by the iterative dynamics h^(*r*)^=*f*(*κ*h^(*r*-1)^+*M*h^(*r*-1)^) for *r*=1,…,5, starting from h^(0)^=flatten(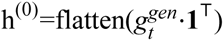⋅**1**^T^) (or, in the inference model, flatten(**1**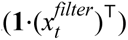^T^), and the final h^(5)^ gives the retrieved conjunctive memory vector.

The inference model receives the generative model’s 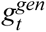 together with the sensory input *x_t_*, and performed the following steps:

1. compression and exponential temporal filtering of the sensory input

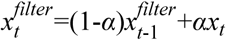

where *α* is a learnable filtering parameter.
2. using the filtered sensation as a query, the inference model’s memory matrix *M_t_*_-1_ retrieves a memory

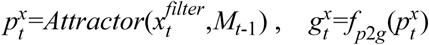
3. Bayesian fusion of 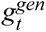 and *g_x_* produce the final structural estimate

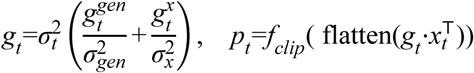

where *σ_t_* =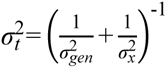 . The variances 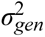 and 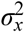 were outputs of learned networks that estimate the uncertainty of the path integration and sensory retrieval, respectively.

The model is trained to minimize the discrepancy between the generative and inference models, with the total loss given by *L*=*L_x_*+*L_p_*+*L_g_*. All network weights except the memory matrices are shared across environments and updates slowly via truncated backpropagation through time. Memory matrices are reset to zero at the beginning of each environment and updated rapidly using Hebbian rules

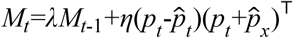

where 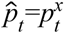 (inference model) or 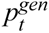 (generative model).

#### Task details

Each environment consisted of a circular track with six laps, each lap containing five cues (S, A, B, C, E) in an order that permuted A, B, C across laps. Sensory input was one-hot vector: reward at positions A, C; no reward at B; start at S and end at E. The agent performed only one action “run”, which advanced the position deterministically. Starting position was randomly selected among the six lap-start states.

Training ran for 100,000 gradient updates, each processing a segment of 10 time-steps (truncated BPTT length) with learning rate=1e−4 using the Adam optimizer. L1 Regularization factors for the structural variable *g* and the conjunctive variable *p* were 0.01 and 0.02 in the regularized condition, respectively, and zero for both in the no-regularization condition. Environments were sampled continuously; each environment was traversed for a duration of approximately 120–330 steps. Memory matrices were reset at the start of each environment, and the binary reward mapping was fixed across environments. Evaluation was performed on 20 held-out environments with the same track structure.

#### Performance evaluation

Prediction accuracy at time step t was defined as the proportion of time steps in which the model’s predicted sensory symbol argmax 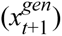 matched the true sensory input *x_t_*_+1_ . Accuracy was computed exclusively over time steps corresponding to previously visited nodes. To characterize the learned representations, the conjunctive variable *p_t_* and structure variable *g_t_* was analyzed analogously to the RNN hidden states.

Our rationale to use *g_t_* as an approximation of hippocampal activity is as follows: The hippocampus combines structural and sensory information, both of which were included in the calculation of *g_t_* using the inference model in TEM. In contrast, 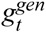 was calculated without access to sensory inputs. As a result, a few rewarded and non-rewarded conditions were merged in 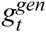 (Supplementary Fig. 21F-G). It should be noted that tasks in the original TEM paper are deterministic, whereas our task contains stochastic cue-position binding at rank 1 and rank 2. Therefore, 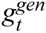 alone can not reliably represent task states in our task. Sensory cues in our task not only serve to perform cue-position binding, as proposed in the original paper, but also play an indispensable role in determining positions. In a stochastic environment, path integration needs continuous calibration from sensory cues.

As *g_t_*, *p* also integrates structural and sensory information. Theoretically, RSM of *p* across different conditions would be similar as *g_t_*. However, because of the clipping, *p* saturated too quickly and could not discriminate different states (Supplementary Fig. 21H). Removing clipping deteriorated the model’s stability during training, potentially from gradient vanishing/exploding. Therefore, we used *g_t_*, but not *p,* as our approximation of hippocampal representations.

## AI Usage Statement

Generative AI was used solely to improve the language and readability of the manuscript. It was not used to generate scientific content, analyses, interpretations, or conclusions.

