## Supplementary Figures for "Abstract representation in the hippocampus predicts spontaneously adopted structure-based behavioral strategy"

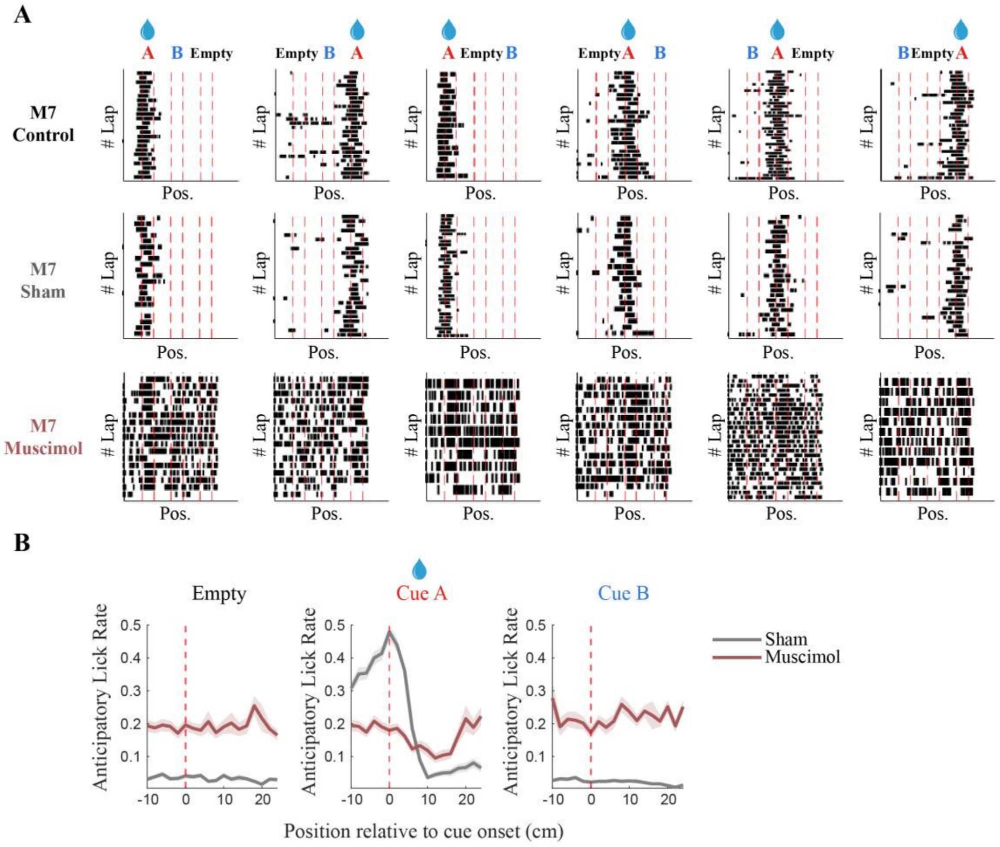


**Supplementary Figure 1. Behavioral effects of bilateral hippocampal silencing. (A)** Lick raster plot for a representative mouse under control, sham and muscimol conditions. Red vertical line indicates potential slots of cues. Black ticks denote licks. **(B)** Anticipatory lick rate (total licks excluding consumption-related licks) at three different slots. Consumption-related licks were defined as licks occurring after reward delivery until the licking rate fell below the threshold twice as large as the baseline level (baseline calculated from -4 to -3 s relative to reward). N=5 mice.

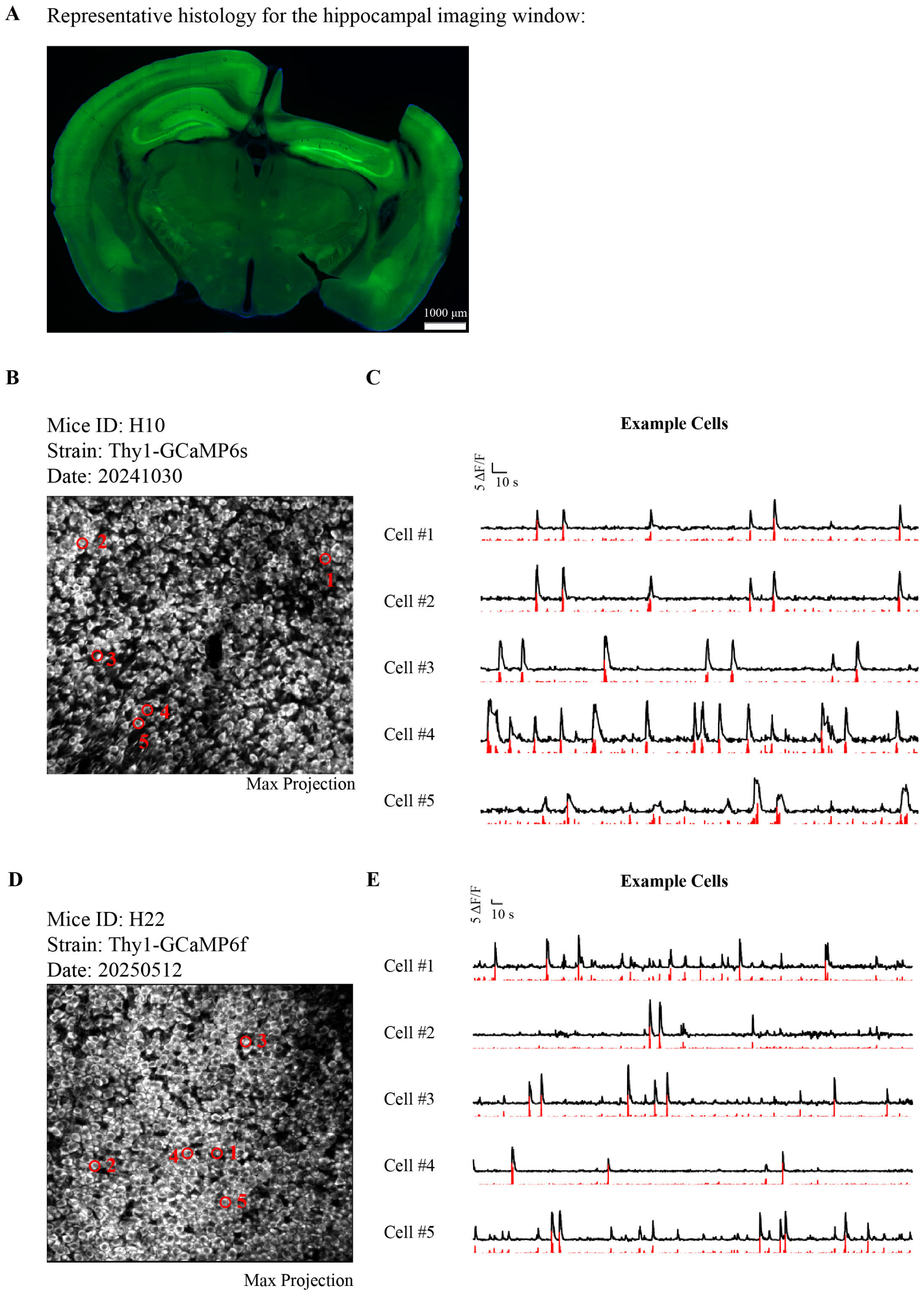


**Supplementary Figure 2. Two-photon calcium imaging in the dorsal CA1. (A)** Representative coronal section verifying the imaging location in the dorsal CA1 region of the hippocampus in a Thy1-GCaMP6f mouse. The black cavity indicates the implantation site of the titanium cannula. **(B)** Example two-photon imaging field from a Thy1-GCaMP6s mouse. Red circles indicate neurons corresponding to the example activity traces shown in panel C. **(C)** Example activity traces from the neurons indicated in panel B. Black traces represent ΔF/F signals, and red traces represent putative ‘spiking’ activity from deconvolution. **(D)** Example two-photon imaging field from a Thy1-GCaMP6f mouse. Red circles indicate neurons corresponding to the example activity traces shown in panel E. **(E)** Example activity traces from the neurons indicated in panel D. Black traces represent ΔF/F signals, and red traces represent deconvolved spike activity.

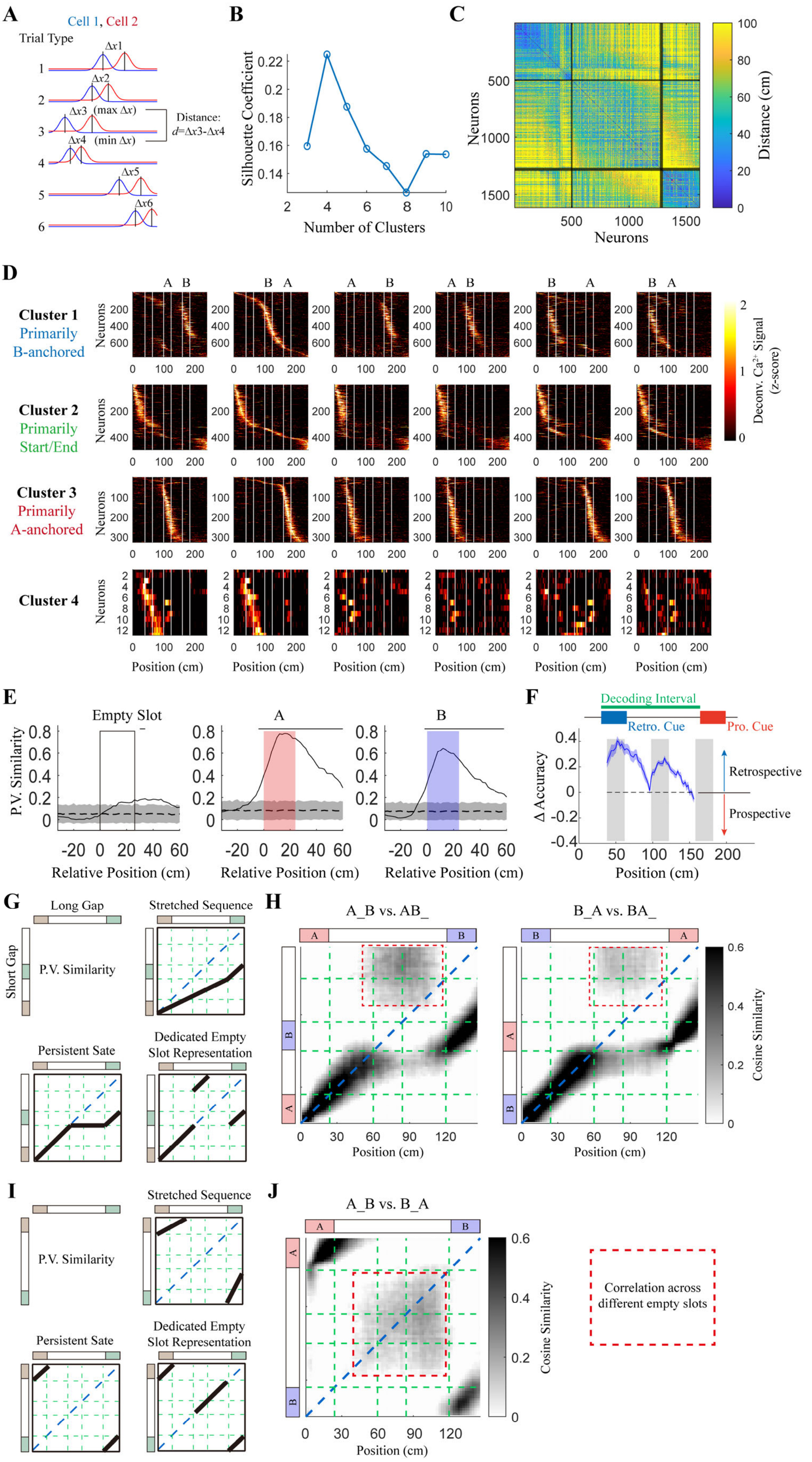


**Supplementary Figure 3. Hippocampal cell ensembles exhibited sequential representations of cued and empty slots. (A)** Definition of functional distance between pairs of neurons based on tuning curves across different trial types. **(B)** Silhouette coefficients obtained from hierarchical clustering across different numbers of clusters. **(C)** Hierarchical clustering of neurons based on functional distance into four clusters. Neurons from all mice were pooled. **(D)** Heat-maps of z-scored activity for the four neuronal clusters identified in panel (C). Neurons were sorted according to response peak location in the Empty–B–A trial type. **(E)** Population vector similarity within the spatial interval spanning 30 cm before to 60 cm after onset of cue A, cue B, and empty slots. Because each cue type appeared once in each trial type, six conditions were included. Pairwise cosine similarities were calculated and averaged across conditions. Dashed curves and shaded regions represent mean ± s.d. of shuffled controls generated by random permutation of neuron identities. Horizontal bars indicate intervals in which real and shuffled data differed significantly ($\alpha = 0.05$; Wilcoxon signed-rank test with Holm–Bonferroni correction). **(F)** Decoding of the retrospective and prospective cues at each position bin based on population neural activity. Linear support vector machine (SVM) was used for decoding. Difference in decoding accuracies (retrospective-prospective) was plotted (curve and shade represent mean and s.e.m., respectively; n = 12 mice). Only the first and second slots were included in the analysis since the third slot did not have the 'prospective' cue. The decoding power for the retrospective cue were higher than the prospective cue throughout the track, with the difference minimized before the onset of the prospective cue, where mice were already able to see it. **(G)** Theoretical predictions of population vector similarity between conditions with short and long gaps separating cue A and cue B under different hypotheses. **(H)** Population vector similarity between conditions with short and long gaps separating cue A and cue B calculated from experimental data. **(I)** Theoretical predictions of population vector similarity between long-gap conditions with different configurations of cue A and cue B under different hypotheses. **(J)** Population vector similarity between long-gap conditions with different configurations of cue A and cue B calculated from experimental data.

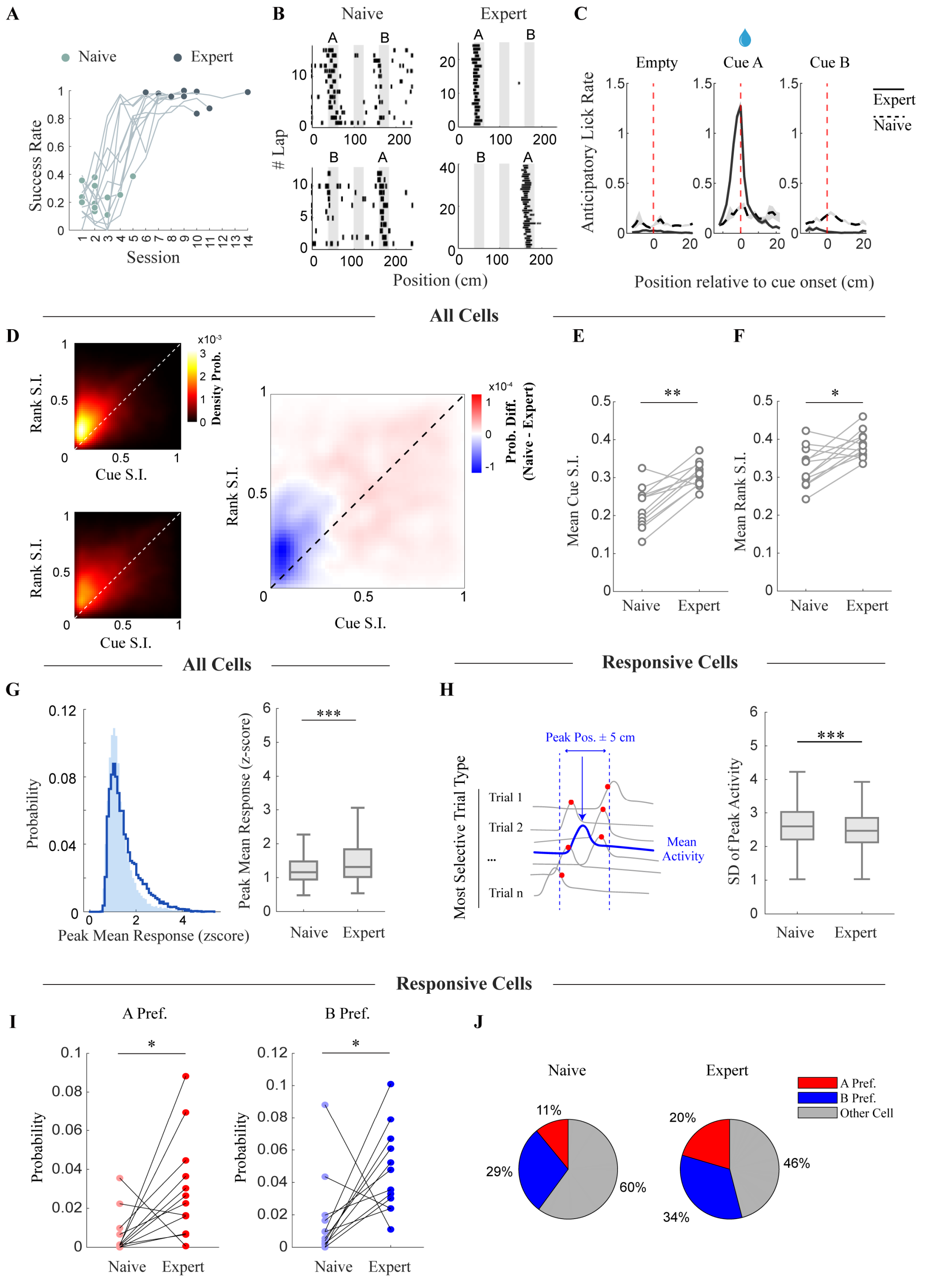


**Supplementary Figure 4. Enhanced feature selectivity in the hippocampus during learning in the A/B/Empty task. (A)** Success rate across all sessions in the A/B/Empty task. N =12 mice. **(B)** Lick raster map of an example mouse from naive to expert in trial types A-Empty-B and B-Empty-A. Black ticks indicate lickings; gray squares denote possible cue locations. **(C)** Anticipatory lick rate (total licks excluding consumption-related licks) at three different slots. Consumption licks were defined as licks occurring after reward delivery until the licking rate fell below the threshold twice as large as the baseline level (baseline calculated from -4 to -3 s relative to reward). Curves and shaded regions represent mean $\pm$ s.e.m., respectively. N=12 mice. **(D)** 2-D density map of cue selectivity indices and rank selectivity indices in naive (upper left) and expert mice (lower left).

n = 13680 cells for naive and n = 14380 cells for expert from 12 mice. The right panel shows the difference in probability density across selectivity index space (expert minus naive). Naive and expert sessions were shown in panel A. **(E)** Mean cue selectivity indices of naive and expert mice. All cells were included. $P = 0.0015$; two-sided Wilcoxon signed-rank test; n = 12 mice. **(F)** Mean rank selectivity indices of naive and expert mice. All cells were included. $P = 0.0122$; two-sided Wilcoxon signed-rank test; n = 12 mice. **(G)** Distribution of peak responses across all neurons (left). Light shaded histograms indicate naive mice, and dark solid lines indicate expert mice. Quantification of the distributions shown in the left panel (right). $P = 6.601 \times 10^{-69}$; two-sided Wilcoxon rank-sum test. **(H)** Schematic illustrating the calculation of trial-to-trial response variability (left). For each neuron, the trial type with the highest peak response was identified as its preferred trial type. Response curves were then averaged across all trials of the preferred trial type (blue line). The peak position of this mean curve was identified, and a ±5 cm window centered on the peak was used to extract the peak activity from each individual trial. Standard deviation was calculated from these trial-by-trial peak activities. To minimize the influence of inactive neurons, the analysis was restricted to highly responsive cells. Distribution of response variability calculated using the method shown in the left panel (right). $P = 5.430 \times 10^{-8}$; two-sided Wilcoxon rank-sum test; n = 898 neurons for naive mice and n = 1726 neurons for expert mice from 12 mice. **(I)** Quantification of cue A-preferring (left) and cue B-preferring (right) cells in naive and expert mice. Cue preferring cells were defined as neurons with a maximum median response > 2 and a selectivity index for the preferred cue > 0.5. $P = 0.0269$ for A-preferring cells; $P = 0.0425$ for B-preferring cells; two-sided Wilcoxon signed-rank test. Neurons included in the analysis were the same as those in panel H. **(J)** Proportions of A-preferring, B-preferring, and other cells in naive and expert mice. Neurons included in the analysis were the same as those in panel H.

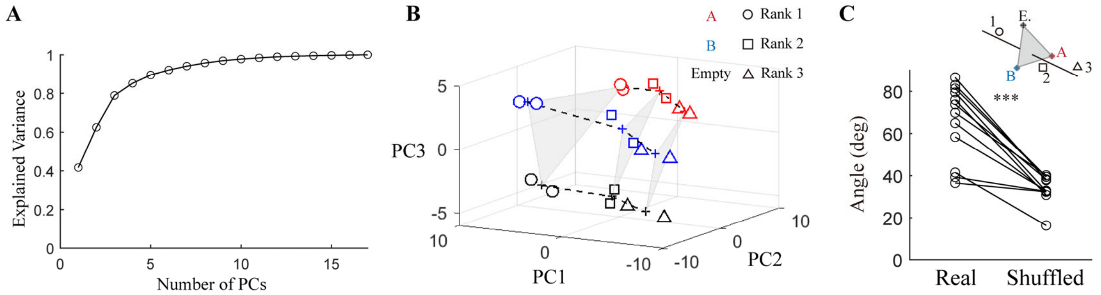


**Supplementary Figure 5. Principal component analysis (PCA) of hippocampal representations in the A/B/Empty task. (A)** Explained variance across different numbers of principal components (PCs). **(B)** Population vectors projected in the 3-dimensional PC space. Red, blue, and black denote cue A, cue B, and the empty slot, respectively. Circles, squares, and triangles denote rank 1, rank 2, and rank 3, respectively. Crosses denoted centers of mass of the two data points with the same cue/rank combination. **(C)** Angle between the cue-coding plane and rank-coding axis, calculated in the 3-dimensional PC space. $P = 0.0004883$, real versus shuffled cell identity; two-sided Wilcoxon signed-rank test; $n = 12$ mice.

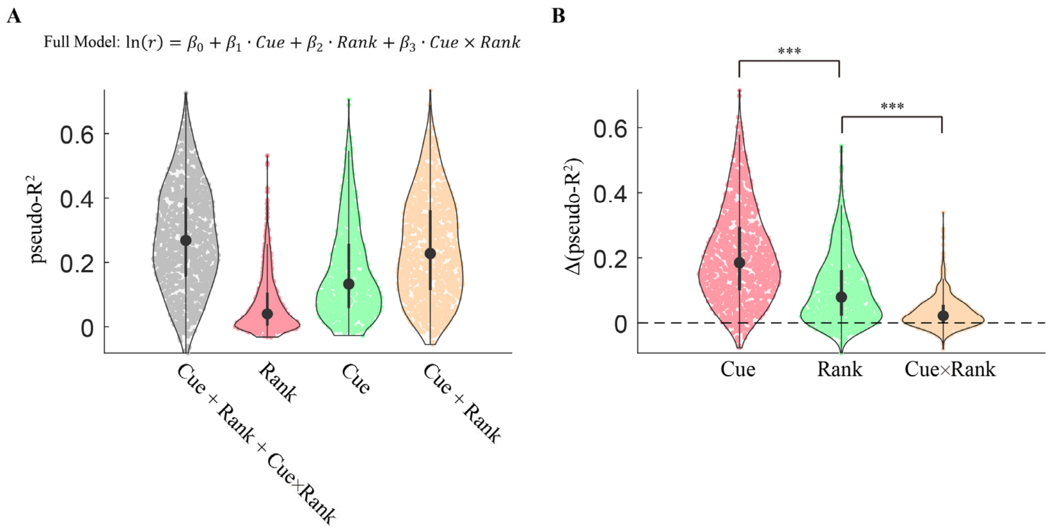


**Supplementary Figure 6. Trial-by-trial activity variance of single hippocampal neurons explained by various task-relevant variables using generalized linear models (GLM). (A)** pseudo-$R^2$ in the full model (gray), and models with rank (pink), cue (green), and the linear summation of rank and cue (yellow) as predictors, respectively. Neurons were pooled from all mice. Only neurons with responses peaks within the three potentially cued slots were included (neurons preferentially activated at trial start and trial end were excluded). **(B)** Relative contributions of cue, rank, and their nonlinear interaction, quantified with Δ(pseudo-$R^2$). Δ(pseudo-$R^2$)(cue) is defined as the subtraction of pseudo-$R^2$(rank-only) from pseudo-$R^2$(full model). Δ(pseudo-$R^2$)(rank) is defined as the subtraction of pseudo-$R^2$(cue-only) from pseudo-$R^2$(full model). Δ(pseudo-$R^2$)(cue×rank) is defined as the subtraction of pseudo-$R^2$(cue+rank) from pseudo-$R^2$(full model). $P < 1 \times 10^{-300}$, cue vs. rank, and rank vs. cue+rank, two-sided Wilcoxon signed-rank test, n=1117 neurons.

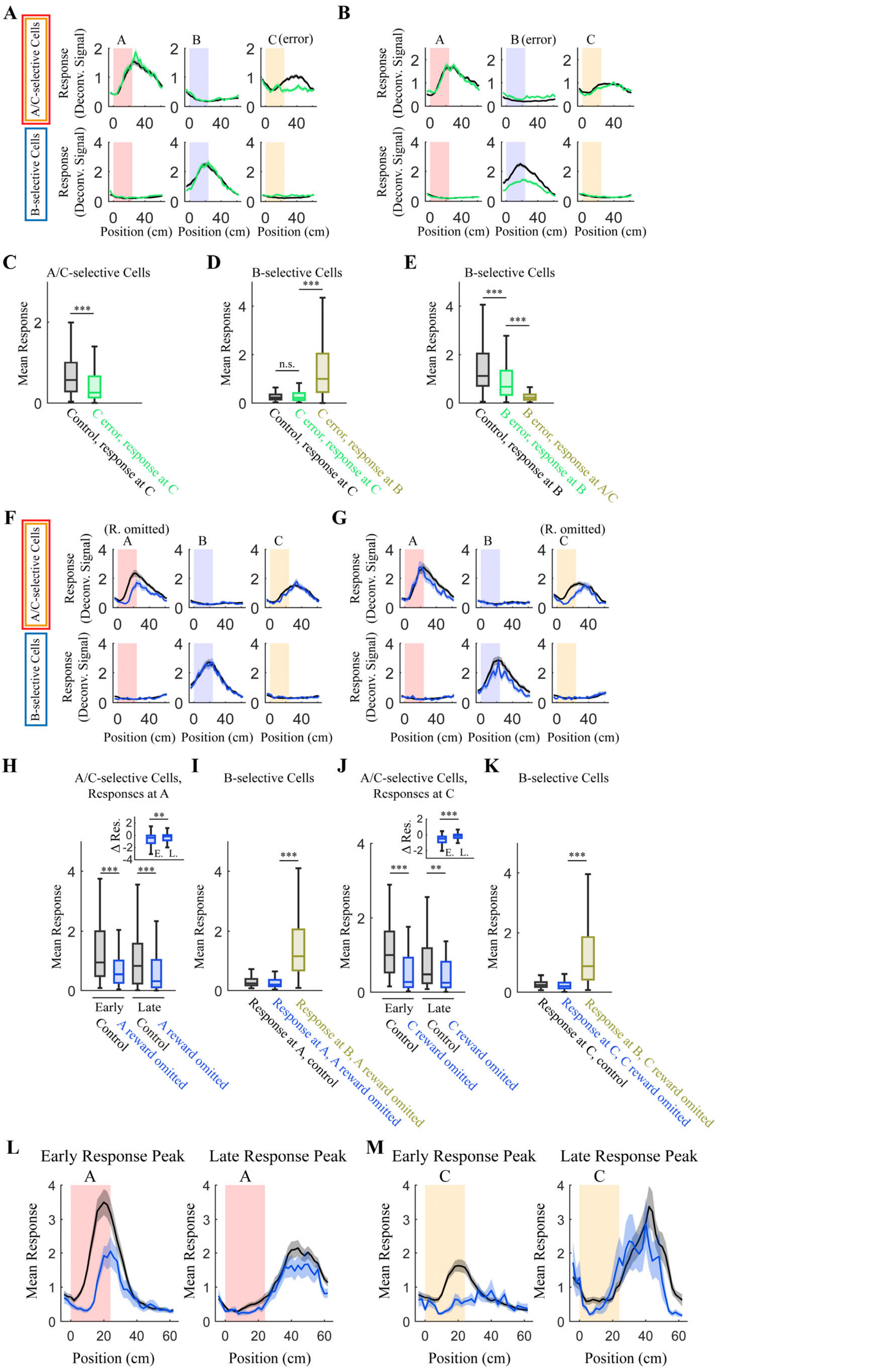


**Supplementary Figure 7**. **Analysis of hippocampal representations on day 1 in error and catch trials. (A)** Responses of A/C-selective cells (upper row) and B-selective cells (lower row) to different cues in control (black) and C-error (green) trials. Curves and shaded areas represent mean ± s.e.m., respectively. **(B)** The same as in (A), but in control (black) and B-error trials (green). **(C)** Quantification of A/C-selective cells in (A). Peak activities in each cell were used for statistical comparison. $P = 5.635 \times 10^{-19}$, response at C in control trials vs. response at C in C-error trials; two-sided Wilcoxon signed-rank test, n=561 cells. **(D)** Quantification of B-selective cells in (A). Peak activities in each cell were used for statistical comparison. $P = 0.0732$, response at C in control trials vs. response at C in C-error trials; $P < 1 \times 10^{-300}$, response at C in C-error trials vs. response at B in C-error trials; two-sided Wilcoxon signed-rank test, n=484 cells. **(E)** Quantification of B-selective cells in (B). Peak activities in each cell were used for statistical comparison. $P < 1 \times 10^{-300}$, response at B in control trials vs. response at B in B-error trials; $P < 1 \times 10^{-300}$, response at B in B-error trials vs. response at A/C in B-error trials; two-sided Wilcoxon signed-rank test, n=724 cells. **(F)** Responses of A/C-selective cells (upper row) and B-selective cells (lower row) to different cues in control (black) and A-catch (blue) trials. Curves and shaded areas represent mean ± s.e.m., respectively. Rewards were omitted at cue A in A-catch trials. **(G)** The same as in (F), but in control (black) and C-catch trials (blue). Rewards were omitted at cue C in C-catch trials. **(H)** Quantification of A/C-selective cells in (F). Peak activities in each cell were used for statistical comparison. $P < 1 \times 10^{-300}$, early phase (0-30cm after the cue onset) response at A in control trials vs. A-catch trials; $P = 0.0001796$, late phase (30-60cm after the cue onset) response at A in control trials vs. A-catch trials; Inset: difference of responses between control and catch trials for early (E.) and late (L.) phases, $P = 0.0044$, early vs. late phases; two-sided Wilcoxon signed-rank test, n=174 cells. **(I)** Quantification of B-selective cells in (F). Peak activities in each cell were used for statistical comparison. $P < 1 \times 10^{-300}$, response at A vs. B in A-catch trials; two-sided Wilcoxon signed-rank test, n=176 cells. **(J)** Quantification of A/C-selective cells in (G). Peak activities in each cell were used for statistical comparison. $P < 1 \times 10^{-300}$, early phase (0-30cm after the cue onset) response at C in control trials vs. C-catch trials; $P = 0.009273$, late phase (30-60cm after the cue onset) response at C in control trials vs. C-catch trials; Inset: difference of responses between control and catch trials for early (E.) and late (L.) phases, $P = 4.778 \times 10^{-8}$, early vs. late phases; two-sided Wilcoxon signed-rank test, n=132 cells. **(K)** Quantification of B-selective cells in (G). Peak activities in each cell were used for statistical comparison. $P < 1 \times 10^{-300}$, response at C vs. B in C-catch trials; two-sided Wilcoxon signed-rank test, n=134 cells. **(L)** Averaged responses to cue A of cells with response peak within 24cm, or beyond 40cm, relative to the cue onset in control (black) and A-catch (blue) trials. Curves and shaded areas represent mean ± s.e.m., respectively. **(M)** The same as in (L), in C-catch trials.

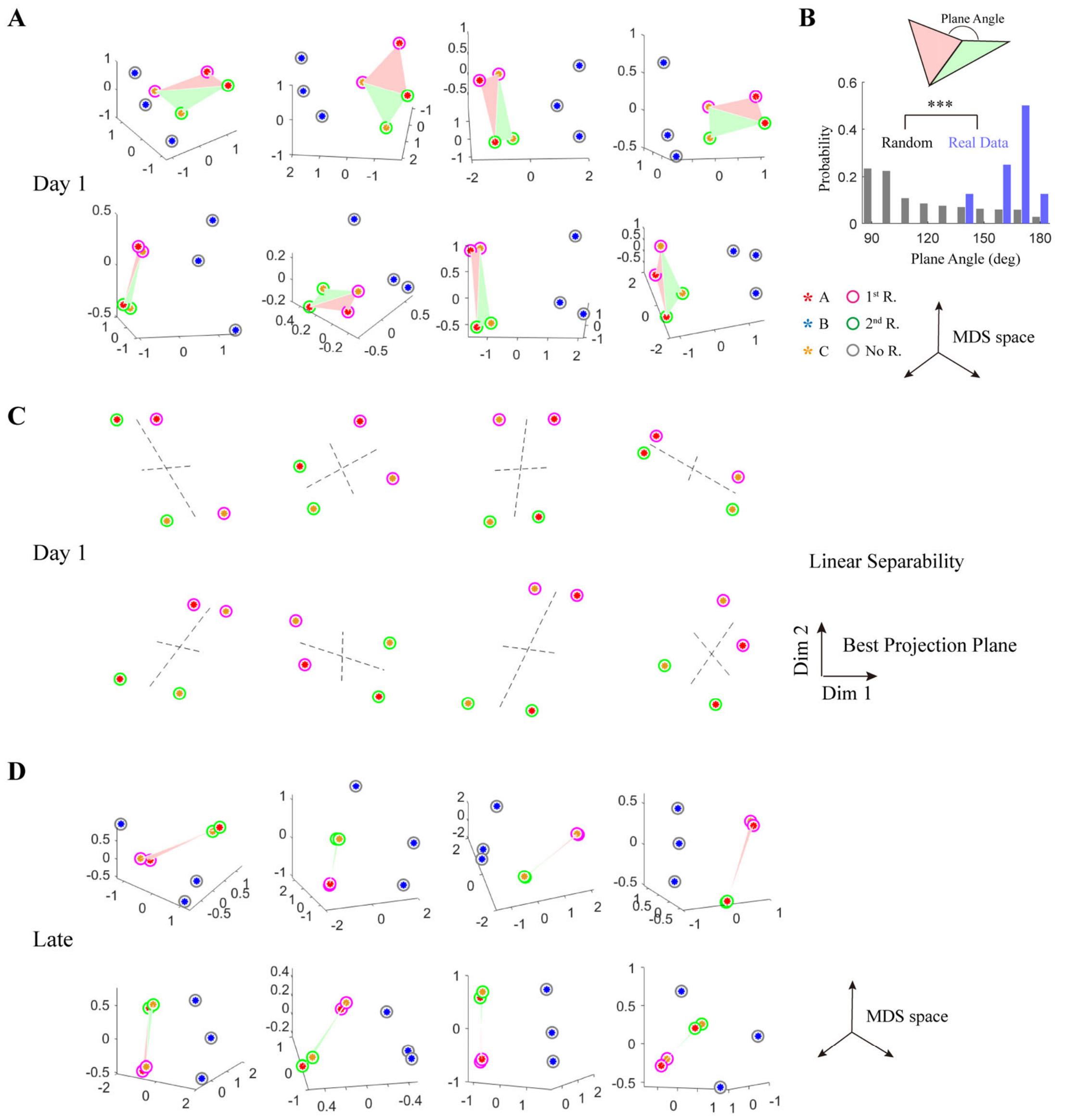


**Supplementary Figure 8. Map transformation in all eight mice in the reward order-encoding group. (A)** Three-dimensional projections of population vectors in the MDS space in each mouse. Red, blue, and yellow stars denote cue A, B, and C, respectively. Magenta and green circles denote the first and the second reward for cue A/C, respectively, whereas gray circles indicate no-reward (cue B). Pink and green triangles denote the two planes used to quantify whether the four points are located within the same plane. Neural activities imaged on day 1 were analyzed. **(B)** Distribution of angles between the pink and green triangles, as shown in (A), in real data (blue) and simulated four random points (gray). $P = 5.645 \times 10^{-5}$, Kolmogorov-Smirnov test, n=8 and 1000 for real and simulated data, respectively. **(C)** A plane was fitted using points from the four rewarded conditions in each mouse. The four points were then projected to this plane to visualize their geometric organization. Neural activities imaged on day 1 were analyzed. **(D)** Same as in (A), with neural activities imaged in late sessions.

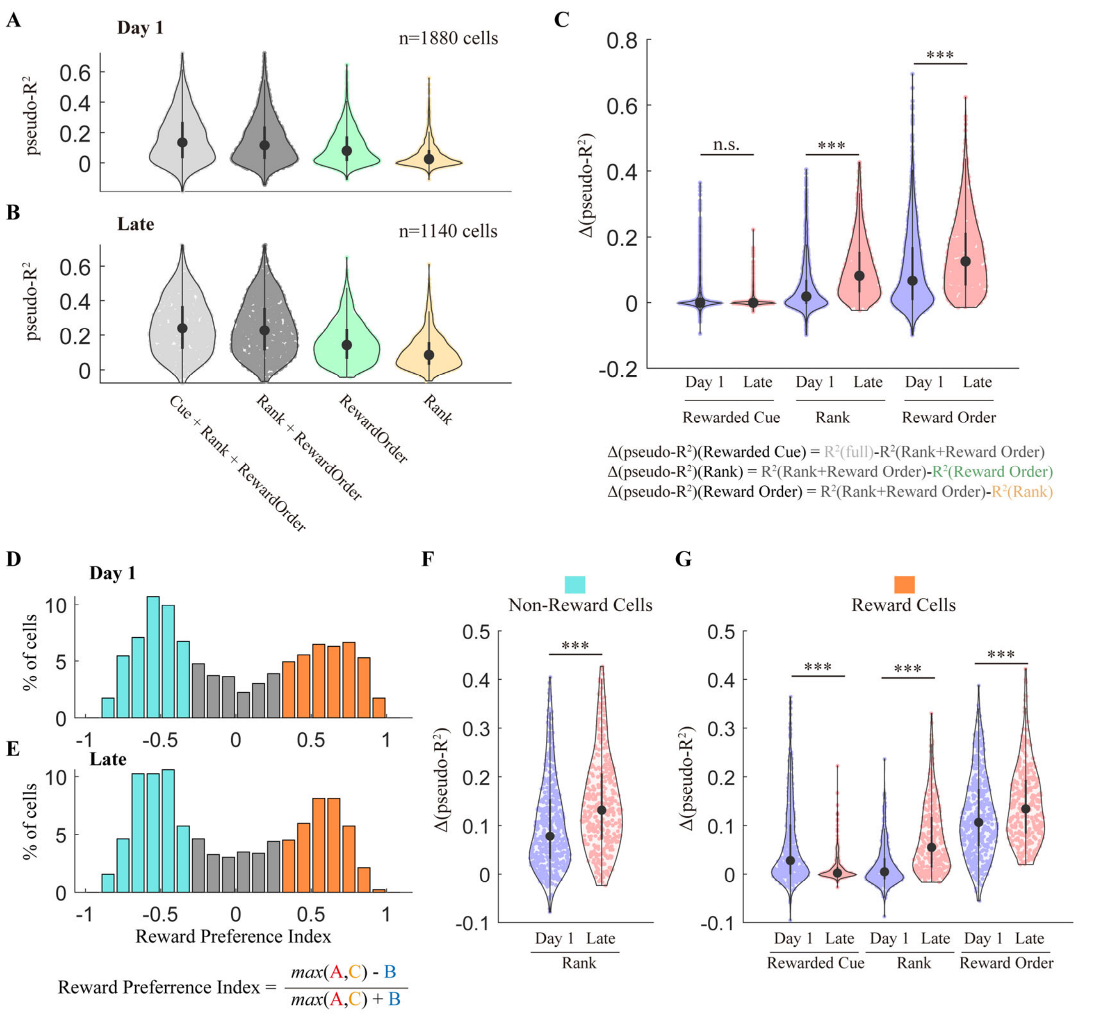


**Supplementary Figure 9. Changes of relative contributions of various task-relevant variables to hippocampal neural activities. (A)-(B)** pseudo-$R^2$ in the full model (light gray), and models with rank+reward-order (dark gray), reward-order (green), and rank (yellow) as predictors, respectively. Neurons were pooled from all mice in the reward order-encoding group. Only neurons with responses peaks within the three potentially cued slots were included (neurons preferentially activated at trial start and trial end were excluded). **(C)** $\Delta$(pseudo-$R^2$) for different variables on day 1 (purple) and late sessions (pink). How each $\Delta$(pseudo-$R^2$) was calculated was described in the figure. $P = 5.645 \times 10^{-5}$ for cue, $< 1 \times 10^{-300}$ for rank, $P$=$3.307 \times 10^{-36}$ for reward-order, day 1 vs. late, two-sided Wilcoxon rank sum test, n=1880 and 1140 cells for day 1 and late, respectively. **(D)-(E)** Distributions of reward preference index. Reward (orange) and non-reward (blue) preferring cells were defined as the index >0.33 or <-0.33, respectively. **(F)** $\Delta$(pseudo-$R^2$) for rank in non-reward preferring cells. $P = 2.716 \times 10^{-12}$, day 1 vs. late, two-sided Wilcoxon rank sum test, n=482 and 382 cells for day 1 and late, respectively. **(G)** $\Delta$(pseudo-$R^2$) for various variables in reward preferring cells. $P = 6.181 \times 10^{-12}$ for cue, $3.550 \times 10^{-38}$ for rank, $2.210 \times 10^{-5}$ for reward-order, day 1 vs. late, two-sided Wilcoxon rank sum test, n=427 and 309 cells for day 1 and late, respectively.

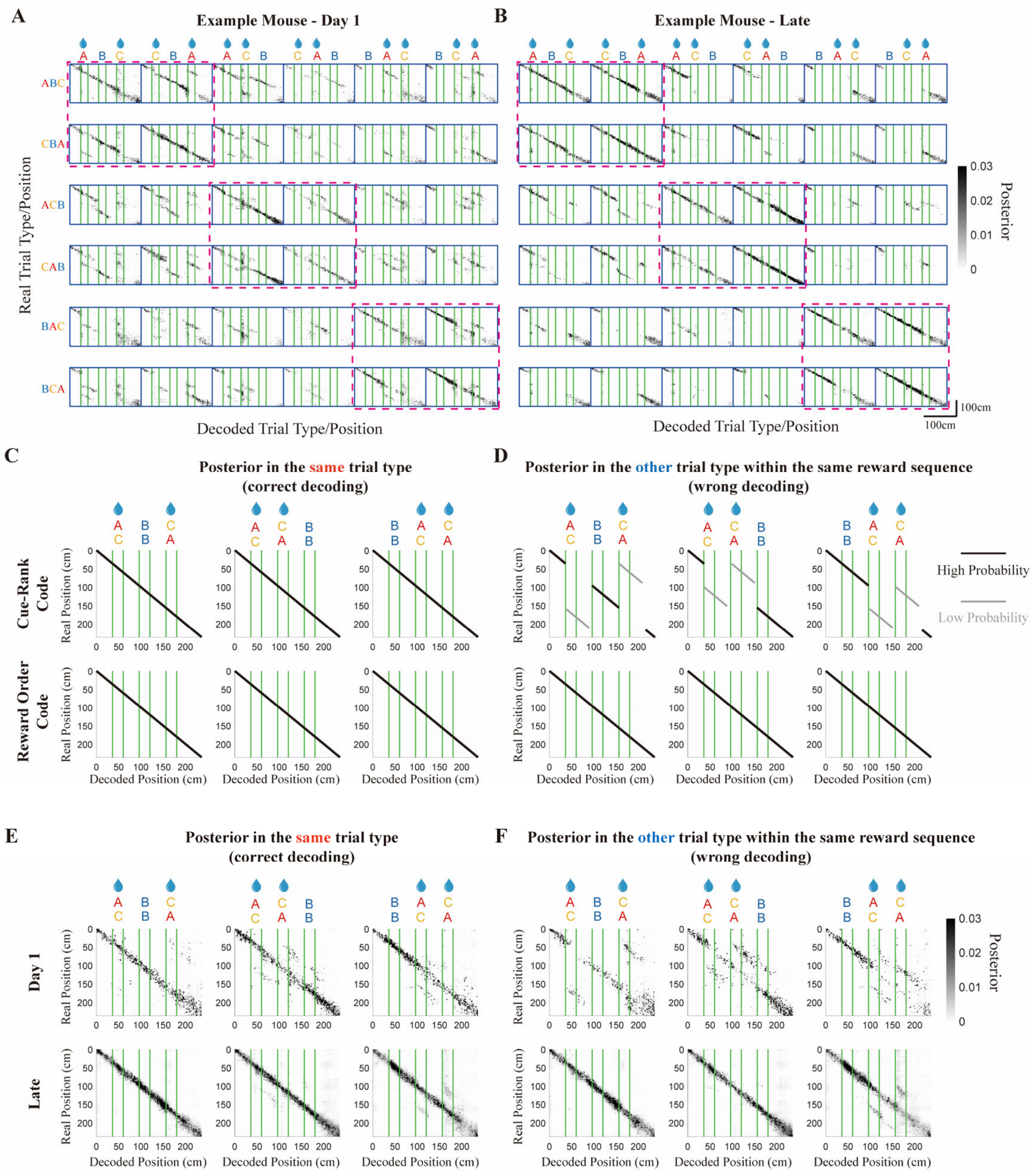


**Supplementary Figure 10. Bayesian decoding of trial-type × position. (A)-(B)** Posterior probability distributions in day 1 (A) and late (B) sessions in one example mouse. **(C)-(D)** Theoretical distributions of posterior probability in the same trial type (C) and the different trial type with the same reward sequence (D), under difference hypotheses about the representation patterns. If the hippocampal representation is a linear combination of cue and rank, partial correlations between A and C are expected. **(E)-(F)** Posterior probability distributions in the same trial type (E) and the different trial type with the same reward sequence (F), calculated from experimental data in day 1 (upper row) and late sessions (lower row). Posterior probability was first calculated in each individual mouse and then averaged across mice (n=8).

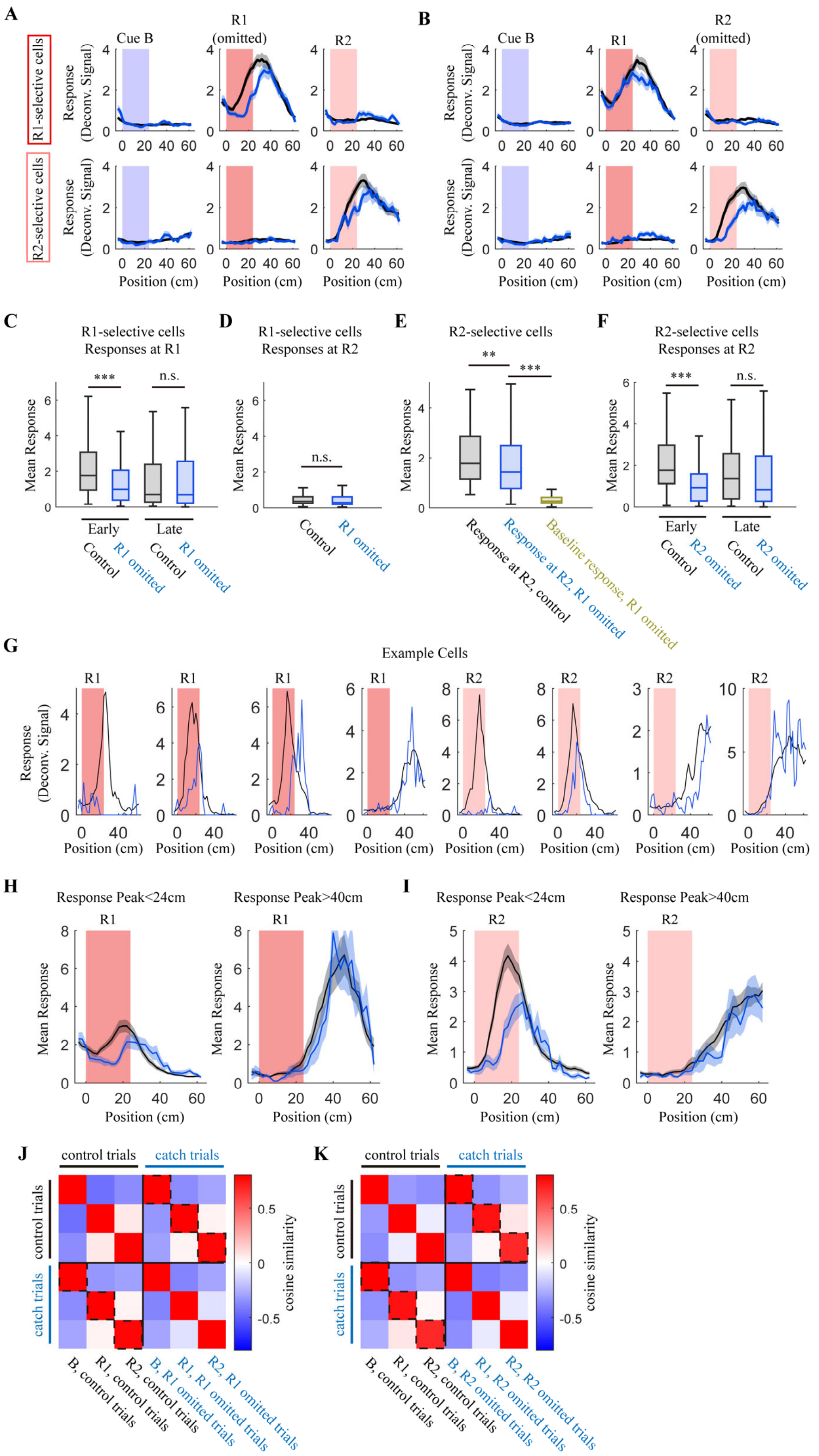


**Supplementary Figure 11**. **Analysis of hippocampal representations in late session catch trials.** **(A)** Responses of R1-selective cells (upper row) and R2-selective cells (lower row) to different slots in control (black) and R1-catch (blue) trials. R1(dark pink) and R2 (light pink) denote the first and second rewards, respectively. Curves and shaded areas represent mean ± s.e.m., respectively. **(B)** The same as in (A) in R2-catch trials. **(C)** Quantification of R1-selective cells in (A). Peak activities in each cell were used for statistical comparison. $P < 1 \times 10^{-300}$, early phase (0-30cm after the cue onset) response at R1 in control trials vs. R1-catch trials; $P = 0.7689$, late phase (30-60cm after the cue onset) response at R1 in control trials vs. R1-catch trials; two-sided Wilcoxon signed-rank test, n=228 cells. **(D)** Quantification of R1-selective cells in (A). Peak activities in each cell were used for statistical comparison. $P = 0.4149$, response at R2 in control trials vs. R1-catch trials; two-sided Wilcoxon signed-rank test, n=228 cells. **(E)** Quantification of R2-selective cells in (A). Peak activities in each cell were used for statistical comparison. $P = 0.0260$, response at R2 in control trials vs. R1-catch trials; $P < 1 \times 10^{-300}$, response at R2 vs. R1 in R1-catch trials; two-sided Wilcoxon signed-rank test, n=127 cells. **(F)** Quantification of R2-selective cells in (B). Peak activities in each cell were used for statistical comparison. $P < 1 \times 10^{-300}$, early phase (0-30cm after the cue onset) response at R2 in control trials vs. R2-catch trials; $P = 0.0558$, late phase (30-60cm after the cue onset) response at R2 in control trials vs. R2-catch trials; two-sided Wilcoxon signed-rank test, n=126 cells. **(G)** Responses of example cells in R1 and R2 selective trials. **(H)** Averaged responses to R1 of cells with response peak within 24cm, or beyond 40cm, relative to the cue onset in control (black) and R1-catch (blue) trials. Curves and shaded areas represent mean ± s.e.m., respectively. **(I)** Same as in (H), in R2-catch trials. **(J)-(K)** Representation similarity matrix across different conditions between control and R1 (J) or R2 (K) catch trials.

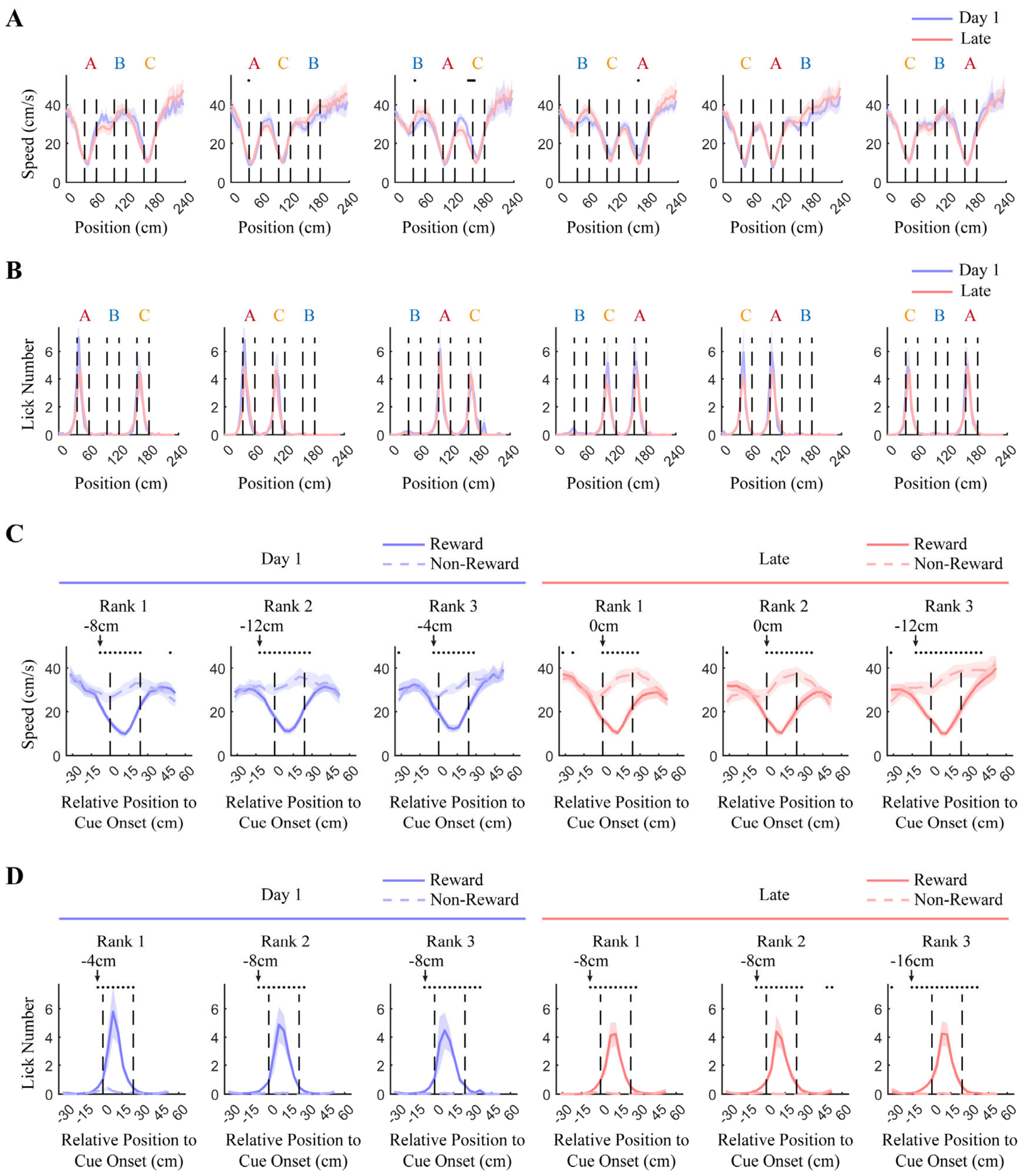


**Supplementary Figure 12. Running speed and lick rate on day 1 and late sessions. (A)** Positional profiles of running speed across different trial types on day 1 (purple) and late sessions (pink). Curves and shaded areas represent mean ± s.e.m., respectively, n=8 mice. Black dots above curves mark bins with significant differences (α = 0.05; day 1 vs. late; two-sided Wilcoxon signed-rank test with Holm–Bonferroni correction; n=8 mice). **(B)** Same as (A), with lick rate on day 1 (purple) and late (pink) sessions. No significant difference was detected across all trial-types/positions (α = 0.05; day 1 vs. late; two-sided Wilcoxon signed-rank test with Holm–Bonferroni correction; n=8 mice). **(C)** Divergence of running speed around reward (solid curves) and non-reward (dashed curves) conditions at different ranks. Curves and shaded areas represent mean ± s.e.m., respectively, n=8 mice. Black dots above curves mark bins with significant differences (α = 0.05; reward vs. non-reward; two-sided Wilcoxon signed-rank test with Holm–Bonferroni correction; n=8 mice). **(D)** Same as (C), with lick rates analyzed.

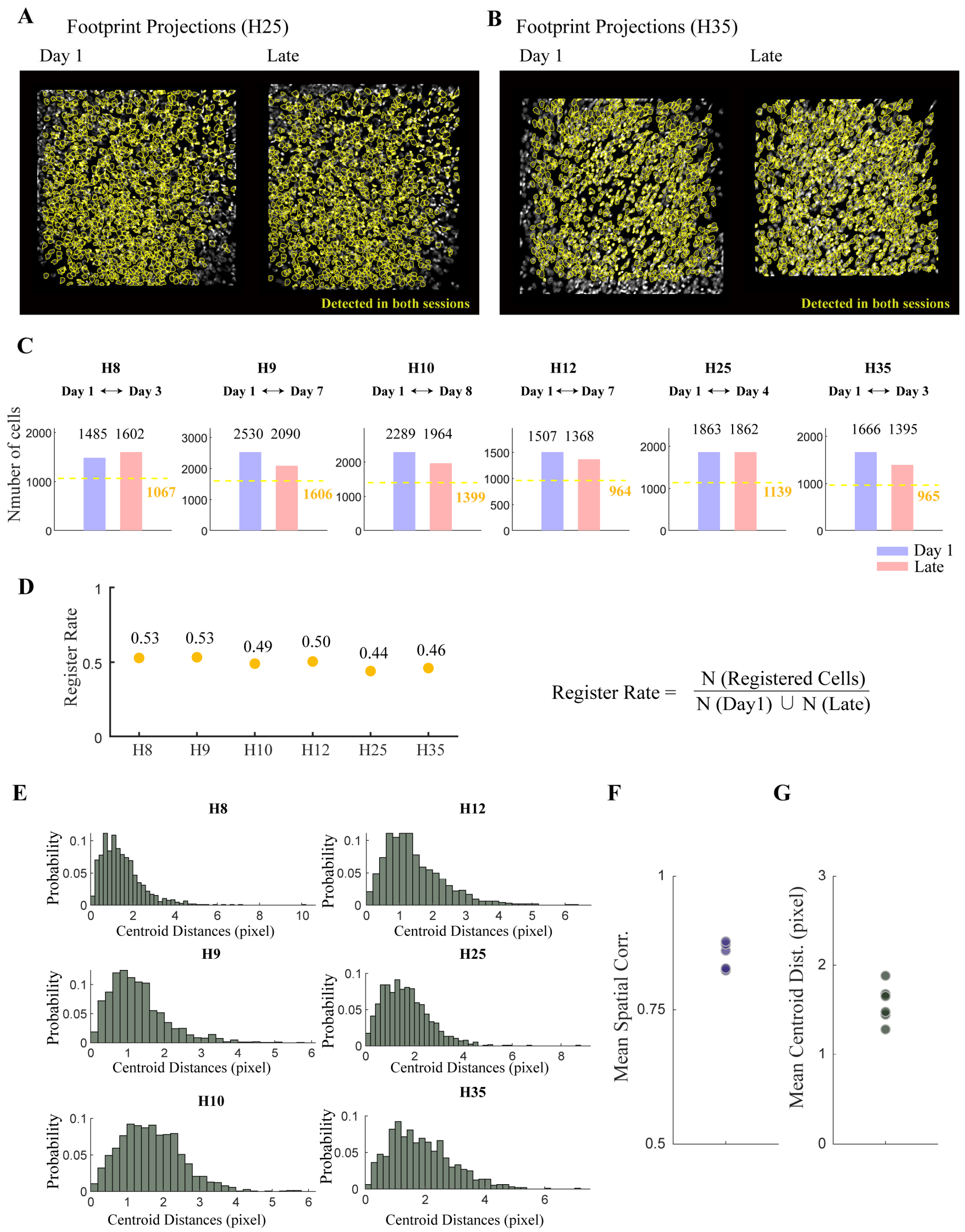


**Supplementary Figure 13. Cell registration across imaging sessions. (A-B)** Footprint projections from two example mice on day 1 and late sessions. Yellow outlines indicate registered neurons. **(C)** Numbers of neurons identified by Suite2p and neurons successfully registered across sessions. Yellow dashed lines indicate the number of registered neurons for each mouse. **(D)** Registration rate for each mouse. **(E)** Distribution of centroid distances for all neuron pairs identified as registered cells. Registered neurons were first matched based on high spatial correlation, and centroid distance was subsequently used as a post hoc validation measure. **(F)** Quantification of mean spatial correlation for registered neurons; n = 6 mice. **(G)** Quantification of mean centroid distance for registered neurons; n = 6 mice.

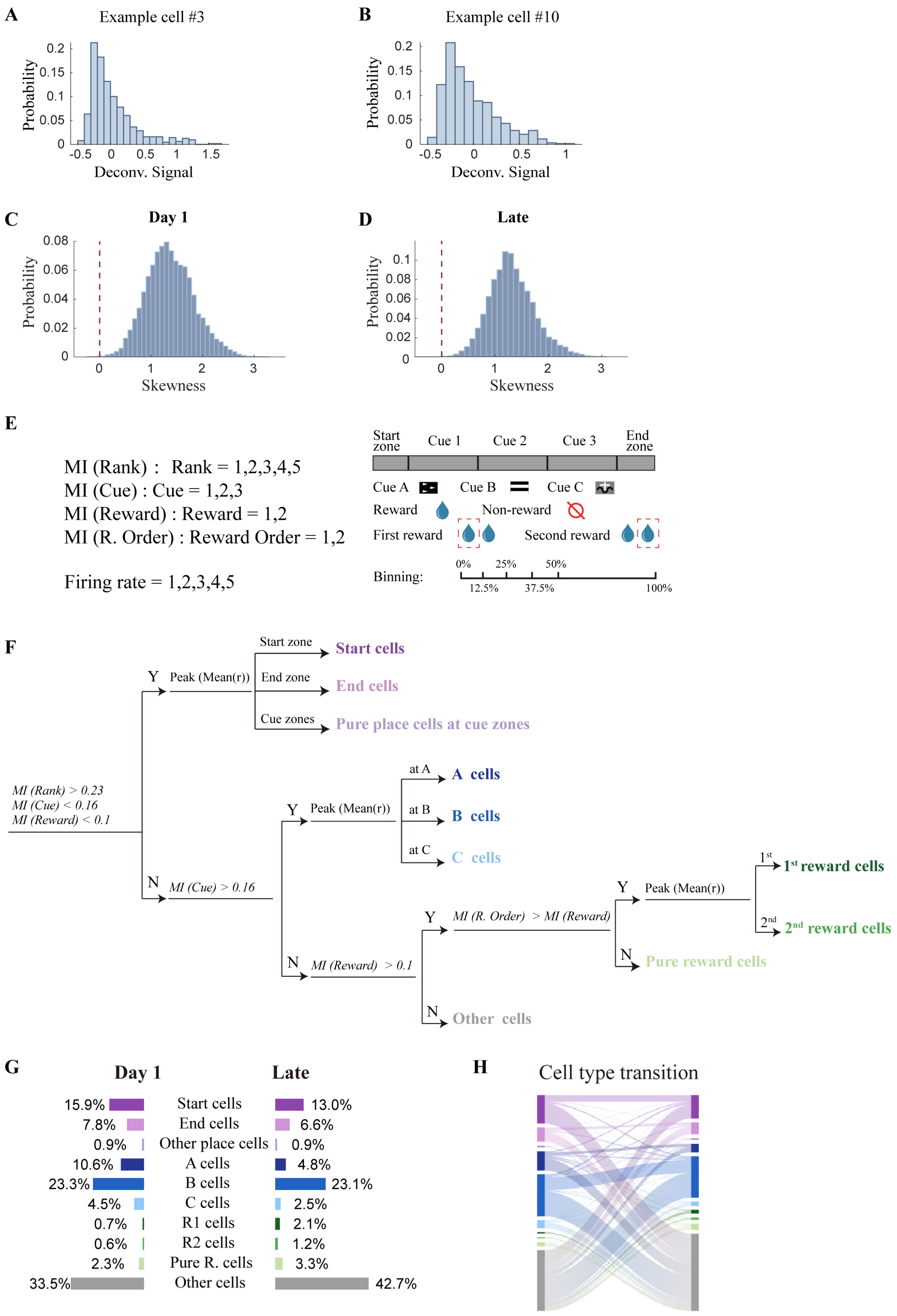


**Supplementary Figure 14. Mutual information analysis and neuronal classification. (A-B)** Distributions of deconvolved activity from two example neurons. **(C-D)** Distributions of response skewness in early and late sessions. Red dashed lines indicate zero skewness. Skewness was calculated from the distribution of deconvolved activity for each neuron. **(E)** Schematic illustration of the variables used for mutual information analysis. Because the response distributions of all neurons exhibited positive skewness, neural responses were discretized using unequally sized bins to achieve a more balanced sampling across response levels. **(F)** Decision tree illustrating the criteria used for neuronal classification. **(G)** Cell-type proportions on day 1 and late sessions. Cells included in the analysis comprised all registered neurons except those classified as 'other' in both day 1 and late sessions. **(H)** Sankey plot of cell-type transitions from day 1 to late sessions. Data and color definitions are the same as in panel G.

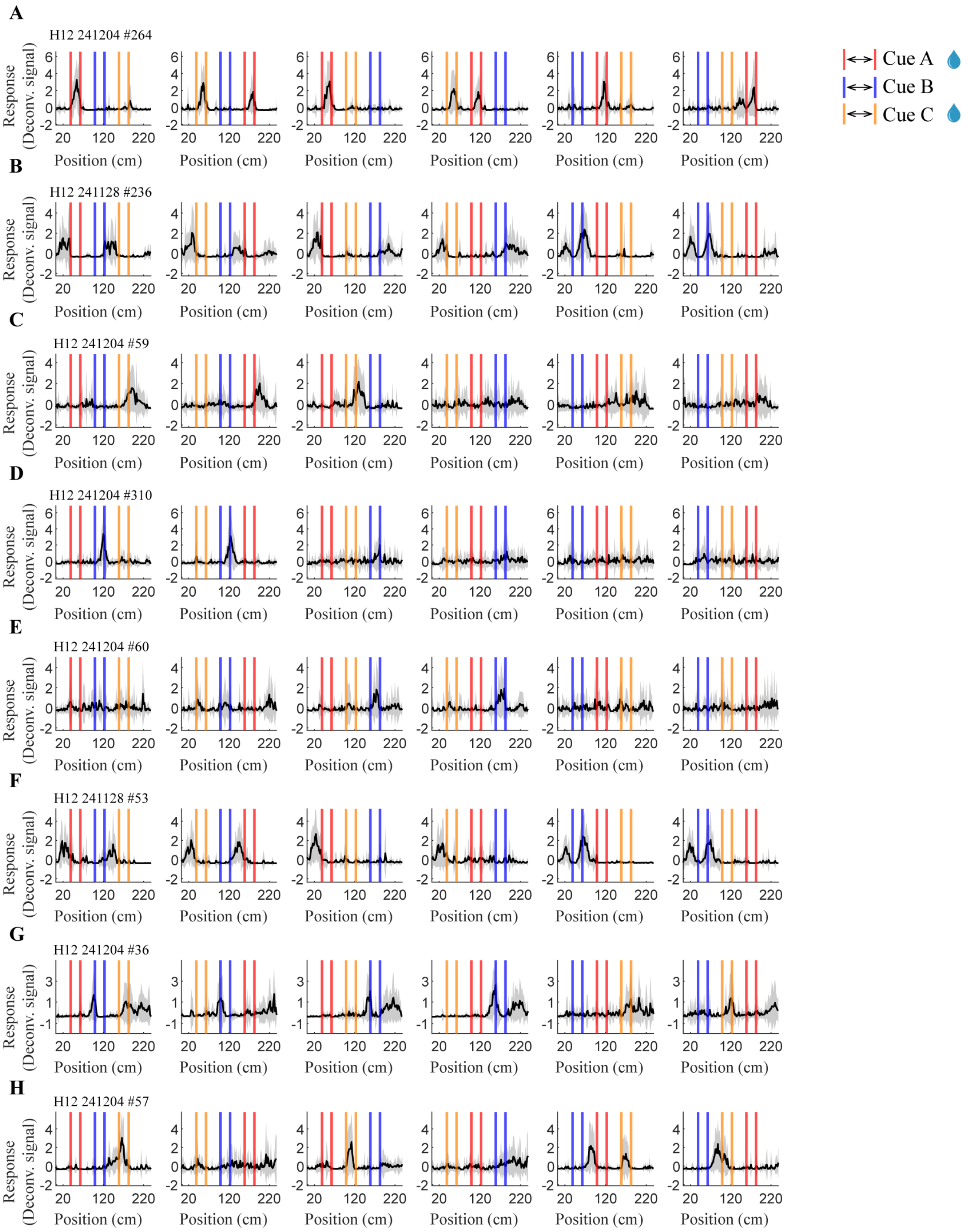


**Supplementary Figure 15. Cells with mixed selectivity in A/B/C task. (A-H)** Representative mixed-selectivity neurons. Red, blue, and yellow lines indicate the positions at which cues A, B, and C were presented, respectively. Black curves and shaded areas represent mean and s.e.m., respectively.

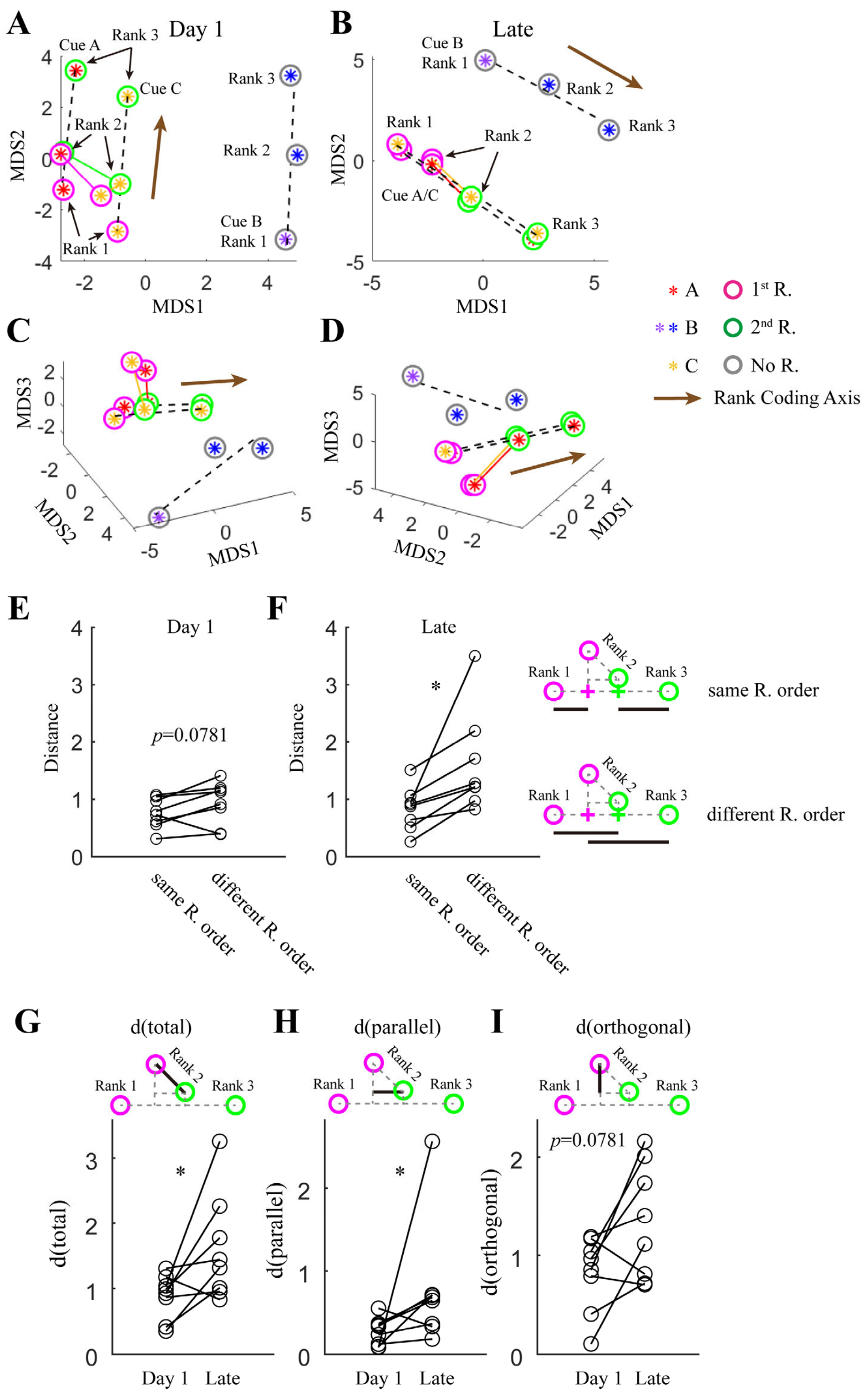


**Supplementary Figure 16. Detailed geometric analysis of neural representations at rank 2. (A)-(B)** The first two dimensions of 3-dimensional projections in the MDS space on day 1 (A) and late sessions (B). Colors of stars and circles represent different cues and reward orders, as in Fig. 2, except for that a special color (purple) was used to label cue B at rank 1. Brown axes represent the rank coding axis. **(C)-(D)** The same as in (A) and (B), showing the third dimension in the MDS space. **(E)-(F)** Separation of rank-2 reward representations based on reward order on day 1 (E) and late sessions (F). $P = 0.0781$ for day 1, 0.0078 for late, same vs. different reward orders; two-sided Wilcoxon signed-rank test; n=8 mice. **(G)** Overall distances of rank-2 reward representations in 3-dimensional MDS space. $P = 0.0391$, day1 vs. late; two-sided Wilcoxon signed-rank test; n=8 mice. **(H)** Distances of rank-2 reward representations in 3-dimensional MDS space parallel to the rank-coding axis. $P = 0.0391$, day1 vs. late, two-sided Wilcoxon signed-rank test; n=8 mice. **(I)** Distances of rank-2 reward representations in 3-dimensional MDS space orthogonal to the rank-coding space. $P = 0.0781$, day1 vs. late; two-sided Wilcoxon signed-rank test; n=8 mice.

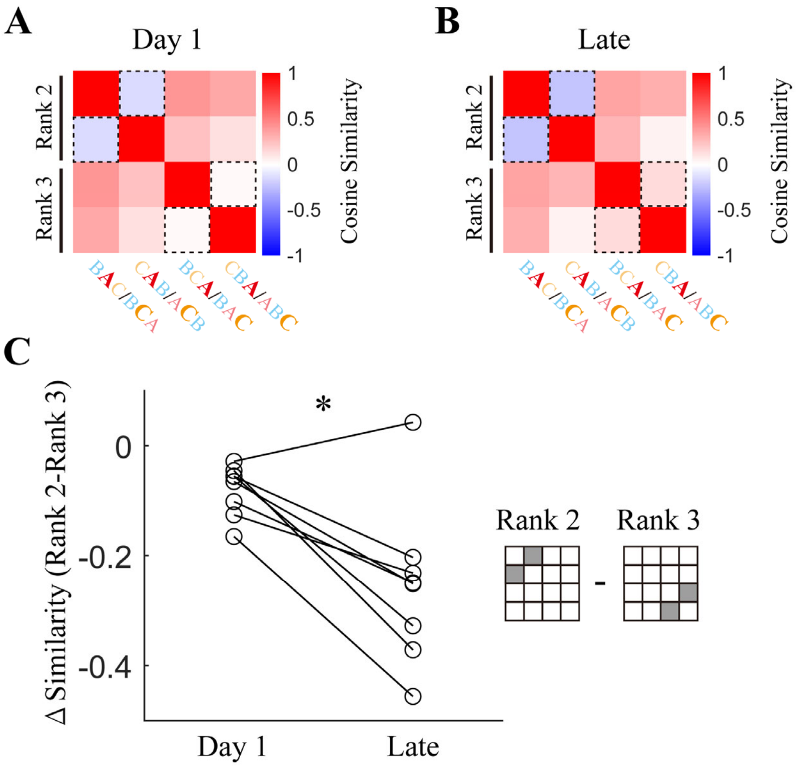


**Supplementary Figure 17. Representation similarity analysis of hippocampal representations at rank-2 and rank-3 with different past trajectories. (A)-(B)** Representation similarity across different conditions on day 1 (A) and late sessions (B). To remove impact from cue-selective response, neural activities in the two trials with the same reward order at each rank were first averaged (for example, BCA and BAC were averaged, to represent 'rank-2 as the first reward'). Cells were pooled from all mice in the reward order-encoding group (n=8). **(C)** The difference in similarity between rank 2 and rank 3 increased from day 1 to late session. $P$ = 0.0156; two-sided Wilcoxon signed-rank test; n=8 mice.

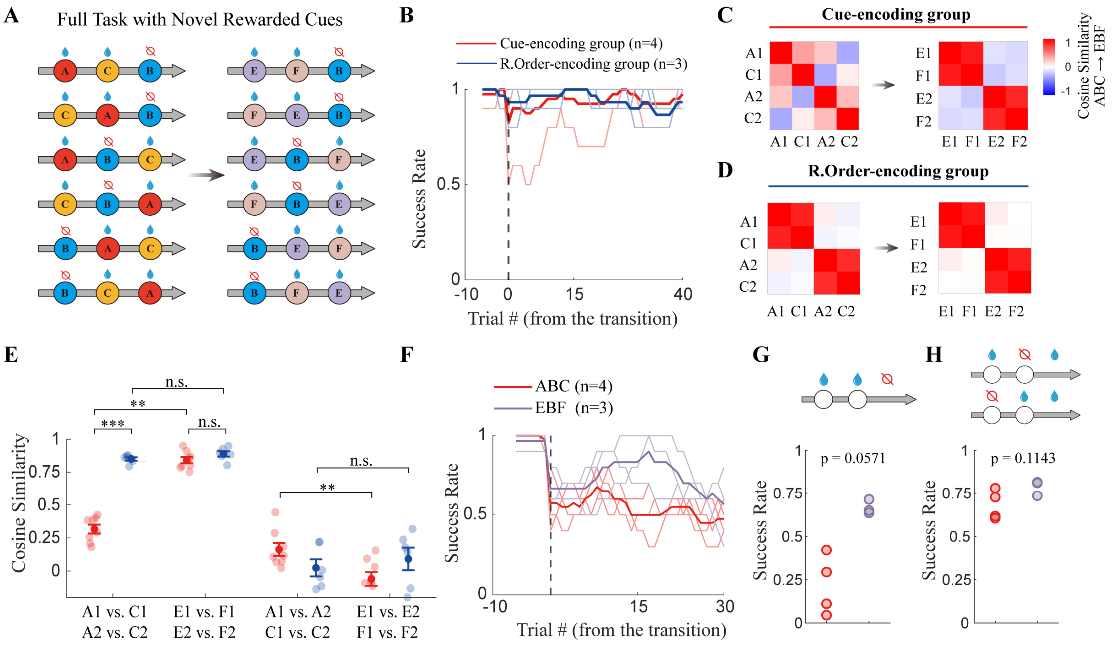

**Supplementary Figure 18. Learning a second problem with novel rewarded cues promoted abstract hippocampal representations and improved reward sequence completion. (A)** Schematic of E/B/F task. **(B)** Success rates following replacement of two novel rewarded cues in different groups of mice. Success rates were calculated using a sliding window of 10 trials. Dark curves indicate the mean, and light curves indicate individual mice; n = 3 mice for the reward order-encoding group and n = 4 mice for the cue-encoding group. **(C)** Cosine similarities of representations of different cues as different reward order under A-B-C and E-B-F conditions. Cells from cue-encoding group mice were pooled together. **(D)** Cosine similarities of representations of different cues as different reward order under A-B-C and E-B-F conditions. Cells from reward-order-encoding group mice were pooled together. **(E)** Quantification of representational similarities for individual mice. Light-colored dots indicate individual mice, and dark-colored dots with whiskers represent mean ± s.e.m. Red and blue denote the cue-encoding group and reward order-encoding group, respectively. $P$ = 6.660 × $10^{-4}$ for A1-C1/A2-C2, cue encoding vs. reward-order encoding group; $P$ = 0.1812 for E1-F1/E2-F2, cue encoding vs. reward-order encoding group; two-sided Wilcoxon rank-sum test; $P$ = 0.078 for cue-encoding groups, A1-C1/A2-C2 vs. E1-F1/E2-F2 and A1-A2/C1-C2 vs. E1-E2/F1-F2; two-sided Wilcoxon signed-rank test. **(F)** Success rates following the third-cue masking task under the E-B-F condition and the corresponding A-B-C condition in the same animals. One mouse was excluded from the E-B-F condition because of poor imaging quality and therefore did not undergo the third-cue masking task under this condition. **(G)** Success rates under conditions in which the last cue was invisible and non-rewarded. Two-sided Wilcoxon rank sum test. **(H)** Success rates under conditions in which the last cue was invisible and rewarded. Two-sided Wilcoxon rank sum test.

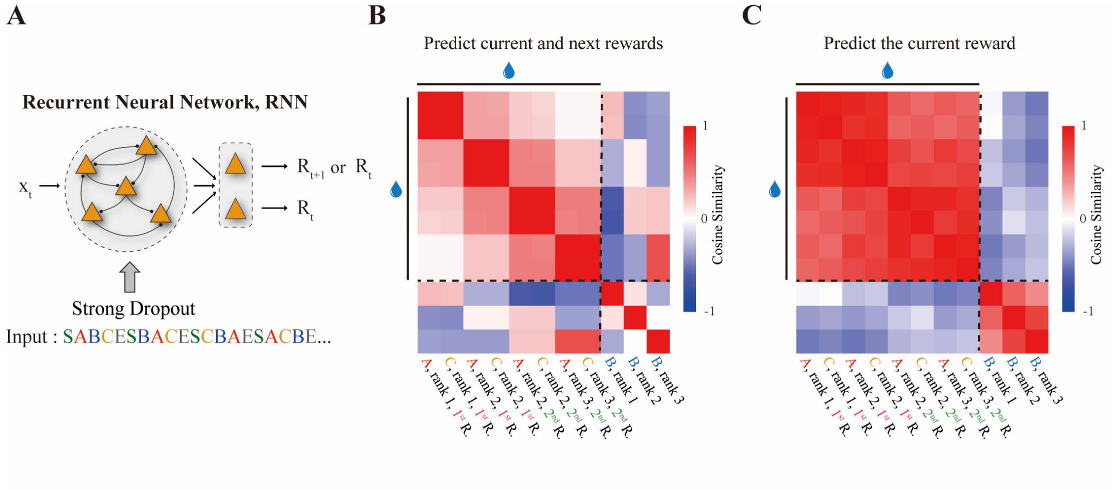


**Supplementary Figure 19. Representational similarity matrix of RNN hidden states. (A)** Schematic of RNN task with and without next-state prediction. To keep the network structure comparable, RNN with next-state prediction generated both $R_{t+1}$ and $R_t$, whereas the RNN without next-state prediction generated $R_t$ twice (i.e., replace the prediction target $R_{t+1}$ by $R_t$ ). **(B)** RSM across different conditions calculated from hidden layer activities of RNN with next-state prediction. **(C)** RSM of RNN without next-state prediction.

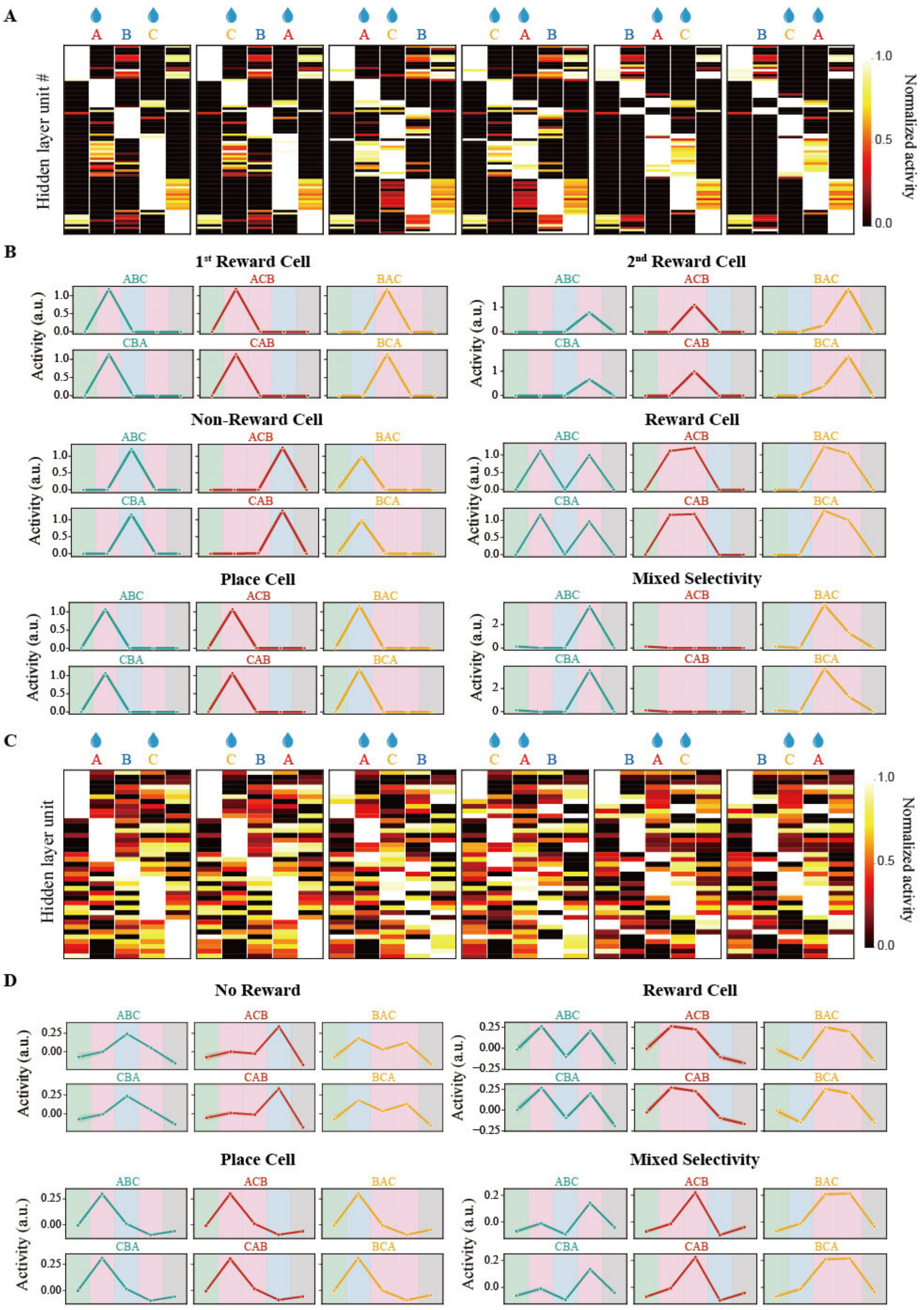


**Supplementary Figure 20. Single cell tuning in RNN and TEM. (A)** Hidden layer activities of RNN. In all heat maps, cells were sorted based on their activity peaks in the ABC trial type (the first column). **(B)** Example cells in RNN. Green, red, blue, and dark gray represent trial start, reward, non-reward, and trial end states, respectively. **(C)** Hidden layer activities of TEM. In all heat maps, cells were sorted based on their activity peaks in the ABC trial type (the first column). **(D)** Example cells in TEM.

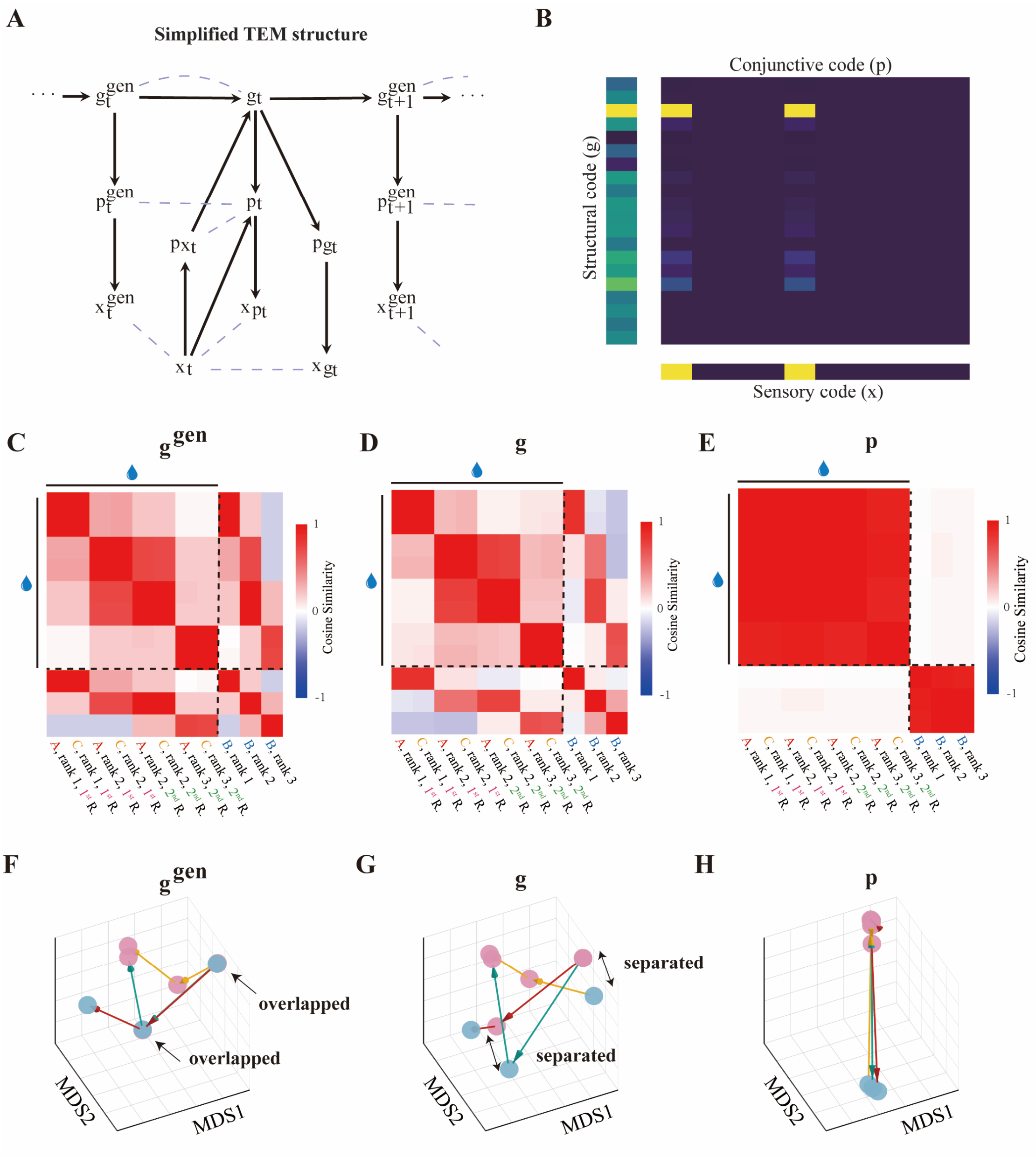


**Supplementary Figure 21. Representations of different components in TEM. (A)** Computation graph of the TEM. Solid arrows represent network forward propagation; dashed lines indicate the loss used to train the network. **(B)** Conjunctive code **p** was the clipped integration of structural code **g** and sensory code x within a single time step. Clipping diminished the structural information in **p** compared to **g**. Left, down-sampled structural code; bottom, two-hot sensory code; middle, reshaped conjunctive code. **(C)-(E)** RSM computed from $\mathbf{g}^{\mathbf{gen}}$, **g**, and **p** activities of the TEM. **(F)-(H)** MDS plots of $\mathbf{g}^{\mathbf{gen}}$, **g**, and **p** activities of TEM.

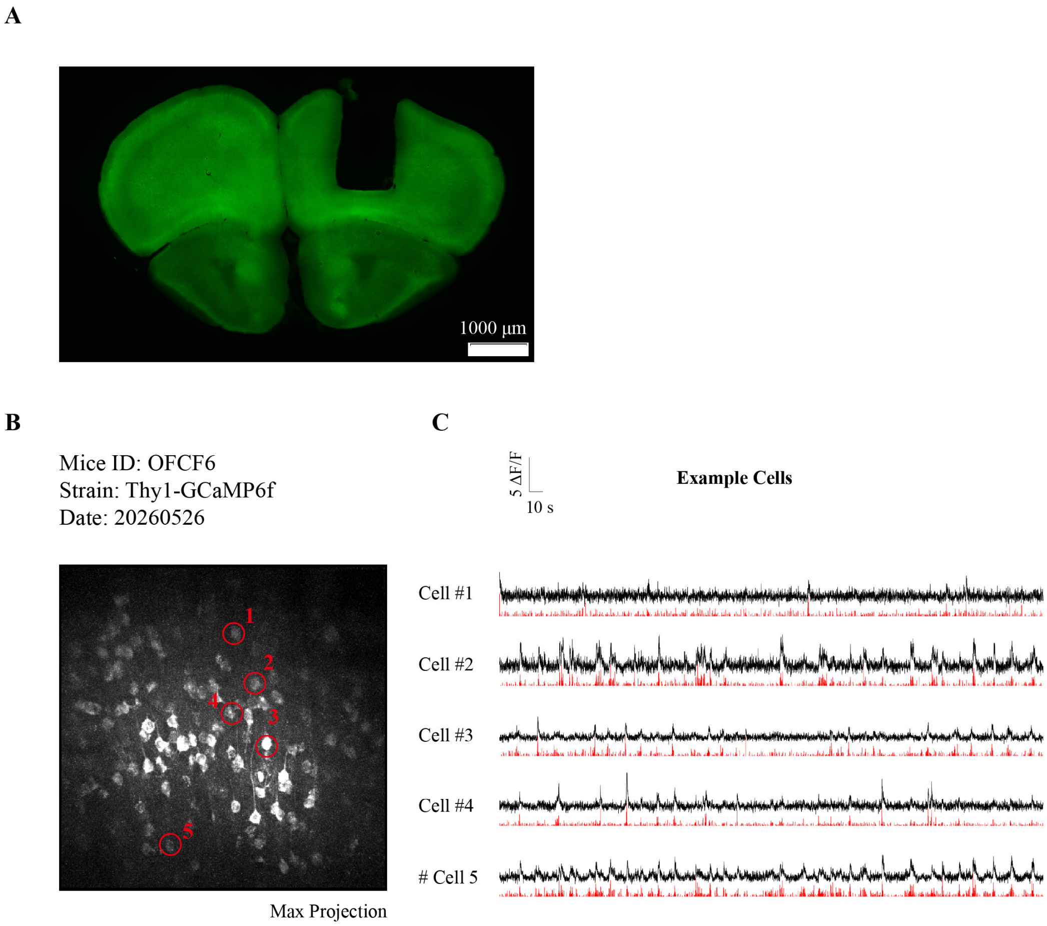


**Supplementary Figure 22. Two-photon calcium imaging in the OFC. (A)** Representative coronal section verifying the imaging location in the orbitofrontal cortex in a Thy1-GCaMP6f mouse. The black cavity indicates the implantation site of the grin lens. **(B)** Example two-photon imaging field from a Thy1-GCaMP6f mouse. Red circles indicate neurons corresponding to example activity traces shown in panel C. **(C)** Example activity traces from the neurons indicated in panel B. Black traces represent $\Delta F/F_0$ signals, and red traces represent putative ‘spiking’ activity from deconvolution.

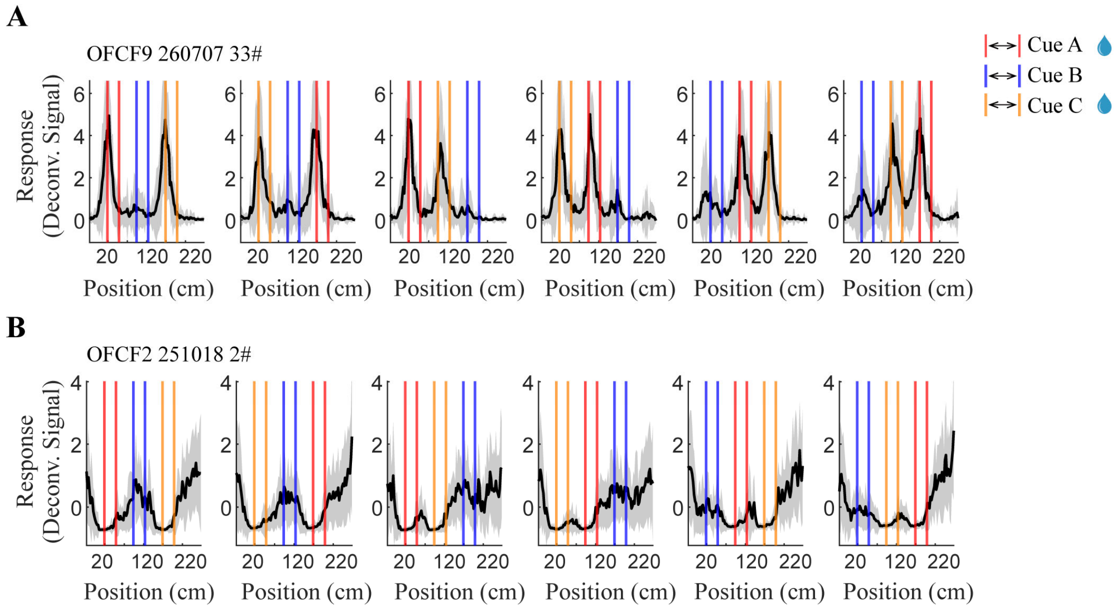


**Supplementary Figure 23. Reward-related activities of example OFC cells**. **(A)** Representative reward preferring neuron. Red, blue, and yellow lines indicate the positions at which cues A, B, and C were presented, respectively. Curves and shaded areas represent mean ± s.e.m., respectively. **(B)** Representative reward suppressed neuron. Red, blue, and yellow lines indicate the positions at which cues A, B, and C were presented, respectively. Curves and shaded areas represent mean ± s.e.m., respectively.
